# INSTINCT: Multi-sample integration of spatial chromatin accessibility sequencing data via stochastic domain translation

**DOI:** 10.1101/2024.05.26.595944

**Authors:** Yuyao Liu, Zhen Li, Xiaoyang Chen, Xuejian Cui, Zijing Gao, Rui Jiang

## Abstract

Recent advances in spatial epigenomic techniques have given rise to spatial assay for transposase-accessible chromatin using sequencing (spATAC-seq) data, enabling the characterization of epigenomic heterogeneity and spatial information simultaneously. Integrative analysis of multiple spATAC-seq samples, for which no method has been developed, allows for effective identification and elimination of unwanted non-biological factors within the data, enabling comprehensive exploration of tissue structures and providing a holistic epigenomic landscape, thereby facilitating the discovery of biological implications and the study of regulatory processes. In this article, we present INSTINCT, a method for multi-sample INtegration of Spatial chromaTIN accessibility sequencing data via stochastiC domain Translation. INSTINCT can efficiently handle the high dimensionality of spATAC-seq data and eliminate the complex noise and batch effects of samples from different conditions through a stochastic domain translation procedure. We demonstrate the superiority and robustness of INSTINCT in integrating spATAC-seq data across multiple simulated scenarios and real datasets. Additionally, we highlight the advantages of INSTINCT in spatial domain identification, visualization, spot-type annotation, and various downstream analyses, including motif enrichment analysis, expression enrichment analysis, and partitioned heritability analysis.

## Introduction

Chromatin accessibility is a measurement of the degree to which trans-acting factors can physically bind with cis-regulatory elements within open chromatin regions during DNA replication or transcription^1,2^. With the advances of single-cell epigenomic techniques^3-7^ in the recent years, abundant single-cell assay for transposase-accessible chromatin using sequencing (scATAC-seq) data have been generated, facilitating the study of heterogeneity among cells and contributing to the exploration of the mechanisms of biological regulatory networks^8^. However, the majority of complex regulatory processes rely on the intercellular transmission of ligand-receptor pairs within the tissue microenvironment, a dependence largely contingent upon the physical proximity of cells^9-11^, whereas scATAC-seq data offers only epigenetic profiles of individual cells without spatial information. This limitation impedes the attainment of a comprehensive understanding of tissues and organs through the analysis of scATAC-seq data.

With the innovation of spatial omics, emerging spatial epigenomic techniques, such as spatial ATAC^12^, spatial-ATAC-seq^13^, and SCA-seq^14^, along with some spatial multi-omics techniques, such as MISAR-seq^15^ and spatial ATAC-RNA-seq^16^, have given rise to spATAC-seq data. Although such data presently do not achieve single-cell resolution, wherein each barcoded spot on a sequencing array typically contains multiple cells, these techniques can provide coordinates for each spot based on its spatial location on the array. This capability enables researchers to integrate spatial information with epigenomic data, thereby facilitating the dissection of the structure and function of tissues and organs at various hierarchical levels. With the growing number of spATAC-seq datasets, the need for joint comparative analysis across multiple samples becomes increasingly urgent to fully leverage the information in each sample and gain a comprehensive understanding of the regulatory processes within the tissue microenvironment.

Integrative analysis has long been a challenging and meaningful topic for both single-cell omics^17-21^ and spatial omics^22,23^. Joint modeling of multiple samples can establish connections between different samples and enhance the statistical power for the analysis of shared biological variations, making samples comparable by effectively identifying and eliminating the unwanted non-biological factors within the data. Furthermore, joint modeling enables more detailed analyses and comparisons of temporal dynamics and spatial structures^24^, shedding light on the fundamental processes of biology and disease^21^. By integrating multiple spATAC-seq samples, researchers can comprehensively characterize spatial tissue structures and obtain a holistic epigenomic landscape, thereby facilitating the discovery of biological implications and contributing to the study of regulatory logic.

Many computational methods have been developed for multi-batch integration of single-cell or spatially resolved transcriptomics (SRT) data, enabling the integration of data from different samples, tissues, donors, developmental stages and technologies^25-31^. SCALEX^25^ is a method based on the variational autoencoder (VAE)^32^ for modeling single-cell RNA-seq or ATAC-seq data. By providing batch information only to the decoder, the batch-free encoder is enabled to merely extract biological information. PeakVI^26^ uses a Bernoulli distribution to model scATAC-seq data and employs a VAE combined with an auxiliary neural network to predict distribution parameters and perform batch correction. SEDR^27^ fuses spatial information with transcriptional profiles by leveraging a masked autoencoder and a variational graph convolutional autoencoder (GCN)^33^, while combining Harmony^20^ to eliminate batch effects. STAligner^28^ employs a graph attention (GAT)^34^ autoencoder for modeling SRT data, utilizing spot triplet loss for batch correction. STitch3D^31^ first pre-aligns multiple SRT samples to establish spatial connections between them, and achieves the removal of batch effects using a GAT-based batch-free encoder and a batch-specific decoder. However, no integration method specifically for spATAC-seq data has been developed.

Several fundamental challenges need to be addressed to accomplish the integrative analysis of spATAC-seq datasets. Firstly, spatial information should be incorporated into the analysis of spATAC-seq data effectively^9,35^. Methods for modeling scATAC-seq data mentioned above generally suffer from a common problem: they lack a design to utilize spatial information. Directly applying these methods to integrate spATAC-seq data may result in wasting the valuable information of local microenvironment, leading to unreliable biological analysis. For instance, after clustering, spots of the same cluster may be scattered in space, making it difficult to identify meaningful spatial domains with strong continuity. Secondly, spATAC-seq data generally exhibits high dimensionality, great sparsity, and suffers from a notable incidence of data missing due to a low capturing rate^3,36^,37. These characteristics result in different data patterns compared to SRT data, rendering some mechanisms for modeling SRT data unsuitable for analyzing spATAC-seq data, and therefore, may not yield satisfactory results. For example, STAligner^28^ identifies nearest neighbors across various samples using Euclidean distance and employs spot triplet learning to bring these identified neighbors, likely belonging to the same category, closer in latent space, thereby mitigating batch effects. However, we found that utilizing cosine similarity is more appropriate for spATAC-seq data, yielding better results compared to using Euclidean distance (Supplementary Fig. 1). Therefore, there is an urgent need for integration algorithms specifically designed for spATAC-seq data. Thirdly, spATAC-seq data presents complex patterns of noise and batch effects, posing challenges in distinguishing biological variations from non-biological factors. A procedure that can effectively remove noise and batch effects while preserving biological variations needs to be introduced, enabling comparability across multiple spATAC-seq samples for joint analysis^38^. Fourthly, the absence of methods capable of annotating spATAC-seq data complicates the exploration of biological significance that relies on annotated domains.

To this end, we present INSTINCT, a method for multi-sample integration of spatial chromatin accessibility sequencing data via stochastic domain translation, to address these gaps. To the best of our knowledge, INSTINCT is the first method specifically designed for integrative analysis of multiple spATAC-seq samples. INSTINCT trains a variant of GAT autoencoder to effectively integrate spatial information and epigenetic profiles, implements a stochastic domain translation procedure^39-41^ to facilitate batch correction, and finally obtains low-dimensional representations of spots in a shared latent space where the batch effects are eliminated and biological variations are preserved. The representations of spots can then be utilized for spatial domain identification, visualization, spot-type annotation and various downstream analyses, including motif enrichment analysis, expression enrichment analysis, and partitioned heritability analysis. Additionally, based on the cross-sample annotation results, INSTINCT can reveal the underlying biological implications, illustrating its potential to guide and contribute to the study of biological traits and pathology.

## Results

### Overview of INSTINCT

INSTINCT is designed based on the premise that, due to noise and batch effects resulting from varying experimental conditions and techniques, spots from different samples may belong to different domains in the feature space, even when some spots share similar biological attributes. With this consideration, INSTINCT aims to properly integrate these domains by eliminating unwanted non-biological bias while preserving biological variations among spots from different samples, thereby facilitating the integrative analysis of multi-sample spATAC-seq data.

As shown in Fig. 1a, INSTINCT is a principal component analysis (PCA)^42^-based method which adopts a two-stage strategy. In stage 1, the model comprises a GAT encoder^34^, a multilayer perceptron (MLP) decoder^43^, and a discriminator. Taking the preprocessed data matrix of multiple samples (*X*) and the concatenated neighbor graph as input, the GAT encoder fuses epigenetic profiles and spatial information of spots, projecting the data from different slices into a shared latent space (*Z*). The MLP decoder reconstructs each spot back to the domain of its original slice. The discriminator is pretrained to simultaneously judge the authenticity of the input data (*X*) and the reconstructed data (*X*_*rec*_), as well as the category of spots. In stage 2, the model further incorporates a noise generator module besides the GAT encoder, MLP decoder, and discriminator that are already trained in stage 1. A stochastic domain translation procedure^39-41^ is implemented to facilitate batch correction (Fig. 1b), wherein different patterns of noise and batch effects are artificially added to the low-dimensional representation of each spot, enabling flexible projection to the domains of different samples (Methods).

**Fig. 1:**
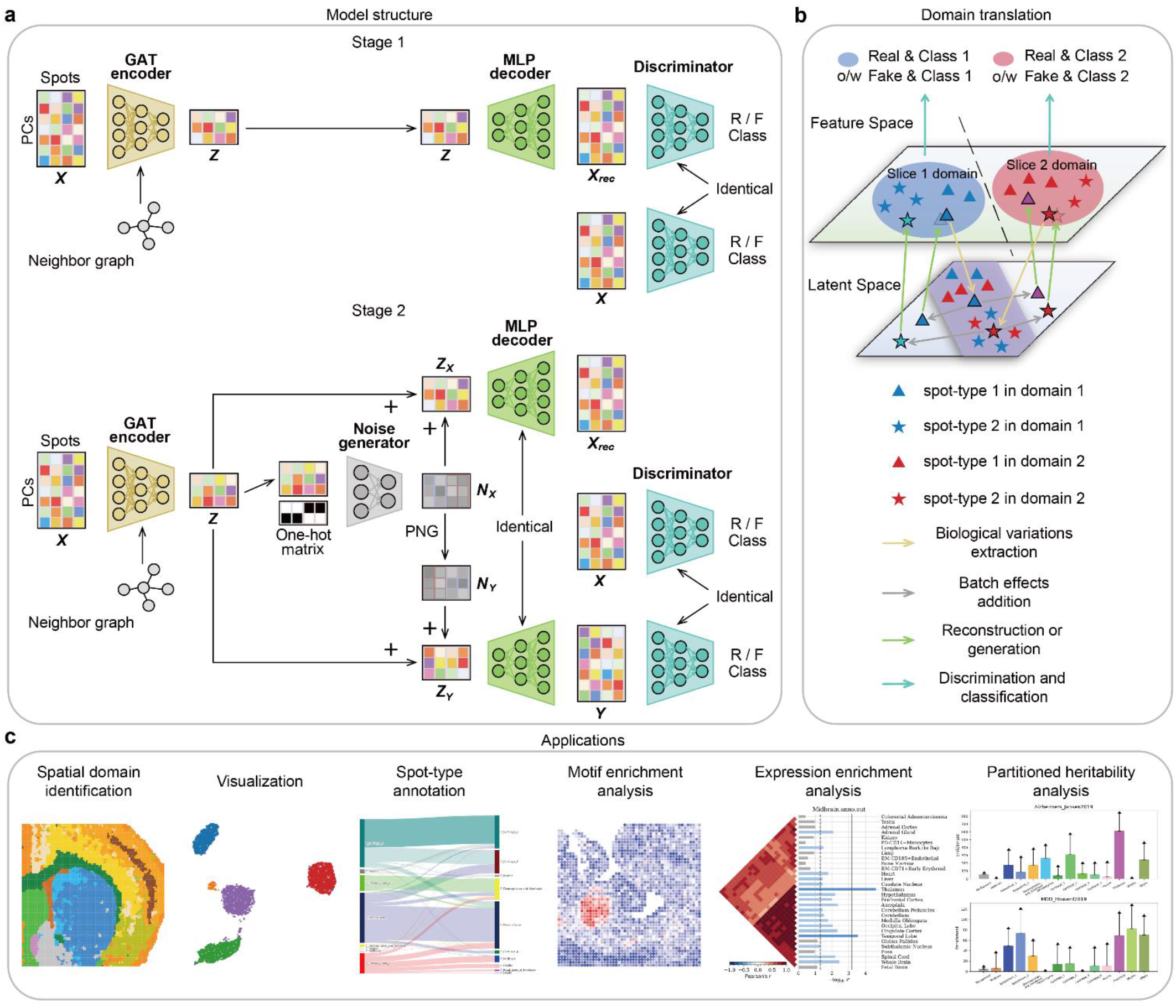
The overview of INSTINCT. **a**, the model structure of INSTINCT. INSTINCT adopts a two-stage strategy. In stage 1, the model of INSTINCT consists of a GAT encoder, a MLP decoder, and a discriminator. Taking the preprocessed data matrix of multiple samples and the concatenated neighbor graph as input, it simply learns to project all the spots from feature space (PCs space) into a low-dimensional shared latent space, while the discriminator is pretrained. During stage 2, a noise generator is added to the model. After the extraction of biological variations performed by the encoder, the noise generator captures the original pattern of batch effects for each spot in the latent space, and a PNG process simulates a different pattern of batch effects for each spot. These two simulated batch effects are then added to the corresponding spot separately, enabling reconstruction back to its origin domain or conversion to a new domain, referred to as the stochastic domain translation process. After training, the latent representation matrix *Z* can be extracted and utilized for various applications and downstream analyses. **b**, the sketch figure of the stochastic domain translation procedure, using the integration process of two samples as an illustration. The GAT encoder, MLP decoder, and discriminator respectively complete the processes of biological variations extraction, reconstruction or generation, as well as discrimination and classification. The batch effects addition procedure is completed by the noise generator and the PNG process. Specifically, the encoder extracts the biological variations within the data and projects it to the latent space. Subsequently, two different patterns of batch effects are added to the low-dimensional representation of each spot, enabling reconstruction to its original domain or generation to the target domain through the decoder. The raw data and the generated data are then discriminated and classified by the discriminator. **c**, the applications of the integration results of INSTINCT.

Specifically, during each training epoch in stage 2, each slice is assigned at random a target domain that is different from its original one, and spots from the slice need to be mapped to this target domain. After the extraction of biological variations performed by the encoder, the noise generator captures the original pattern of batch effects of each spot in the latent space using the latent representation matrix and a one-hot matrix indicating the original slices of spots, and a pseudo-noise generation (PNG) process simulates a different pattern of batch effects for each spot based on the output of the noise generator and the target domain of that spot (Fig. 1a, lower). These two patterns of batch effects are then added separately to the low-dimensional representation of the corresponding spot, enabling either reconstruction to the domain of its original slice (spot representation in *X*_*rec*_) or generation to the randomly selected target domain (spot representation in *Y*) through the decoder. The discriminator further performs the discrimination and classification process by simultaneously judging the authenticity of the input data and the generated data, as well as the category of spots, thereby ensuring a more accurate data generation and domain translation process through adversarial learning. This approach encourages the encoder to extract merely the biological variations and eliminate the batch effects effectively. It is worth noting that when mapping a spot to a target domain, our approach does not require the target domain to originally contain that specific spot-type. Instead, we implement a clamp strategy for the loss function to prevent domain-specific spot-types from being incorrectly mixed with others, thereby preserving sufficient biological variations (Methods).

After training, the latent representation matrix *Z*, which contains joint representations of spots without noise and batch effects, is used for spatial domain identification, visualization, spot-type annotation, and various downstream analyses such as motif enrichment analysis, expression enrichment analysis, and partitioned heritability analysis (Fig. 1c, Methods).

### INSTINCT shows superiority and robustness in comprehensive simulated scenarios

We first conducted comprehensive and realistic simulated experiments in various scenarios to validate the ability of INSTINCT in data integration.

Inspired by SpatialPCA^44^, we adopted a similar approach to simulate spATAC-seq datasets (Fig. 2a, Methods). Initially, we utilized simCAS^45^ to generate three scATAC-seq datasets that exhibited varying degrees of batch effects, each comprising 10,000 cells distributed among five cell-types labeled as integers 0 to 4. Each of these three datasets was used to generate one spATAC-seq sample. We pre-defined a single-cell resolution slice of size 120 × 150 as a template, which was divided into five domains represented by domain 0 to 4, respectively. Each domain corresponded to a major cell-type with the same number as its index (Fig. 2a), which was assigned to locations within that domain with a probability of *r*, and other cell-types were each assigned with a probability of 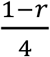. Subsequently, we defined each 3 × 3 cell grid as a spot to simulate the resolution of spATAC-seq techniques. The category of each spot was determined by the cell-type that predominantly occupied that spot. Evident batch effects between the three simulated slices can be observed in all scenarios (Supplementary Fig. 2).

**Fig. 2:**
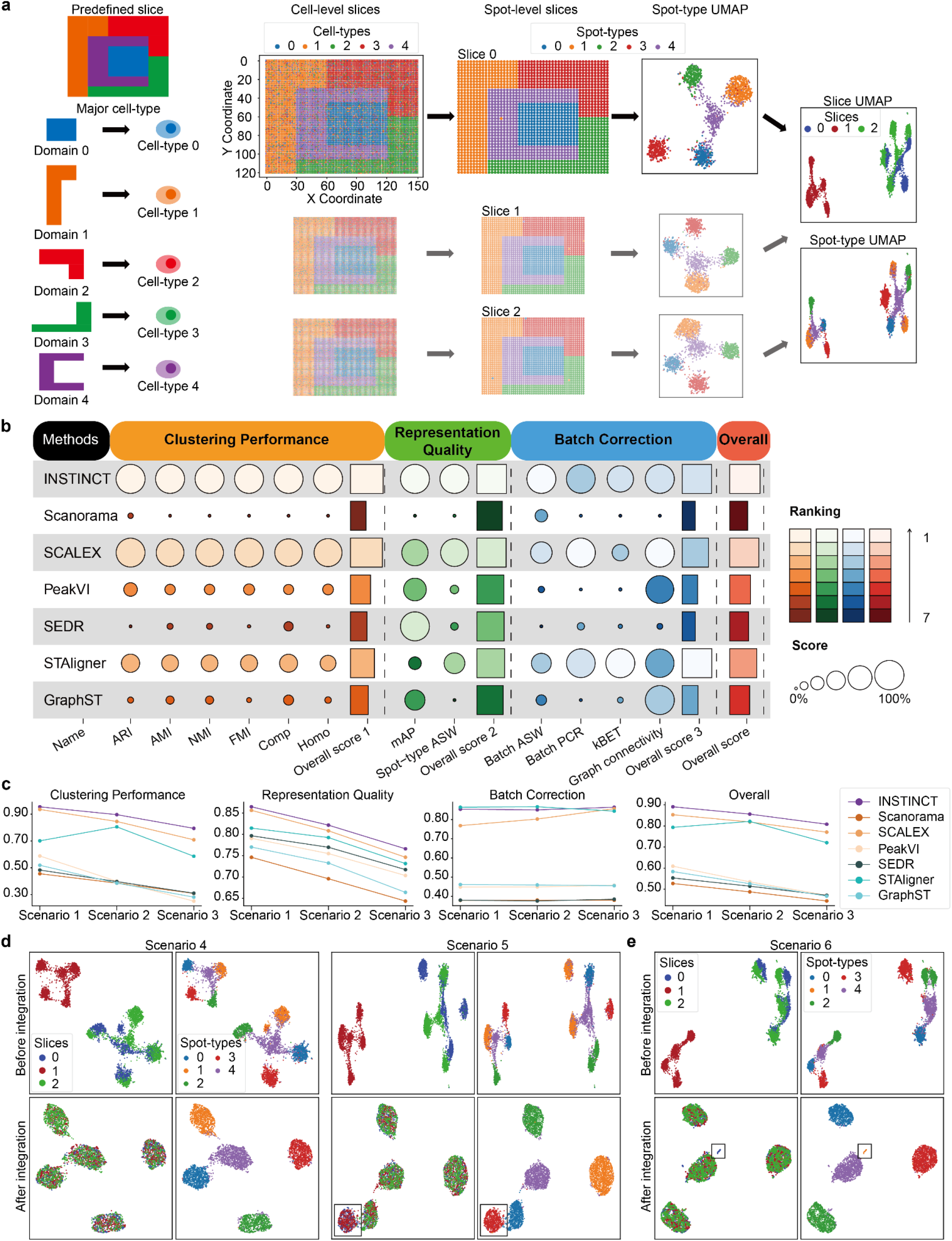
The simulation procedure and the performance of INSTINCT in simulated scenarios. **a**, the predefined template comprises five spatial domains, named by integers 0 to 4, each possesses a major cell-type within cell-type 0 to 4. Three simulated single-cell datasets with batch effects are generated by simCAS, each used to generate one spatial slice. Initially, each location in all slice is assigned with a cell-type, and a cell belongs to this cell-type in the corresponding single-cell dataset is randomly selected and allocated to the location to form three cell-level slices with single-cell resolution. Subsequently, with 3 × 3 region defined as a spot, the three slices are merged into spot-level slices, respectively. The category of each spot is determined by the cell-type that predominantly occupied that spot. Evident batch effects between simulated slices can be observed. **b**, the overview table of metrics calculated in scenario 1 demonstrates the superior performance of INSTINCT. INSTINCT reaches the highest score in 9 of the 12 metrics, and ranks second in 2 metrics. It also ranks first on the group-specific overall scores of clustering performance and representation quality, as well as the final overall score, and ranks second on the overall score of batch correction. **c**, the level of confusion among spot-types increases continuously and batch effects become more complex across the first three scenarios. INSTINCT consistently outperforms the benchmarking methods. **d**, two different transformations are applied to the data of scenario 1 to generate scenario 4 and 5, in order to simulate scenarios where the proportions of spot-types differ between slices. INSTINCT results in satisfying integration outcomes. **e**, based on scenario 1, scenario 6 is generated to test the ability of INSTINCT to preserve rare spot-types. INSTINCT succeed in preserving the rare spot-type which mainly exists in the first slice.

For scenario 1, the probability *r*, which is used to assign major cell-types to domains, is set to 0.8. We integrated the three slices and compared INSTINCT with several baseline integration methods for both single-cell omics and spatial omics (Fig. 2b, Methods). Integration methods for single-cell omics include Scanorama^21^, SCALEX^25^, and PeakVI^26^, with the latter two methods already demonstrated to perform exceptionally in integrating scATAC-seq data. Meanwhile, methods for spatial omics encompass SEDR^27^, STAligner^28^ and GraphST^29^, all designed for SRT data, as no integration method has been specifically developed for spATAC-seq data. We compared the capabilities of these methods in three aspects: clustering performance, representation quality, and batch correction, using a total of 12 metrics (Methods), with the first two aspect assessing biological conservation. INSTINCT achieved the highest score in 9 of the 12 metrics and second in two. It also ranked first on the group-specific overall scores of clustering performance and representation quality, as well as the final overall score, and ranked second on batch correction. Visualizations in Supplementary Fig. 3 also demonstrate the superior data integration capability of INSTINCT. Most of the baseline methods only integrated the two slices where the batch effects between them were not very pronounced (slices 0 and 2), but failed to mix the data from the other slice with them. However, INSTINCT evenly mixed the spots from the three samples, with spots of the same type gathering together while different types remain separated. While STAligner also integrated data from different batches, it mistakenly mixed different spot-types together, which resulted in significantly inferior clustering performance compared to INSTINCT.

We reduced the value of *r* continuously across the first three scenarios, resulting in increasing level of confusion among spot-types and progressively more complex batch effects (Supplementary Fig. 2, Methods). The same procedure was applied to evaluate the performance of all the methods in scenario 2 and 3. In the progressively challenging scenarios, INSTINCT consistently outperformed other methods (Fig. 2c, Supplementary Fig. 4). Specifically, in terms of the group-specific scores for clustering performance and representation quality, as well as the final overall score, INSTINCT consistently performed the best across all three scenarios, demonstrating its superiority and robustness. While in batch correction, it ranked second in the first two scenarios, with scores slightly lower than STAligner. As the scenarios became more challenging, INSTINCT surpassed STAligner in the third scenario, securing the first place. SCALEX also outperformed STAligner in the third scenario and came close to INSTINCT, demonstrating its relatively strong ability in batch correction. A detailed comparison of batch correction between INSTINCT and SCALEX is provided in Supplementary Text 1 and Supplementary Fig. 5. SCALEX and STAligner performed relatively well compared to other baseline methods. However, as the scenarios became more complex, and different types of spots became increasingly mixed before integration, the confusion in the low-dimensional representations obtained by SCALEX also increased. Consequently, distinguishing between different spot-types became more difficult. The same issue consistently occurred for STAligner across all the three scenarios. In contrast, INSTINCT not only evenly mixed data from three slices but also accurately distinguished between different spot-types in the latent space across scenarios 1 to 3 (Supplementary Figs. 3 and 6). This suggests that INSTINCT possesses the ability to not only effectively remove noise and batch effects but also accurately capture and highlight biological variations, contributing to subsequent analyses and downstream tasks. Additionally, we compared INSTINCT with two traditional PCA-based integration methods, Harmony^20^ and Seurat^46^, which ignore spatial information. INSTINCT demonstrated significantly better performance in both quantitative and qualitative analyses (Supplementary Text 2, Supplementary Fig. 7). For example, in visualizations of integration results, while the cluster for spot-type 0 identified by Harmony and Seurat contained many misclassified spots from other types, the cluster identified by INSTINCT contained nearly no other spot-types, highlighting the importance of utilizing spatial information.

In certain situations, vertically adjacent slices may not exhibit complete overlap, where different slices display varying proportions of spot-types, or a slice may lack some spot-types found in other slices. This issue can also arise in samples from different developmental stages. Therefore, based on scenario 1, we designed scenarios 4 and 5 to simulate two cases where the proportions of spot-types differ between slices (Methods). In scenario 4, we applied different cropping to the three slices from scenario 1 to generate three new slices (Supplementary Fig. 8). For slice 0, the proportion of spot-type 1 noticeably decreased, while for slice 2, the proportions of spot-types 2 and 3 decreased significantly. Despite the clear differences in the number and proportion of certain spot-types across different slices, INSTINCT still yielded excellent results when integrating the three slices in this scenario (Fig. 2d). In scenario 5, we retained slice 0 and slice 1 from scenario 1 and removed one domain from slice 2 (Supplementary Fig. 8). We applied INSTINCT to scenario 5 and observed that, although there was no spot identical to spot-type 3 in slice 2, INSTINCT did not incorrectly mix different spot-types in slice 2 together with spot-type 3 in other slices, as marked in Fig. 2d. These experiments demonstrate the flexibility and reliability of INSTINCT in various situations.

We further simulated scenario 6 based on scenario 1 to validate the ability of INSTINCT to preserve rare and specific spot-types (Methods). We excluded a certain domain from slices 1 and 2 of scenario 1, retaining only a 5 × 5 area of that domain in slice 0 and removing the rest, resulting in a rare spot-type with merely about 25 spots (Supplementary Fig. 8). After data integration performed by INSTINCT, the rare spot-type was perfectly preserved in latent space, as marked in Fig. 2e, indicating that INSTINCT can effectively identify unique biological signals in samples and prevent the over-mixing of different spot-types. To further explore the ability of INSTINCT to distinguish rare spot-types, we designed simulated scenario 7 based on a new template and three new simulated scATAC-seq datasets, consisting eight spot-types, with spot-types 5, 6, and 7 being rare (Methods). INSTINCT outperformed baseline methods in both quantitative and qualitative analysis. Specifically, through visualization, we found that INSTINCT not only effectively eliminated batch effects across the three slices but also perfectly distinguished the three rare spot-types, while none of the baseline methods could simultaneously reasonably mix the three samples and preserve all rare spot-types (Supplementary Text 3, Supplementary Fig. 9).

### INSTINCT integrates multiple samples across different developmental stages

We applied INSTINCT to a mouse brain dataset^15^ generated by MISAR-seq (MISAR-seq MB). The dataset comprises eight spATAC-seq data slices corresponding to developmental stages E11.0, E13.5, E15.5, and E18.5, with two vertically adjacent slices from each stage. We applied INSTINCT and baseline methods to integrate the four section 1 (S1) slices with annotations from each stage and evaluated the integration performance using various metrics. Since certain spot-types appear merely in specific slices, we included two additional metrics, isolated label ASW and isolated label F1-score^47^, to the metric group of representation quality to evaluate the preservation of slice-specific spot-types (Methods). Metrics that required spot-type information were calculated based on the provided annotations (Fig. 3e). INSTINCT ranked first in 10 out of 14 metrics and second in one. In terms of overall scores, INSTINCT secured the top position in clustering performance, representation quality, as well as the final overall score, and came second in batch correction (Fig. 3a, b).

**Fig. 3:**
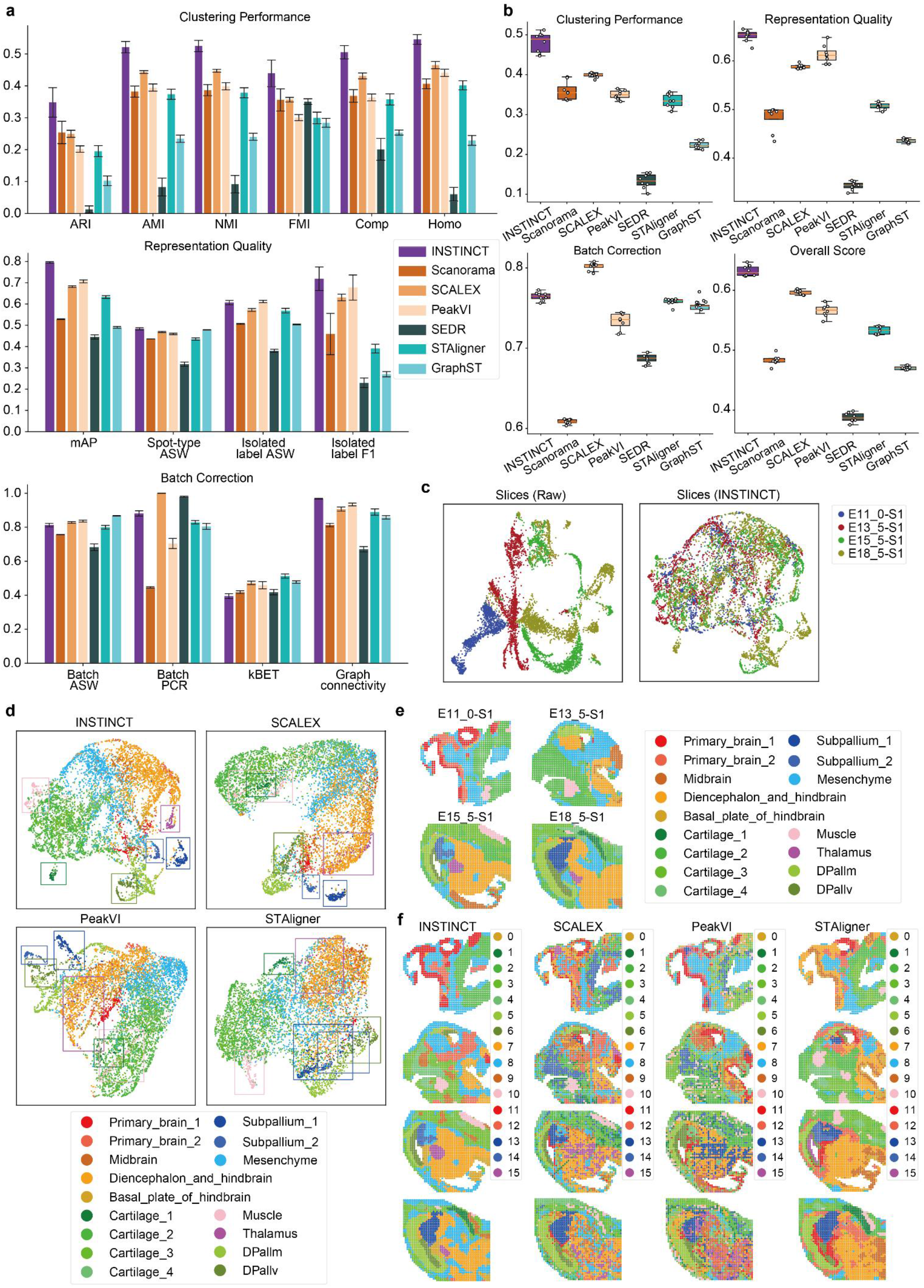
The comparison of evaluation metrics, visualizations and clustering results on the MISAR-seq MB dataset. **a**, the individual scores of each evaluation metric for INSTINCT and benchmarking methods. Two new metrics assessing the preservation of sample-specific spot-types, isolated label ASW and isolated label F1-score, are added to the group of representation quality. INSTINCT ranks first in 10 out of 14 evaluation metrics and second in one. **b**, the group-specific overall scores and the final overall score. INSTINCT ranks first in clustering performance, representation quality, as well as the final overall score, and second in batch correction. **c**, UMAP visualization of data from different developmental stages before integration and after integration performed by INSTINCT. **d**, UMAP visualization of integration results, with spots colored according to the provided annotations. Results of the four methods that rank in the top for the final overall score are displayed. **e**, the provided annotations of different slices. **f**, the clustering results of the four methods that rank in the top for the final overall score, with identified clusters first mapped to true labels and then colored accordingly.

We visualized the data before and after integration (Methods). Prior to integration, noticeable batch effects were observed among the four slices. However, after integrating with INSTINCT, the data from different slices were effectively mixed, removing the batch effects between them (Fig. 3c, Supplementary Fig. 10a). Through visualizations (Fig. 3d, Supplementary Fig. 10c), we found that INSTINCT not only accurately distinguished major categories such as cartilage, mesenchyme, and cerebral regions (e.g., diencephalon and hindbrain), but also preserved many rare spot-types, including thalamus, ventricular zone of the dorsal pallium (DPallv), a subtype of cartilage (cartilage_1), the two subtypes of subpallium (subpallium_1 and subpallium_2), and muscle. Each of these rare categories was highlighted with a box in the same color as the corresponding spot-type in Fig. 3d. Besides, most of these rare spot-types only concentrated in one or two developmental stages. For instance, subpallium_1 merely exists in slice of E18.5, while thalamus and subpallium_2 only exist in slices of E15.5 and E18.5. INSTINCT ensured that these types of spots retained their unique biological characteristics in the low-dimensional representation, enabling them to be distinguishable from other spot-types after integration. In contrast, most of the baseline methods could barely differentiate several major categories with significant biological differences and erroneously mixed many rare spot-types into these major categories (Fig. 3d, Supplementary Fig. 10b, c), resulting in the failure to capture and preserve the biological variations between different spot-types. For instance, SCALEX and PeakVI only distinguished subpallium_1 and subpallium_2 from other spot-types, mixing cartilage_1 and muscle with the other three cartilage subtypes (cartilage_2, cartilage_3, and cartilage_4). Besides, the thalamus could not be separated from the diencephalon and hindbrain. Although PeakVI can distinguish part of DPallv, the remaining part of DPallv was merged with the mantle zone of the dorsal pallium (DPallm). While STAligner was able to distinguish the muscle category, it failed to identify other rare spot-types. Moreover, the remaining three methods were unable to preserve any rare spot-types. It is worth noting that none of the methods distinguished thalamus, DPallv, and cartilage_1 like INSTINCT did. This demonstrates that INSTINCT can not only effectively remove batch effects but also preserve sufficient biological variations, thereby obtaining meaningful results in practical applications.

Subsequently, we jointly clustered the low-dimensional representations for each spot learnt by INSTINCT and mapped the identified clusters to the annotated categories (Fig. 3f, Methods). INSTINCT accurately identified numerous spatial domains across slices that hold practical significance. For instance, in the E11.0 slice, INSTINCT captured a relatively intact structure of the primary brain (cluster 11 and 12). Consistently across the other three slices, INSTINCT identified DPallm (cluster 5) and DPallv (cluster 6). Particularly, the structures of DPallm and DPallv identified in the E15.5 and E18.5 slices closely resembled the provided annotations. Cartilage_1 (cluster 1) and subpallium_1 (cluster 13) were accurately identified in the E18.5 slice. Subpallium_2 primarily exists in the E15.5 slice and only remains in a small portion at the lower end of the subpallium in the E18.5 slice. We found that this domain (cluster 14) was accurately identified not only in the E15.5 slice but also in the E18.5 slice. Thalamus (cluster 15), present only in the slices of the E15.5 and E18.5, was also accurately identified. Compared to other methods, INSTINCT also exhibited more accurate and comprehensive recognition of large domains, such as mesenchyme (cluster 8), cartilage (cluster 1 to 4), and diencephalon and hindbrain (cluster 7).

All baseline methods performed poorly in the spatial domain identification task (Fig. 3f, Supplementary Fig. 10d). While the three integration methods for scATAC-seq data were able to cluster most spots within certain large spatial domains into a single category, the clustering results were not spatially coherent. For example, SCALEX, while relatively accurately depicting the boundary of a large domain representing the diencephalon and hindbrain (cluster 7), exhibited scattered spots of multiple different clusters within it. Although PeakVI was able to identify the structure of the primary brain in E11.0 slice, it also incorrectly includes spots from other clusters within that structure. This not only rendered the identified spatial domains unreliable but also led to the failure to identify domains occupying smaller portions of the slices, such as thalamus, which none of the integration methods for single-cell omics recognized. This is because spATAC-seq data, representing the combined epigenetic profiles of multiple cells at each spot^16^, is more prone to confusion and less distinguishable between different categories compared to scATAC-seq data. Moreover, these methods did not consider the association of each spot with its surrounding environment by overlooking the spatial information provided by spATAC-seq data. While the three integration methods for SRT data could identify continuous spatial domains, their mechanisms were not suitable for spATAC-seq data. For instance, although STAligner could identify domains with continuity, the structure of some identified domains, such as caritilage_1 (cluster 1), significantly differed from the provided annotations and exhibited a larger volume (Fig. 3e, f). INSTINCT overcomes the limitations of both types of methods by implementing mechanisms suitable for spATAC-seq data and leveraging spatial information. The low-dimensional representations of spots can be utilized to yield superior results in spatial domain identification by accurately identifying continuous domains with boundaries consistent with those of real tissues.

### INSTINCT acquires joint representations of samples across different scales

We utilized INSTINCT to integrate three samples of mouse postnatal day 21/22 (P21/22) brains^16^ generated by spatial ATAC-RNA-seq (spatial ATAC-RNA-seq MB), a multi-omics sequencing technique. It is worth noting that although these three slices were sequenced from similar developmental stages of the same organ, the first two slices (slice 0 and 1) have a size of 50 × 50 barcodes, while the third slice (slice 2) contains 100 × 100 barcodes. This resulted in the sequencing of brain tissues of different scales, with slice 2 essentially encompassing the entire hemisphere of the coronal brain, while slice 0 and 1 only contained approximately one-quarter of its size.

We used INSTINCT to integrate these three samples with different tissue scales and performed clustering (Methods). The spots exhibited more obvious multi-group gathering patterns, with 13 clusters identified. Among these, cluster 11 and 12 were unique to slice 2, and cluster 9 was majorly found in slice 2 (Fig. 4b). We colored spots based on their clusters and observed that these clusters exhibited strong continuity in space (Fig. 4c). According to the anatomical reference of the coronal brain of adult mouse from Allen Brain Atlas website^48^ (Fig. 4a), we found that clustering restored the true tissue structure and identified spatial domains with practical significance. These include the caudate putamen (clusters 0 and 6), lateral ventricle (cluster 10), corpus callosum (cluster 3), lateral septal nucleus (cluster 4), and nucleus accumbens (cluster 9). Additionally, hierarchical structures were identified in the cortex (clusters 1, 2, 5, 8, and 11), and a small domain representing the island of Calleja (cluster 12), which only exists in slice 2, was also identified.

**Fig. 4:**
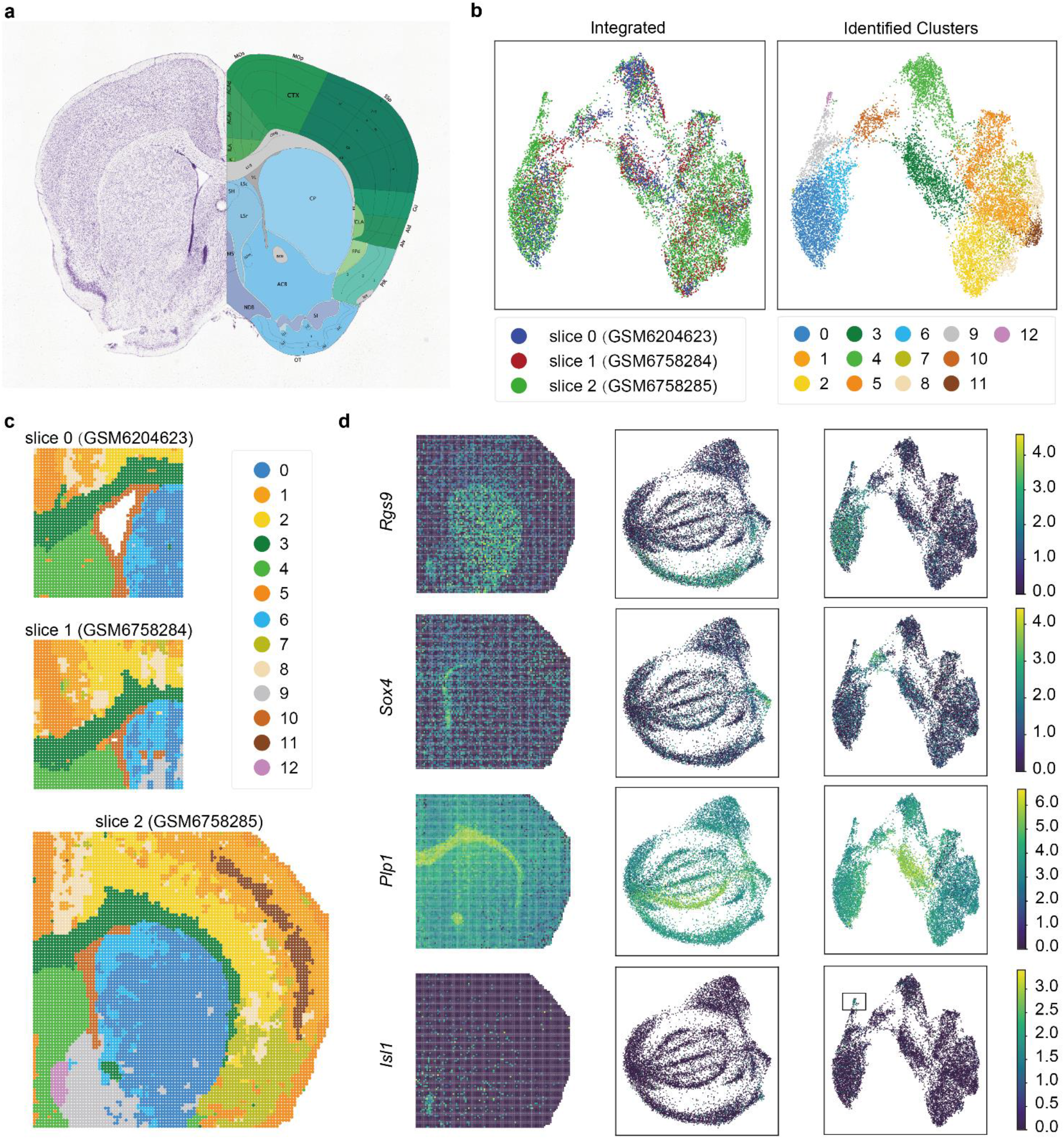
Integration results of INSTINCT on samples with different tissue scales from the spatial ATAC-RNA-seq MB dataset. **a**, Nissl (left) and anatomical annotations (right) from the Allen Mouse Brain Atlas and Allen Reference Atlas – Mouse Brain. Allen Mouse Brain Atlas, mouse.brain-map.org and atlas.brain-map.org. **b**, UMAP visualizations of the integration outcome and clustering results of INSTINCT. Biological variations of different spot-types are highlighted and 13 clusters are identified through clustering. **c**, spots are colored according to their specific clusters and displayed by their spatial arrangement. The clusters exhibit strong continuity in space, indicating the identification of spatial domains. **d**, visualizations of DEGs for identified clusters based on their expression levels.

Since spatial ATAC-RNA-seq involves multi-omics sequencing, the dataset also includes paired SRT data for these three slices, which provided the transcriptional landscapes. Using the SRT data corresponding to each spot, we identified the differentially expressed genes (DEGs) for each identified region (Methods) and visualized the expression levels of some marker genes (Fig. 4d, Supplementary Fig. 11). For example, in the caudate putamen, marker genes such as *Rgs9*^49^, *Pde10a*^50^, *Gng7*^51^, *Bcl11b*^52^, and *Foxp1*^53^ showed significantly higher expression levels. INSTINCT made the spots in this domain more clustered compared to before integration. *Sox4*^54^, *Dlx1*^55^, and *Zbtb20*^56^ were predominantly expressed in the lateral ventricle. Before integration, these spots were not clearly distinguished from others, but after integration, they were separated and aggregated in a small area. Cluster 3, representing the corpus callosum, showed high expression of genes such as *Plp1*^57^ and *Mbp*^58^. While spots of cluster 3 were mixed internally and difficult to distinguish from other categories before integration, INSTINCT made them separately gathered and identified as a distinct category through clustering. Marker genes for the island of Calleja, such as *Isl1*^59^ and *Rreb1*^60^, were highly expressed only in cluster 12 unique to slice 2, indicating that INSTINCT, when integrating samples with different tissue scales, can preserve non-shared rare spot-types. Additionally, the expression of *Dgkg*^61^ in the lateral septal nucleus and *Mef2c*^62^ in the cortex further demonstrated that INSTINCT better distinguishes between spot-types that were previously difficult to differentiate (Supplementary Fig. 11), proving its effectiveness not only in integrating samples with different tissue scales but also in highlighting the biological variations between different spot-types.

### INSTINCT facilitates the integrated analysis of whole mouse organisms

We applied INSTINCT to a mouse embryo dataset generated by spatial-ATAC-seq (spatial-ATAC-seq ME)^13^, which contains one sample from E11 developmental stage and two samples from E13 stage, with the sequencing depth of the ME13_1 slice being significantly higher than that of the other two slices, resulting in over 590,000 peaks and significantly higher fragment counts (Fig. 5a). The other two slices have only about 290,000 peaks, but higher ratio of reads in transcription start sites (TSS) compared to the ME13_1 slice. Consequently, the batch effect between ME13_1 and the other two slices was pronounced (Fig. 5b).

**Fig. 5:**
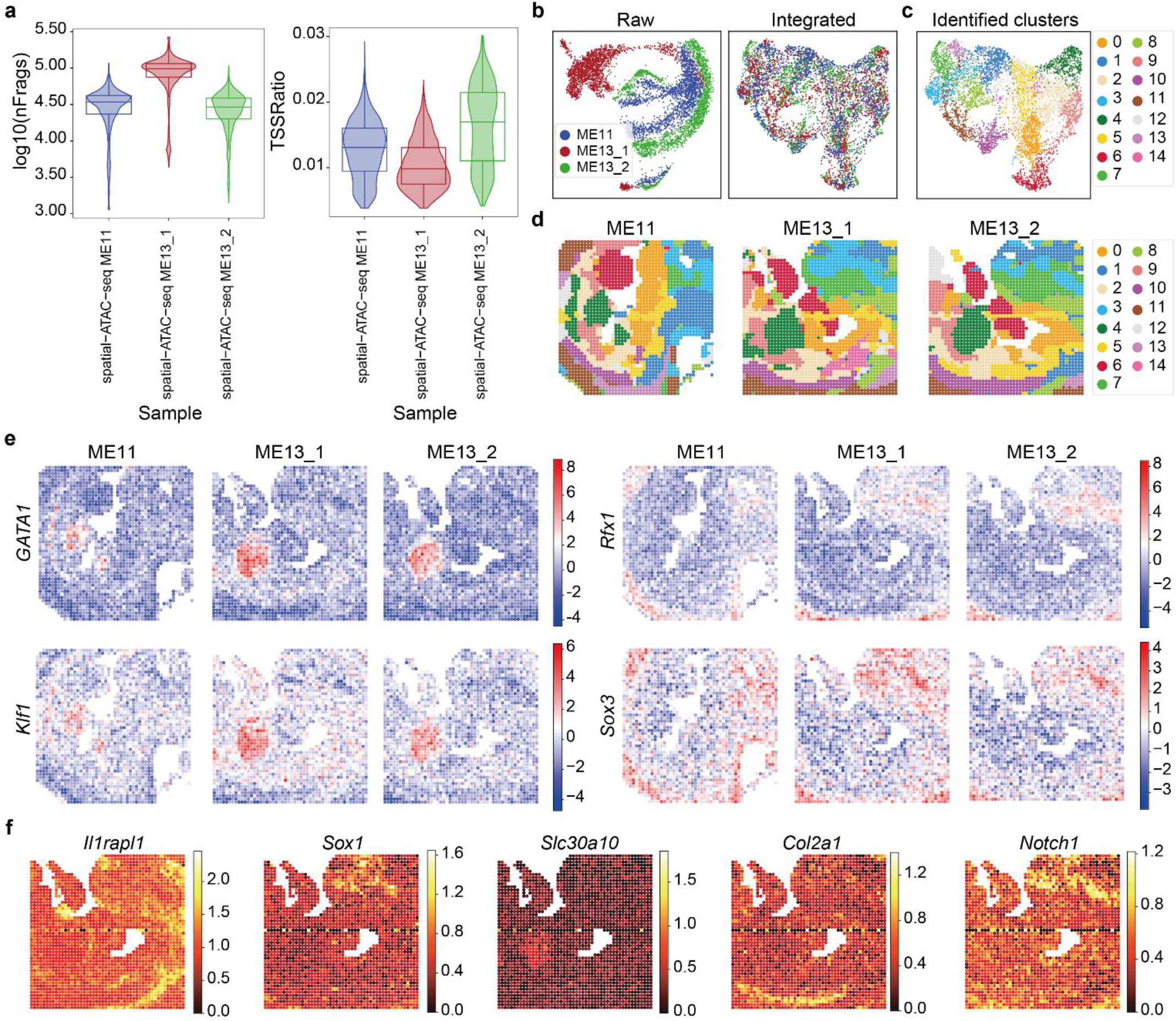
Integration results of INSTINCT on samples with varying sequencing depths from the spatial-ATAC-seq ME dataset. **a**, comparison of the log-normalized fragment counts and the TSS ratios across the three samples from the spatial-ATAC-seq ME dataset. **b**, UMAP visualizations of the raw data and the integrated results of INSTINCT, with spots colored by slice affiliation. **c**, UMAP visualization of the clustering results of INSTINCT, with spots colored based on identified clusters. **d**, spatial arrangement of clusters, with spots colored according to their specific clusters. **e**, motif enrichment analysis results for the three slices, with spots colored based on the motif enrichment score with respect to specific motifs. **f**, spatial arrangement of DEGs for identified clusters based on their expression levels.

We used INSTINCT to integrate these three slices and successfully mixed the spots from them after integration. Joint clustering identified 15 clusters, and the spatial arrangement of these clusters showed obvious similarities across the three slices (Fig. 5c, d). According to the anatomical annotation of major tissue regions provided by the original study^13^, we were able to jointly identify biological spatial domains across the three slices based on the clustering results, including the midbrain (cluster 1), forebrain and ventricles (cluster 3), liver (cluster 4), limb (cluster 6), spine (cluster 10), and spinal cord (cluster 11).

We then performed motif enrichment analysis following previous researches^63-66^ (Methods). After identifying the cluster-specific peaks for each cluster, we combined them into a peak set and used the data corresponding to these peaks as the filtered data, which was then used to infer the enriched transcription factor (TF) binding motifs within these peaks. We found that the clustering results after integration with INSTINCT could help reveal cluster-specific motifs (Fig. 5e, Supplementary Fig. 12). For example, we found that the binding motifs of the key hematopoietic transcription factors *GATA1*^67^ and *GATA4*^68^, which are highly related to liver development, were significantly enriched in cluster 4, which indicates the liver. The binding motif of *Klf1*, which has been validated to be expressed in the central macrophages of erythroblastic islands (CMEIs) in murine fetal liver and plays an important role in the digestion of the DNA of extruded red blood cell nuclei during maturation^69^, was also found to be enriched in cluster 4. Moreover, previous studies have demonstrated the importance of the RFX family^70^ and SOX family^71,72^ in central nervous system (CNS) development, and through motif enrichment analysis, we found that the binding motifs of *Rfx1, Sox2*, and *Sox3* were enriched in cluster 3, representing the forebrain and ventricles, and in cluster 11, representing the spinal cord, confirming this point.

Next, we applied ArchR^73^ to generate the gene score matrix for the ME13_1 slice, which has the highest sequencing depth, and identified DEGs for all clusters (Methods). The identified DEGs showed high agreement with the recognized spatial domains (Fig. 5f, Supplementary Fig. 13). For instance, *Il1rapl1*, which plays a crucial role in cerebellar development by establishing local excitation/inhibition balance^74^, as well as *Cntn4* and *Ptprd*, which are involved in neurodevelopment^75,76^, were highly expressed in brain-related cluster 8. *Sox1* and *C130071C03Rik* were predominantly expressed in cluster 3, which represents the forebrain and ventricles, consistent with previous knowledge^77-79^. The manganese efflux transporter *Slc30a10*, required for manganese excretion by the liver^80,81^, was primarily expressed in cluster 4. Cluster 10, which represents the spine, was enriched with genes essential for skeletal development, such as *Fgfr2*^82,83^, *Col2a1*^84^, and *Mir140*^85^. Notably, *Notch1* was highly expressed in both cluster 3 and cluster 11, which represents the spinal cord, highlighting its significant role in mammalian CNS development^86,87^.

The DEGs identified for clusters were also used to perform gene ontology (GO) analysis (Methods), yielding reasonable results (Supplementary Fig. 13). According to the analysis, the GABAergic synaptic transmission process was enriched in cluster 1, which represents the midbrain, consistent with previous study^88^. In the CNS-related clusters, cluster 3 and cluster 11, the processes of neuron generation and differentiation were enriched. Additionally, since Wnt family is known to control various processes during limb development^89,90^, the canonical Wnt signaling pathway was expectedly enriched in cluster 6, which represents the limb.

We further applied INSTINCT to another mouse embryo dataset^12^ generated by spatial ATAC (spatial ATAC ME), which consists of six annotated slices from developmental stages E12.5, E13.5, and E15.5, with two vertically adjacent slices from each stage. We conducted both quantitative and qualitative analyses to compare INSTINCT with baseline methods on this dataset (Supplementary Text 4, Supplementary Figs. 14 to 23). INSTINCT ranked first in clustering performance, representation quality, and the final overall score. It is important to note that due to the low sequencing depth and data quality of this dataset, most baseline methods over-mixed different spot-types. However, INSTINCT did not excessively mix all slices and accurately categorized the spots into four major groups: CNS, meningeal and peripheral nervous system (PNS), mesenchyme and limb, and liver, consistent with the results reported in the original study^12^. Besides, INSTINCT performed much better in spatial domain identification task. For instance, it accurately identified the liver (cluster 5) in slices of E12.5 and E13.5. In the E15.5 slice, it identified spatial relationships within brain tissue, distinguishing between forebrain (cluster 1), midbrain (cluster 8), and hindbrain (cluster 2).

These experiments indicate that INSTINCT can integrate multiple samples of whole organisms across various scenarios and accurately capture underlying biological variations, demonstrating robustness while effectively avoiding over-mixing different spot-types.

### INSTINCT reveals biological implications through cross-sample annotation

As spATAC-seq data becomes increasingly abundant, the need for cross-sample annotation methods, which are crucial for understanding biological processes across different samples and revealing biological implications based on annotated domains, grows more urgent. However, there is currently no computational annotation algorithm specifically designed for spATAC-seq data that can use labeled spATAC-seq data as a reference to automatically annotate target unlabeled spATAC-seq datasets, without relying on manual modifications. INSTINCT addresses this gap by enabling cross-sample annotation, enhancing the analysis and interpretation of spatial epigenomic landscapes and facilitating the discovery of biological insights.

Initially, we used the integration results of various methods to perform cross-sample spot-type annotation on simulated datasets with predefined ground truth as quantitative validation (Methods). Specifically, we completed this by employing a k-nearest-neighbor (kNN) classifier, a model-free approach which acquire annotations based on the joint representations obtained by integration algorithms. We evaluated the annotation results using four metrics (Methods): accuracy, kappa, macro F1-score (mF1), and weighted F1-score (wF1). INSTINCT consistently ranked first on all four metrics across scenarios 1 to 3 (Supplementary Fig. 24a to c). According to the confusion matrices of the annotation results provided by INSTINCT for each slice across simulated scenarios 1 to 3, INSTINCT was able to annotate spots in each category (Supplementary Fig. 24d).

Rather than focusing on quantitative comparisons, we are more interested in whether INSTINCT can provide interpretable biological insights in the annotated spatial domains, thereby expanding its application to exploratory and discovery scenarios. Therefore, for the MISAR-seq MB dataset^15^, we used the S1 slices with provided annotations from four developmental stages to annotate the corresponding unannotated S2 slices, respectively. Before integration, batch effects existed between each S1 and its corresponding S2 slice (Supplementary Fig. 25a). Initially, we preprocessed the samples using the preprocessing procedure provided by INSTINCT, and then directly reduced the dimensionality of the data to match the latent representations of spots obtained by INSTINCT using PCA. We found that when annotating samples with complex structures, such as slices from E15.5 and E18.5, spots labeled as the same type were spatially discontinuous, resulting in inaccurate identification of tissue structure (Supplementary Fig. 25b). Moreover, due to batch effects, spots of a certain type in S1 slices were not mixed with those labeled as the same type in corresponding S2 slices, rendering the annotation results unreliable.

We then integrated each pair of S1 and S2 slices from each developmental stage separately using INSTINCT and performed annotation using the joint low-dimensional representations. We observed that after integration, batch effects between the two vertically adjacent slices were eliminated (Supplementary Fig. 26a), and the spatial structures of the annotations in S2 slices were consistent with those of the corresponding S1 slices (Supplementary Fig. 26b). For instance, in E18.5 slices which possessed the most complex structures, the relative spatial relationships between different spatial domains annotated in S2 were highly consistent with those in S1 (Fig. 6a). INSTINCT successfully annotated many important structures such as subpallium_1, DPallv, DPallm, thalamus, and their shapes and volumes were nearly identical to the corresponding regions in S1. Notably, in the S1 slice of E18.5, there were two types of spots with very low proportions, subpallium_2 and muscle. Similar proportions of these two types of spots were also annotated in similar spatial locations in the S2 slice. We conducted GO analysis (Fig. 6b) using SRT data paired with the S2 slice of E18.5, and found that for the annotated subpallium_2, there was enrichment in the biological process of forebrain neuron differentiation, consistent with the results obtained using four S1 slices in the original study^15^. For the category annotated as muscle, enrichment was observed in many biological processes related to muscle, including muscle contraction and skeletal muscle contraction, consistent with the enrichments observed in the original study^15^.

**Fig. 6:**
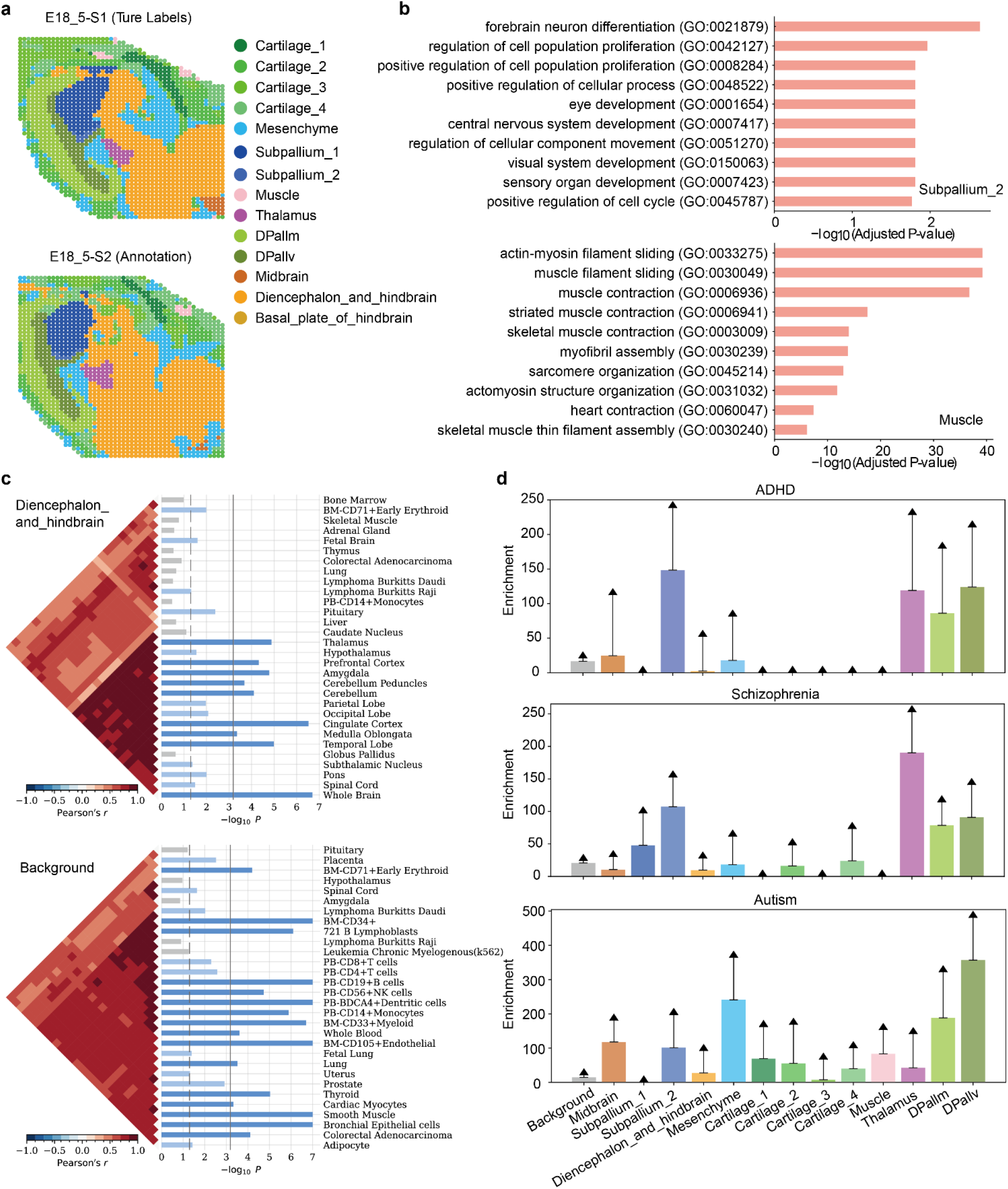
Annotation results and downstream analyses on the MISAR-seq MB dataset. **a**, the provided annotations of the S1 slice of developmental stage E18.5 and the annotation outcome of the corresponding S2 slice based on the integration results of INSTINCT. **b**, GO analysis performed for the two rare spot-types, subpallium_2 and muscle, identified in the S2 slice. **c**, results of expression enrichment analysis on the spot-type-specific peaks of diencephalon_and_hindbrain and background peaks. The top 30 significantly enriched tissues are displayed. **d**, the results of partitioned heritability analysis performed on the spot-type-specific peaks and background peaks for three diseases, including ADHD, schizophrenia, and autism.

We further identified spot-type-specific peaks for each annotated spot-type, using the remaining peaks as background peaks (Methods), and performed expression enrichment analysis for the single nucleotide polymorphisms (SNPs) in each set of spot-type-specific peaks and the set of background peaks, respectively (Fig. 6c, Supplementary Figs. 27 and 28, Methods). We found that genes in spot-type-specific peaks of all cerebral regions were predominantly enriched in brain-related tissues. Conversely, genes in the specific peaks of non-brain-specific tissues, such as cartilage and muscle, did not exhibit enrichment in brain-related tissues. Additionally, genes in the background peaks were generally not enriched in brain-related tissues. This further confirms that the annotation provided by INSTINCT can help distinguish spots from different biological systems with distinct functionalities.

Furthermore, we utilized the SNPs found in each set of spot-type-specific peaks and the set of background peaks to perform partitioned heritability analysis (Methods). We found that in many psychiatric disorders and mental-related phenotypes, the enrichment of heritability in cerebral regions was significantly higher than that in mesenchyme, cartilage, and muscle (Fig. 6d, Supplementary Figs. 29 and 30), and it tended to be biased towards certain specific regions. For instance, attention deficit hyperactivity disorder (ADHD) is associated with the pallium and thalamus^91^. Our result validated this association, as its heritability was significantly enriched in subpallium_2, DPallv, DPallm, and thalamus, while other regions exhibited relatively lower levels of enrichment. For schizophrenia, heritability was significantly enriched in the thalamus, highlighting its importance in the implicated circuitry, which has been validated in previous studies^92,93^. Autism has been linked to the dorsal pallium^94^, and the result indicated that the enrichment of heritability in DPallv was significantly higher than in DPallm, aiding in pinpointing a specific location within the tissue. This emphasizes that INSTINCT can reveal biological implications in the annotated domains, thereby guiding and contributing to the studies of biological traits and pathology.

### Ablation studies validate the effectiveness of the mechanisms implemented in INSTINCT

To demonstrate the effectiveness of the mechanisms adopted in INSTINCT, we conducted a series of ablation experiments on four annotated S1 slices of the MISAR-seq MB dataset^15^.

Firstly, we performed ablation experiments on the designed loss functions (Methods). It was observed that the exclusion of any loss function resulted in a degradation of the performance of the model (Supplementary Fig. 31), thereby substantiating the efficacy of all loss functions.

Additionally, since INSTINCT includes a noise generator and discriminator for achieving stochastic domain transformation (Methods), we conducted ablation experiments on the model structure. We compared the integration effectiveness of the complete INSTINCT model with the model where the discriminator was removed, as well as the model where both the discriminator and noise generator were removed. We did not consider the model where only the noise generator was removed due to the structural constraints of INSTINCT. We found that the complete model achieved the best performance, demonstrating the effectiveness of the discriminator and noise generator modules (Supplementary Fig. 32). Additionally, we observed that the performance of the model was better when both the discriminator and noise generator were removed compared to when only the discriminator was removed. This suggests that the discriminator plays a crucial role in constraining noise simulation and stochastic domain translation.

Moreover, since we employed a clamp strategy in INSTINCT to prevent the erroneous mixing of different types of spots while enhancing the conservation of spot-types unique to a particular slice (Methods), we conducted an ablation experiment for it. we found that adopting this strategy led to improvements in all three aspects including clustering performance, representation quality, and batch correction (Supplementary Fig. 33k, l). It is noteworthy that employing this strategy enhanced the performance of INSTINCT on isolated label ASW and isolated label F1-score, demonstrating the ability of this strategy to preserve sample-specific spot-types.

A more detailed analysis of the mechanisms implemented in INSTINCT and their effects on data integration can be found in Supplementary Text 5 and Supplementary Fig. 33.

## Discussion

Recent advances in spATAC-seq technologies have led to an increasing amount of spATAC-seq data, which simultaneously provides epigenomic landscapes and spatial information. However, existing integration methods for other omics are not well-suited for integrating this data modality. We propose INSTINCT as the first method capable of effectively integrating spATAC-seq data, demonstrating superiority across multiple tasks and providing a guidance for spATAC-seq data integration. INSTINCT enables the integrative analysis of multiple samples, facilitating the discovery of biological insights that might be missed when analyzing individual slices (Supplementary Text 6, Supplementary Figs. 34 and 35). Besides, INSTINCT can fully utilize spatial information, allowing the identification of tissue regions with stronger spatial continuity compared to integration methods developed for scATAC-seq data. Moreover, the stochastic domain translation step in INSTINCT enables the noise generator to effectively extract noise and batch effects from the low-dimensional representations, allowing the representations of spots to effectively highlight biological variations. We also demonstrated that the outstanding integration results of spATAC-seq data provided by INSTINCT can be utilized for cross-sample annotation and further reveal biological implications in the annotated domains. Furthermore, we found that INSTINCT can also integrate SRT samples, achieving performance comparable to established methods specifically designed for SRT data integration, and in some cases, even surpassing certain methods (Supplementary Text 7, Supplementary Fig. 36).

Nevertheless, INSTINCT still has some limitations. Firstly, INSTICNT does not directly model the data distribution as some methods do. For example, PeakVI models the count of each peak within each cell using a Bernoulli distribution and possesses the ability to correct for the peak accessibility signal at a single-region resolution, enabling it to accurately identify key cis-regulatory elements^26^. However, INSTINCT lacks this ability. Secondly, to obtain latent representations of spots from different datasets, INSTINCT requires to train a specific model for each integration task. In contrast, some methods, such as SCALEX, can integrate multiple batches of data in an online manner without the need for retraining the model^25^. This limitation makes INSTINCT unsuitable as a data storage repository and impractical for integrating large-scale datasets containing a substantial number of spots. Thirdly, although INSTINCT has relatively low time costs compared to baseline methods, typically completing the integration task within several minutes, its memory usage is relatively high (Supplementary Text 8, Supplementary Fig. 37, Supplementary Tabs. 1 to 3).

Moreover, we envision potential avenues for further optimizing INSTINCT. Firstly, inspired by methods developed for SRT data^31^, exploring an efficient cell-type decomposition method and embedding it into INSTINCT could elevate spATAC-seq data to single-cell resolution, enabling more precise exploration of biological properties at different scales. Secondly, constructing associations between different samples through effective alignment algorithms could offer a 3D perspective^29,31^,95, facilitating a more comprehensive study of the influence of the microenvironment. Thirdly, by replacing the GAT encoder with an MLP, INSTINCT can be adapted for integrating multi-batch single-cell omics data that do not contain spatial information (Supplementary Text 9, Supplementary Fig. 38). Fourthly, the structure of the decoder can be altered to a single-layer fully connected network. By multiplying the weight parameter matrix of the decoder and the matrix composed of the principal component vectors obtained in the data preprocessing stage, we can establish the connection between the low-dimensional representations and the original peaks, thereby enabling the identification of spot-type-specific or cluster-specific peaks in a model-specific way^65^. Lastly, we have demonstrated that using cosine similarity to find neighbors is superior to using Euclidean distance for the integration of spATAC-seq data. Other mechanisms more suitable for this data modality should also be gradually explored.

## Methods

### Data preprocessing

A single sample (slice) of spATAC-seq dataset consists of a spot-by-peak data matrix and a corresponding spatial coordinate matrix. We define *E*= [*e*_*ij*_] ∈ ℕ^*n*×*p*^ as the data matrix of a single slice, where each row represents a spot and each column represents a feature, specifically for spATAC-seq data, a peak. *E* is typically a high-dimensional matrix with high sparsity. We take *D*= [*d*_*ij*_] ∈ ℝ^*n*×2^ as the spatial coordinate matrix, where each row represents a 2D coordinate vector corresponding to a barcoded spot on the sequencing array. We employed a standardized preprocessing procedure for multi-samples 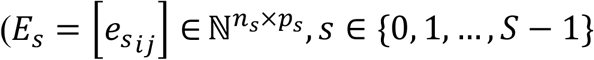, where *S* is the total number of slices, and *n*_*s*_ and *p*_*s*_ are the number of spots and number of peaks for the *s*-th slice, respectively) prior to integration. Given that each sample each possesses its specific set of peaks as features, and peaks from different peak sets may exhibit overlapping regions but are typically not identical, it become impractical to directly concatenate the data matrices of different samples.

Therefore, we initially conducted a peak merging process across samples according to the instructions provided by Signac^96^. To be specific, we first obtained the specific peak set for each slice, where each peak contains information including the chromatin number, start site, and sequence length. We then combined these specific peak sets into a unified peak set and proceeded to merge the peaks. If any two peaks overlapped or were adjacent (the end site of one peak matching the start site of another on the same chromatin), we merged them and defined the shortest region encompassing these peaks as the merged peak. This process continued until no two peaks overlapped or were adjacent, resulting in a merged peak set. We then compared this merged peak set with the specific peak set of each slice. A peak from the merged peak set was retained if at least one peak in each specific peak set was found to be a subregion of that peak; otherwise, it was removed. We then filtered out the peaks in the merged peak set with sequence lengths greater than 10,000 bp or less than 20 bp. Finally, the values from the original data matrix were transferred to the corresponding merged peaks and accumulated to form a series of new data matrices (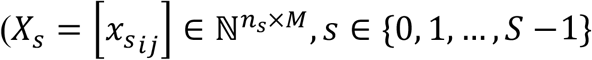, *s* ∈ {0, 1, …, *S* − 1}, where *M* is the number of merged peaks for all slices) with an identical peak set. We then concatenated all the slices to form a larger data matrix *X*_*concat*_ ∈ ℕ^*N*×*M*^, where 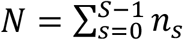 is the total number of spots across all slices. We have tested the time cost of the peak merging process and found it acceptable. Even when merging 10 slices with nearly 30,000 spots and over 3.28 million peaks, numbers significantly exceeding those of typical spATAC-seq datasets, the process was completed within half an hour (Supplementary Tab. 4).

We converted the concatenated read count matrix into a fragment count matrix following a previously established method^97^. It has been proven that the fragment count matrix better preserves quantitative regulatory information, thereby enhancing the analysis of ATAC-seq data^97^.

Subsequently, according to the sequencing depth of different datasets, we filtered out peaks accessible in spots fewer than a certain proportion and removed spots with zero count using EpiScanpy (v0.4.0) package^98^. For the spatial ATAC-RNA-seq MB dataset^16^, this proportion was set to 0.02; for the spatial-ATAC-seq ME dataset^13^, this proportion was set to 0.05; and for the spatial ATAC ME dataset^12^, this proportion was set to 0.003. For other datasets, it was set to 0.03. Then, we applied the term frequency-inverse document frequency (TF-IDF) transformation, which has been widely used as the scaling method for epigenomics data^7,66,96,99-101^, on the concatenated data matrix *X*_*concat*_. Finally, we performed PCA on the transformed matrix and retained the top 100 principal components as *X*_*pca*_ ∈ ℝ^*N*×100^. For ease of subsequent elaboration, we refer to *X*_*pca*_ as *X*.

### Construction of spatial neighbor graph

To fully leverage the spatial information provided by spATAC-seq data, we constructed a spatial graph based on the spatial coordinate matrices. Initially, we computed pairwise distances between spots within each individual slice. The thresholds for adjacency determination of each slice was set to 1.5 times the minimum distance between spots within that slice. Pairs of spots closer than the corresponding threshold were deemed adjacent, while those farther apart were considered non-adjacent, resulting in a symmetric adjacency matrix 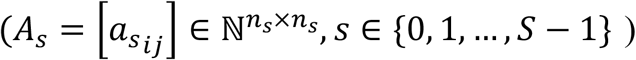 for each slice. Here, 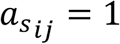 indicates adjacency between the *i*-th and *j*-th spot in the *s*-th slice, while 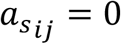 denotes non-adjacency. These matrices were then arranged diagonally to construct an overall spatial neighbor graph *A* ∈ ℕ^*N*×*N*^, where spots from different slices were considered non-adjacent.

### The model of INSTINCT

INSTINCT is an integration method which takes the first 100 principal components of the TF-IDF-transformed transformation data matrix (*X*) and the overall spatial neighbor graph (*A*) as input, and generates a low-dimensional representation for each spot in all slices where batch effects have been removed. It employs a stochastic domain translation procedure^39-41^, trained through adversarial learning^102^, as the key mechanism responsible for batch correction and the preservation of biological variations. The low-dimensional representations of spots can be used for spatial domain identification, visualization, spot-type annotation and various downstream analyses such as motif enrichment analysis, expression enrichment analysis and partitioned heritability analysis to elucidate biological principles and facilitate biological discoveries.

The model of INSTINCT comprises four components: an encoder, a noise generator, a decoder and a discriminator. The structure and mechanism of these modules are detailed below.

#### Encoder

The encoder adopts a GAT, which, for each input node, integrates not only the information of the node itself but also that of its related nodes based on an input graph. This enables INSTINCT to fully leverage the spatial information provided by spATAC-seq data. The graph attention mechanism of the *l*-th layer is as follows:

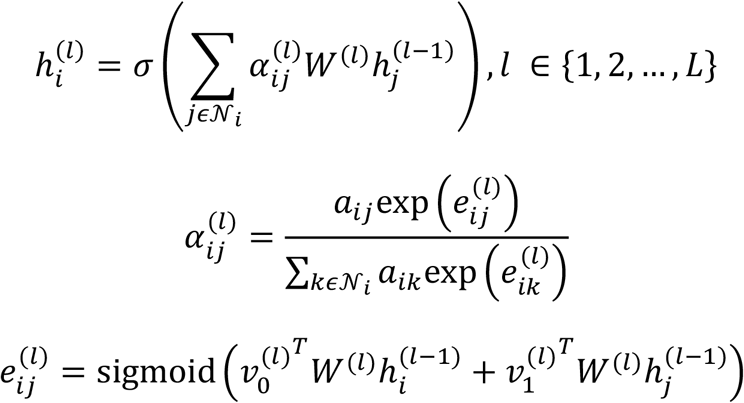

where 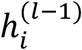 and 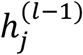 represent the input vectors of the *i*-th and *j*-th spots, respectively, while 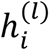 denotes the output representation of the *i*-th spot. 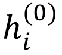 is equivalent to the *i*-th row of *X. W*^(*l*)^, 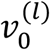 and 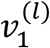 are the trainable parameters of the *l*-th layer. *a*_*ij*_ signifies the attention score of the *i*-th and *j*-th spots for the *i*-th spot, measuring the impact of the *j*-th spot on the *i*-th spot. σ(·) is the activation function. The output of the *L*-th layer is the low-dimensional latent representation matrix *Z*= [*z*_*ij*_] ∈ ℝ^*N*×*P*^, where *P* represents the dimensionality of the shared latent space.

#### Noise generator

The noise generator is structured as a linear network comprising only a single layer, dedicated to capturing the pattern of the noise and batch effects for each spot. It takes as input the concatenation of the low-dimensional representation matrix *Z* and a one-hot matrix *Q* ∈ ℝ^*N*×*S*^, which encodes the slice index of each spot. Specifically, each row of *Q* is the one-hot vector indicating the slice index of the corresponding spot, with a dimension equal to the number of slices.

We assume that the noise and batch effects of each spot are influenced by the characteristics of that spot, the characteristics of its surrounding environment, and the specific attributes of the slice it belongs to. Since the encoder utilizes graph attention mechanism, the representations of spots in *Z* already incorporates information of each spot and its surrounding environment, while the slice-specific information is encoded in *Q*.

The noise generator outputs a matrix *N*_*X*_ ∈ ℝ^*N*×*P*^, identical in size to *Z*, where each row signifies the representation of noise and batch effects for the corresponding spot in the low-dimensional space.

#### Decoder

The decoder is a MLP that takes as input the low-dimensional representation matrix containing noise and batch effects, and maps each spot to the domain of a specified slice. Since INSTINCT involves a stochastic domain translation procedure which translate each spot to different domains, a process to be detailed later, the decoder is responsible for both reconstructing spots back to domains of their original slices and mapping spots to the domains of other slices. The output of the decoder is either the reconstructed data matrix *X*_*rec*_ ∈ ℝ^*N*×100^ or the generated fake data matrix *Y* ∈ ℝ^*N*×100^.

#### Discriminator

The discriminator is an essential component for adversarial learning. We employ the discriminator to simultaneously discern the authenticity of the data and the category of each spot. The discriminator has a shared first layer and separate second layers which result in two outputs: a vector *d*_*src*_ ∈ ℝ^*N*^ for predicting the authenticity of the data and a matrix *D*_*cls*_ ∈ ℝ^*N*×*S*^ for predicting the category of the data, i.e., the sample index of each spot. In *d*_*src*_, higher values indicate that the representation of the corresponding spot is more likely to be from real data. This architecture, widely used in image processing for adversarial training^39,40^, offers the advantage of flexible translation between multiple domains using a single encoder and decoder.

We designate *X* as real data, and *X*_*rec*_ and *Y* as fake data. The index of the slice where the spot originally belongs serves as the category label for the corresponding spot in *X* and *X*_*rec*_. Additionally, we assign the index of the specified target slice chosen during the stochastic translation procedure as the category label for the corresponding spot in *Y*, which will be detailed later.

The encoder of INSTINCT utilizes two single-head GAT layers. The first layer reduces the dimensionality of the input representations of spots to 50, and the second layer further reduces it to 30. We use ELU^103^ as the activation function between these two layers. The decoder is a MLP comprising two layers: the first layer expands the dimensionality to 50, and the second layer further expands it back to 100. ELU is also used as the activation function in the decoder. The shared first layer of the discriminator reduces the dimensionality to 50, and the separate second layers decrease the dimensionality to the dimension of the corresponding output, respectively. The noise generator is a single-layer linear network, with the sum of the latent space dimension (30) and the total number of slices as the input dimension, and 30 as the output dimension.

INSTINCT adopts a two-stage strategy. In stage 1, we exclusively utilize the encoder, decoder, and discriminator to construct the model, with the encoder and decoder together form the generator. The operational procedure unfolds as follows:

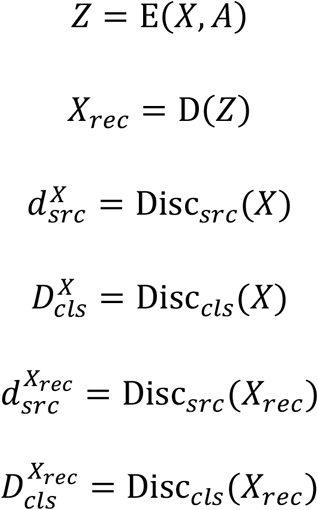

where E(·), D(·), Disc_*src*_(·) and Disc_*cls*_(·) denote the encoder, the decoder, the part where the discriminator distinguishes between real and fake data, and the part where the discriminator distinguishes between data categories, respectively. The encoder takes *X* and *A* as input, yielding *Z* as illustrated earlier, while the decoder directly uses *Z* as input to reconstruct the input data matrix. In this stage, we also pretrain the discriminator to distinguish between the fake data matrix *X*_*rec*_ and the real data matrix *X*, as well as to classify the spots based on the indices of their original slices.

Three types of loss functions are employed in stage 1 to train the model, including adversarial loss, classification loss, and reconstruction loss.

The adversarial loss is employed to encourage the generator to produce data more closely resembling real samples, thereby enhancing its capability, while also improving the precision of the discriminator to distinguish between samples generated by the generator and the real data. It is calculated as follow in stage 1:

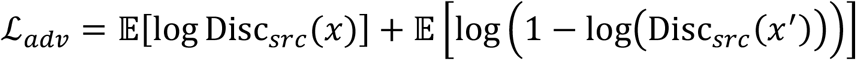

where *x* and *x*^′^ represent spot representation from *X* and *X*_*rec*_, respectively. The discriminator maximizes this function to distinguish between real and fake data, while the generator minimizes it to confound the discriminator.

The classification loss consists of the following two components:

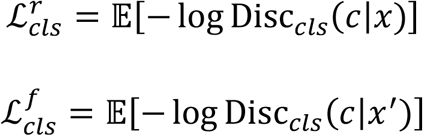

where *c* represents the category label (slice index) of the spot *x* or *x*^′^. We use 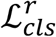 to train the discriminator to distinguish spots from different slices, and 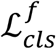 to train the generator to accurately map spots to domains of their original slices.

The reconstruction loss is used to enhance the ability of the model to reconstruct the input data. By minimizing the reconstruction loss, the model learns effective data representation and reconstruction techniques, thereby improving the quality and accuracy of its generated data. We used mean square error (MSE) as the reconstruction loss:

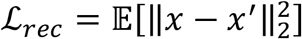

Therefore, the total objectives for the generator and discriminator in stage 1 are displayed below:

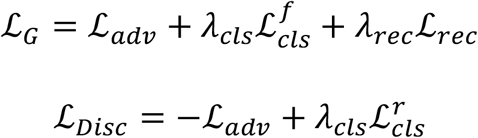

where *λ*_*cls*_ and *λ*_*rec*_ are hyperparameters which control the contribution of the classification loss and the reconstruction loss to the total objectives, respectively.

We declare that the training in stage 1 serves merely as a pretraining step, aimed at enabling the model to initially learn feature extraction and data reconstruction, and allowing the discriminator to acquire a certain level of data discrimination capability. This is expected to enhance the effectiveness of batch effect removal during training in stage 2. However, the low-dimensional representations learned in this step still contain noise and batch effects.

In stage 2, we add the noise generator module to the model and employ a stochastic domain translation process to facilitate the removal of noise and batch effects. The operational mechanism is is detailed below:

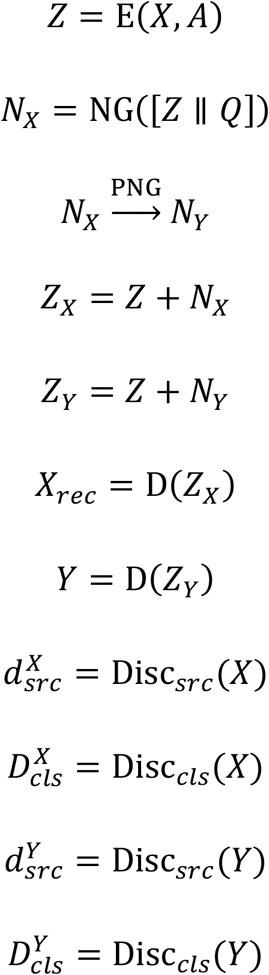

where *Q* is the one-hot matrix indicating the slice indices as stated above, NG(·) represents the noise generator, ∥ denotes concatenation, and PNG represents the PNG process.

First of all, taking *X* and *A* as input, the encoder integrates the information from each spot with its neighborhood information to generate *Z*, same as the process in stage 1.

After that, we concatenate *Z* with the one-hot matrix *Q*, which indicates the slice indices of spots as stated above, and input it to the noise generator. The noise generator is used to capture the patterns the of noise and batch effects for each spot in the latent space. It outputs *N*_*X*_, which contains the representations of noise and batch effects for spots. *N*_*X*_ is subsequently added to *Z* to generate the latent representations of spots with batch effects, denoted as *Z*_*X*_, which is then utilized to reconstruct the input data matrix via the decoder. Since the batch-specific information (*Q*) is only exposed to the noise generator and the decoder, the encoder is encouraged to only extract biological information from *X* when generating *Z*.

However, it is challenging to ensure that the noise generator exclusively captures noise and batch effects, as *Z* is also exposed to it. To address this, we employ a stochastic domain translation process that allows the noise generator to better simulate noise and batch effects, and further ensures that the encoder extracts sufficient biological variations, which are crucial for distinguishing different spot-types.

The core idea of stochastic domain translation is that by adding different patterns of batch effects to a spot in the shared latent space, the decoder can flexibly map it to the domain of any slice in the feature space, regardless of its actual slice of origin. This approach enhances the ability of the noise generator to capture the differences in batch effect patterns across slices, thereby enabling the encoder to retain sufficient biological variations. The discriminator is used to ensure the effectiveness of the translation process through discrimination and classification.

Based on *N*_*X*_, we generate *N*_*Y*_ through the PNG process to simulate the representations of noise and batch effects that enable spots to be mapped to the domains of other slices. Specifically, during each training epoch in stage 2, each slice is assigned at random an index, representing the domain of the target slice to which the spots in the source slice (the original slice of spots) should be mapped during that epoch. Subsequently, for the *i*-th source spot in the *s*-th source slice, we find *k* nearest neighbors within the target slice for the source spot using the representations in *Z*. The nearest neighbors are selected based on the magnitude of cosine similarity. These neighbors are spots within the target slice that are most likely to possess similar biological characteristics as the source spot. We then average the representations of the noise and batch effects corresponding to these spots in *N*_*X*_ to determine the noise and batch effects required for mapping the source spot to the domain of the target slice. By repeating this process for each spot, we obtain *N*_*Y*_, where each row represents the pseudo-noise-and-batch-effects for the corresponding spot. *N*_*Y*_ is then added to *Z* to yield *Z*_*Y*_, after which the decoder is used to map each spot to its designated target domain.

In stage 2, we task the discriminator with distinguishing between the real data matrix *X* and the fake data matrix *Y*, while also discerning the categories of spots. Here, the categories of spots corresponding to *Y* are defined by the target domains.

Four types of loss functions are employed in stage 2 to train the model, including adversarial loss, classification loss, reconstruction loss, and a new loss function called latent loss.

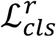 and ℒ_*rec*_ remain the same as in stage 1, while ℒ_*adv*_ and 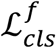 are calculated as follows:

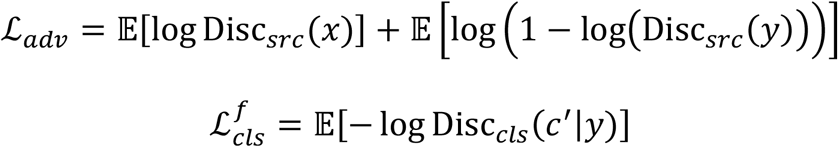

Where *y* represent spot representation *Y*, and *c*^′^ is the assigned target category correspond to *y*. 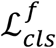 is used to train the generator to accurately map spots to domains of target slices.

We developed the latent loss to better remove the batch effects between samples:

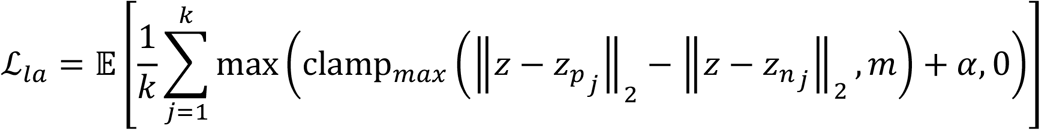

where *z* is the spot representation from *Z*. For each spot, we identify *k* positive samples and *k* negative samples. A positive sample, a negative sample, and the spot form a triplet. We use triplet loss^104^ to encourage the spot to be close to its positive samples and far from negative samples in the latent space. The positive samples are the nearest neighbors selected for the spot during the PNG process, while the negative samples are randomly chosen from the same slice as the spot, following the manner of STAligner^28^. Here, 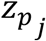 and 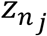 represent the *j*-th positive sample and the *j*-th negative sample found for *z*, respectively, and *α* denotes the hyperparameter of triplet loss.

However, we cannot guarantee that all positive samples found for a particular spot are the same type as that spot. In fact, if a spot belongs to a type specific to a certain slice, or if the number of spots of that type in other slices is less than *k*, then some or all of the positive samples found for it will not be of the same type as the spot itself. Using triplet loss to encourage the spot to be close to these incorrect positive samples may result in the inability to identify rare or specific spot-types, which is undesirable. Additionally, since negative samples are randomly selected, it is impossible to avoid selecting samples of the same type as the spot. Therefore, we employ a clamp strategy, denoted as clamp_*max*_(·). Triplets where the difference between the distance of the spot to the positive sample and the distance of the spot to the negative sample exceeds a hyperparameter *m* are not included in the backpropagation process. This strategy helps in removing positive samples that are too distant from the spot (potentially incorrect positives) or negative samples that are too close to the spot (potentially incorrect negatives).

The total objectives for stage 2 are as follows:

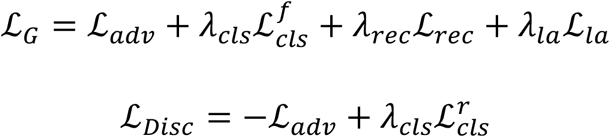

where *λ*_*la*_ controls the contribution of latent loss fucntion to the total objectives. It is important to note that the forms of ℒ_*adv*_ and 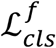 have changed compared to those in stage 1, as stated above.

Compared to conventional autoencoders, the incorporation of the noise generator and discriminator modules, along with the utilization of the stochastic domain translation procedure, enables the low-dimensional representations of spots with added noise and batch effects to be seamlessly reconstructed back to the domain of their original slices or generated to any specified domain of another slice. This approach allows the noise generator to effectively capture noise and batch effect patterns across different slices, thereby facilitating the more efficient removal of batch effects from the low-dimensional representations while preserving as much biological variations as possible. After training in both stages, *Z* can be extracted as the low-dimensional representations of spots without noise and batch effects, suitable for various applications and downstream analyses.

For all experiments reported in this paper, we set *λ*_*cls*_ = *λ*_*rec*_= 10, *λ*_*la*_ = 20, *k* = 50, *m* = 10, and *α* = 1. The generator was optimized with a learning rate of 0.001, while the discriminator used 0.0005. The number of training epochs for both stages were set to 500.

To assist users in using INSTINCT under various scenarios, we conducted sensitivity analysis on all the hyperparameters involved in our method and provided guidance for parameter settings (Supplementary Text 10, Supplementary Figs. 39 to 41, Supplementary Tabs. 5 and 6).

### Simulations

We conducted extensive simulation experiments to validate the performance of INSTINCT. Inspired by SpatialPCA^44^, a method developed for SRT, we adopted a similar approach to generate spATAC-seq datasets. The data information for different scenarios are summarized in Supplementary Tabs. 7 to 13.

Specifically, we utilized simCAS^45^, a method for simulating scATAC-seq data, to generate multiple single-cell datasets with batch effects. For the reference data and parameters used in the simulation, we employed the discrete mode of simCAS, with the *tree_text* parameter set to “(((0:0.2, 1:0.2):0.2, 2:0.4):0.5, (3:0.5, 4:0.5):0.4);”, and followed its online tutorial. We utilized the summary statistics of the Buenrostro2018 dataset^105^ provided at https://github.com/Chen-Li-17/simCAS/tree/main/data as the training data. Besides, simCAS provides an optional step for adding batch effects to simulated data. According to simCAS, batch effects are categorized into those originating from technical variations and those originating from biological factors. By adjusting the parameters for introducing these two types of batch effects, we produced three scATAC-seq datasets exhibiting varying degrees of batch effects. Each dataset comprises 10,000 cells distributed among five cell-types, with 2,000 cells for each type labeled as 0 to 4, respectively. The parameters for the batch effects added to these three datasets were as follows: for dataset 0, we set the mean of batch effects from biological factors to 0 with a standard deviation of 0.5. For dataset 1, we set the mean of batch effects from technical variations to 0 with a standard deviation of 0.1. For dataset 2, we set the mean of batch effects from technical variations to 0 with a standard deviation of 0.1 and the mean of batch effects from biological factors to 0 with a standard deviation of 0.5.

After that, we pre-defined a single-cell resolution slice of size 120 × 150 as a template, where each location corresponds to integer coordinates ranging from 0 to 149 for the x-axis and 0 to 119 for the y-axis. This template was divided into five domains represented by domain 0 to 4, respectively. Each domain contained a different number of locations: 2700, 4500, 3150, 3600, and 4050, each corresponding to a major cell-type with the same number as its index. Initially, each position was assigned a cell-type, where in a specific region, the cell-type corresponding to that region was assigned with a probability of *r*, while the remaining cell-types were each assigned with a probability of 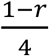. Subsequently, based on the assigned cell-types for each location, a cell corresponding to that cell-type was randomly selected from the scATAC-seq dataset and allocated to that position. By repeating this process for each simulated scATAC-seq dataset, we obtained three single-cell resolution slices. Each spot, defined as a 3 × 3 cell grid, aggregated data from the nine cells within it by summation, with the prevalent cell-type among the nine cells dictating the type of that spot. The coordinates of the spot were determined by the coordinates of the central cell. Thus, three single-cell resolution slices were transformed into corresponding spatial resolution slices theoretically containing five spot-types, denoted as spot-type 0 to 4. We iteratively decreased the value of *r*, resulting in increasing levels of confusion among spot-types and progressively more complex batch effects across three scenarios, where scenarios 1, 2, and 3 corresponded to *r* values of 0.8, 0.7, and 0.6, respectively. We point out that the value of *r* should not be too small because it may lead to a discontinuous ground truth, resulting in pre-defined domains losing their practical significance.

In certain situations, vertically adjacent slices may not exhibit complete overlap, where different slices display varying proportions of spot-types, or a slice may lack some spot-types found in other slices. This issue can also arise in samples from different developmental stages. Therefore, based on scenario 1, we created scenarios 4 and 5 to simulate two cases where the proportions of spot-types differ between slices (Supplementary Fig. 8). In scenario 4, we applied different cropping to the three spatial resolution slices from scenario 1, preserving 80% of the spots on the right side, middle, and left side, respectively, to generate three new slices. In scenario 5, we retained slice 0 and slice 1 from scenario 1 and removed domain 2 from slice 2.

We simulated scenario 6 based on scenario 1 to test the ability of INSTINCT to preserve rare and specific spot-types. We excluded domain 1 from slices 1 and 2 of scenario 1, retaining a 5 × 5 area of domain 1 in slice 0 and removing the rest (Supplementary Fig. 8). Only about 25 spots of spot-type 1 were kept across the three slices, mostly concentrated in slice 0. It is worth noting that, due to the specific way we determined spot-types, there are several spots belonging to spot-type 1 in other domains.

To further explore the ability of INSTINCT to distinguish rare spot-types, we designed simulated scenario 7, which comprises more spot-types compared to the previous six scenarios. Specifically, we employed the discrete mode of simCAS, with the *tree_text* parameter set to “((((0:0.2,1:0.2):0.2,2:0.4):0.5, (3:0.5,4:0.5):0.4):0.1, ((5:0.1,6:0.4):0.3, 7:0.2):0.2);”, producing three scATAC-seq datasets with varying batch effects. Each dataset comprises 8,000 cells distributed among eight cell-types, with 1,000 cells per type labeled as 0 to 7. Then, we introduced another pre-defined single-cell resolution slice as the template and generated three spatial resolution slices, each comprising eight spot-types. Each slice contains a total of 2,000 spots, with spot-types 5, 6, and 7 being rare, comprising approximately 50 spots per slice.

### Spatial domain identification

For datasets with provided annotations, we initially applied a Gaussian mixture model-based clustering algorithm implemented in the scikit-learn (v1.4.0) package^106^, setting the number of components to the number of spot-types, following the approach of STitch3D^31^, to cluster the latent representations of spots in *Z*. Subsequently, we utilized the *match_cluster_labels* function provided in the online tutorial of PASTE^107^ to map the cluster labels to the provided annotations. For datasets without annotations, we applied the Leiden algorithm^108^ implemented in the Scanpy (v1.9.8) package^109^ with the default settings to cluster the latent representations of spots.

### Visualization

For visualizing the integration results, we used the uniform manifold approximation and projection (UMAP) algorithm^110^ to further reduce the dimensionality of the spot representations in *Z* and visualized them in a two-dimensional space. Following Portal^111^, the algorithm was run with parameters set to 30 nearest neighbors, a minimum distance of 0.3 and utilizing the correlation metric. For visualizing the raw data, we initially preprocessed the data following the procedure outlined above. Subsequently, we employed UMAP algorithm with the same settings to reduce the dimensionality of the spot representations in *X*_*pca*_.

### Spot-type annotation

We performed spot-type annotation by employing a kNN classifier, a model-free approach whose annotation efficacy heavily depends on the effectiveness of the low-dimensional representations across samples. For each spot in the target sample to be annotated, we selected a specified number of nearest neighbors from the annotated reference sample. The label assigned to each spot was determined by the majority class among its nearest neighbors. We set the number of nearest neighbors to 20, consistent with the approach outlined in Portal^111^.

### Gene ontology analysis

Initially, we conducted differential gene expression analysis using the *rank_genes_groups* function in the Scanpy package with the Wilcoxon test as the method. A gene was considered differentially expressed if it exhibited a log2-fold change greater than 0.2 and an adjusted *p*-value (*q*-value) less than 0.05. Subsequently, the DEGs were utilized for performing GO analysis via the algorithm provided in the <u>GSEApy</u> (v1.1.2) package^112^, employing *GO_Biological_Process_2021* as the reference database.

### Determination of spot-type-specific peaks

We first filtered the peaks according to the preprocessing procedure and applied TF-IDF transformation to the sample. Then, we utilized the *rank_genes_groups* function in the Scanpy package with the Wilcoxon test as the method. For each spot-type, we filtered out peaks those exhibited log2-fold change no more than 0.2, and selected 300 peaks with the lowest adjusted *p*-value (*q*-value) as its specific peaks. For spot-types whose peaks were no more than 300 after filtering, we kept all peaks as their specific peaks. Peaks not chosen as specific peaks were retained as background peaks.

### Expression enrichment analysis

Following EpiAnno^65^, we initially mapped the genome coordinates of the spot-type-specific peaks and background peaks to GRCh37/hg19 using LiftOver^113^. Subsequently, we conducted expression enrichment analysis utilizing the SNPsea (v1.0.3) package^114^ with default settings for SNPs in each set of spot-type-specific peaks and the set of background peaks, respectively.

### Partitioned heritability analysis

We first mapped the genome coordinates of the spot-type-specific peaks and background peaks to GRCh37/hg19. Then, we used LDSC (v1.0.1) package^115^ to quantified the enrichment of heritability for phenotypes related to brain. Following CASTLE^101^, we used European samples from the 1000 Genomes Project as the reference panel, and the summary statistics for GWAS were all obtained from https://doi.org/10.5281/zenodo.7768714.

### Motif enrichment analysis

We performed motif enrichment analysis following previous researches^63-66^. First, we applied the *rank_genes_groups* function in the Scanpy package^109^, using the Wilcoxon test as the method, to the data. For each cluster, we filtered out peaks that exhibited a log2-fold change of no more than 0.2 and selected 1,000 peaks with the lowest adjusted *p*-value (*q*-value) as its specific peaks. We combined the cluster-specific peaks of each cluster into a peak set and used the data corresponding to these peaks as the filtered low-dimensional data, and then utilized chromVAR (v1.26.0) package^63^ to infer the enriched TF binding motifs within these peaks.

### Data collection and description

Detailed information of spATAC-seq and SRT samples are listed in Supplementary Tab. 1.

### Baseline methods

We compared INSTINCT with six baseline methods, including Scanorama^21^, SCALEX^25^, PeakVI^26^, SEDR^27^, STAligner^28^, and GraphST^29^. Among these, Scanorama is originally designed for integrating scRNA-seq data. SCALEX is designed for online integration of single-cell omics data, and its superior performance in integrating scATAC-seq data has been demonstrated in the original study^25^. PeakVI is specifically developed for scATAC-seq data integration. SEDR, STAligner, and GraphST are methods developed for integrating SRT data.

For each method, we conducted eight iterations of experiments by varying the random seed. Data were filtered before used, and all parameters were maintained at their default settings. We followed the tutorial provided by each method to conduct the experiments.

### Evaluation metrics

We compared the capabilities of the integration methods in three aspects: clustering performance, representation quality, and batch correction. We regarded clustering performance and representation quality as two separate aspects of biological conservation, noting that representation quality depends solely on the model, whereas clustering performance also relies on the clustering algorithm. We assessed clustering performance using six metrics: Adjusted Rand Index (ARI), Adjusted Mutual Information (AMI), Normalized Mutual Information (NMI), Fowlkes-Mallows Index (FMI), completeness score (Comp), and homogeneity score (Homo), leveraging functions from the scikit-learn package^106^. For evaluating representation quality, we employed mean average precision (mAP)^116^, average silhouette width across spot-types (spot-type ASW), average silhouette width across isolated labels (isolated label ASW), and isolated label F1-score, utilizing the scib (v1.1.4) package^47^. Regarding batch correction assessment, we computed average silhouette width across batches (batch ASW), principal component regression across batches (batch PCR), k-nearest-neighbor batch effect test (kBET), and graph connectivity using the scib package. The scores of each group were averaged to obtain the corresponding group-specific overall score. The group-specific overall scores for the three aspects were further averaged to obtain the final overall score.

For the evaluation of spot-type annotation, we calculated accuracy, kappa, macro F1-score (mF1), and weighted F1-score (wF1) using k-fold cross-validation, with each fold corresponding to one slice.

#### ARI

ARI standardizes the Rand Index (RI) to quantify the similarity between two sets of labels. The higher the ARI value, the more similar the two sets of clusters are. The formula is as follows:

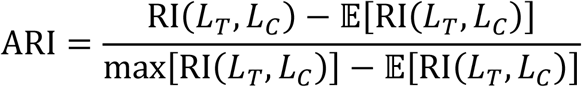

where *L*_*T*_ and *L*_*C*_ are true labels and clustering labels, respectively. RI(·) denotes the function of RI, and 𝔼[RI] represents the expected RI under random assignment.

#### AMI

AMI is a measure used to compare the similarity between two sets of labels by adjusting the Mutual Information (MI) score. The higher the AMI value, the more similar the two sets of clusters are. AMI score is calculated as follows:

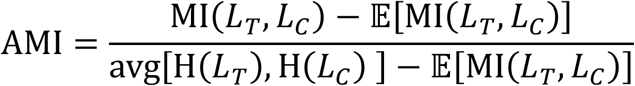

where MI(·) denotes the function of MI and H(·) represents the function of entropy.

#### NMI

NMI calculates the normalization of MI score. Higher NMI score represents better agreement between the two sets of clusters. The formula is as follows:

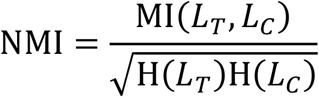

#### FMI

FMI focuses on the precision and recall of the clustering results, with a higher score indicating better clustering performance. It is calculated as follows:

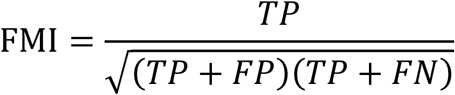

where *TP* is the number of pairs of elements that are in the same cluster in both the true labels and the clustering labels, *FP* is the number of pairs of elements that are in the same cluster of the true labels but in different clusters of the clustering labels, and *FN* is the number of pairs of elements that are in different clusters of the true labels but in the same clusters of the clustering labels.

#### Comp

Completeness score evaluates if members of the same true label class are grouped together, with a higher score indicating better grouping. It is calculated as follows:

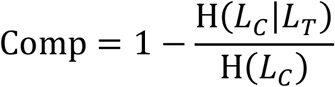

#### Homo

Homogeneity score evaluates whether a single cluster contains only data samples from a single class according to the provided annotations, with a higher score indicating better grouping. The formula is as follows:

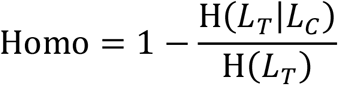

#### mAP

mAP quantifies how well the labels of the nearest neighbors of each spot match its own label, with a higher value indicating the better preservation of the consistency. For the calculation methods, we followed its original work^116^. The number of the nearest neighbors considered was set to 30.

#### Spot-type ASW

The silhouette width is a measure of how similar a sample is to others of its own class compared to different classes^117^, with ASW representing the average silhouette width for all samples. Spot-type ASW is the ASW score calculated with respect to spot-type labels, ranging from -1 to 1, where a higher score indicates that spots are more similar to those of their own type than to others. We used the function implemented in the scib package^47^, which scales the spot-type ASW score to a value between 0 and 1, with a higher score indicating better representation quality. The scaling formula is as follows:

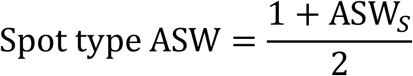

where ASW_*S*_ is the original spot-type ASW score before scaling.

#### Isolated label ASW

Isolated label ASW evaluates the conservation of spot-types unique to individual samples by calculating the ASW with respect to these labels. It ranges from -1 to 1, where a higher score indicates that spots are more similar to those of their own type than to others. We computed it with the function implemented in the scib package, which scales the isolated label ASW score to a value between 0 and 1, with a higher score indicating better preservation of unique spot-types. The scaling formula is as follows:

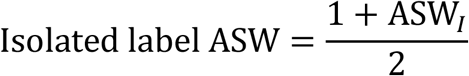

where ASW_*I*_ is the original isolated label ASW score before scaling.

#### Isolated label F1-score

Isolated label F1-score is also a metric developed to assess the preservation of the spot-types unique to individual samples. It calculates the F1-scores^118^ for these slice-specific spot-types and takes the mean. It returns a value between 0 and 1, with a higher score indicating better preservation of unique spot-types. We used the function implemented in the scib package and kept its default settings.

#### Batch ASW

Batch ASW measures of how similar a sample is to its own slice compared to other slices, calculated with respect to slice indices. It ranges from -1 to 1, with a score closer to 0 indicating that spots from different slices are well mixed. We computed it using the function implemented in the scib package, which scales the score to a value between 0 and 1, where a higher value indicates a better mixing of data from different slices. The formula is as follows:

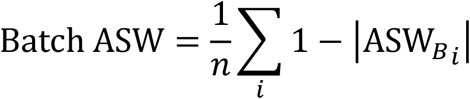

where 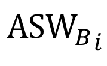 is the original batch ASW score calculated for the *i* -th spot-type before scaling, and *n* is the number of spot-types.

#### Batch PCR

Batch PCR evaluates the effectiveness of batch correction by comparing the variance contribution of batch-specific information to the data matrix before and after integration^47,119^. The concatenation of the raw data from multiple samples is the data matrix before integration and the concatenation of latent embeddings of spots represent the data matrix after integration. We used the function implemented in the scib package, which scales the batch PCR score to a value between 0 and 1, where a higher value indicates better elimination of batch-specific information. The scaling formula is as follows:

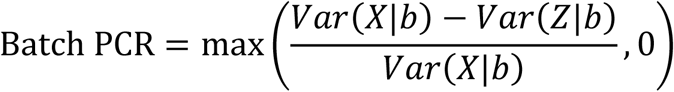

where *X* and *Z* represents the data matrix before and after integration, and *b* is the batch variable.

#### kBET

kBET measures the removal of batch effects by adopting a Pearson’s *χ*^2^-based test to determine whether the batch label distribution in the kNN of spots is similar to the global distribution of the batch labels^119^. It is calculated within each spot-types and returns the rejection rate, with a smaller value indicating better mixing of spots. We computed it using the function implemented in the scib package, which summarizes the kBET scores for all spot-types and scaled the overall score to a value between 0 and 1, where a higher value indicates better removal of batch effects. The scaling formula is as follows:

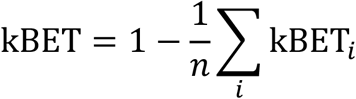

where kBET_*i*_ is the original kBET score calculated for the *i*-th spot-type before scaling, and *n* is the number of spot-types.

#### Graph connectivity

Graph connectivity measures whether the kNN graph directly connects all spots of the same type^47^. The algorithm is detailed as follows:

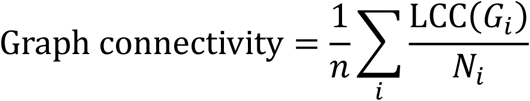

where *n* is the total number of spot-types, *N*_*i*_ is the number of spots of the *i*-th spot-type, *G*_*i*_ represents the subset kNN graph that contains only spots from the *i*-th spot-type, and LCC(·) returns the number of nodes in the largest connected component of the graph. The value of graph connectivity score is higher than 0 and no larger than 1, with a higher value indicating that more spots with the same identity are connected in the integrated kNN graph. We computed it with the function implemented in the scib package.

#### Accuracy

Accuracy is the proportion of correctly annotated spots to the total number of spots and is calculated as follows:

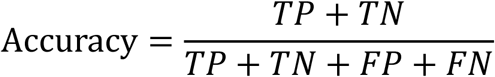

where *TP, FP, TN* and *FN* denote the number true positive samples, false positive samples, true negative samples, and false negative samples, respectively.

#### Kappa

Kappa measures inter-rater agreement for categorical items. It compares the observed agreement between two raters to the agreement that would be expected by chance, with a higher score indicating better annotation performance. The formula is as follows:

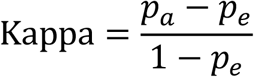

where *p*_*a*_ represents the proportion of observed agreement between two raters, and *p*_*e*_ is the proportion of agreement expected if the raters are to rate completely randomly.

#### mF1

mF1 takes the mean of the F1-scores^118^ calculated for all spot-types, with a higher score indicating better annotation performance. The formula is as follows:

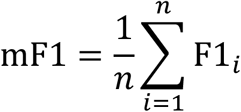

where *n* is the number of spot-types and F1_*i*_ represents the F1-score of the *i*-th spot-type.

#### wF1

wF1 calculates the weighted average of F1-scores for all spot-types, where the weight of each F1-score is the proportion of the number of spots for that type relative to the total number of spots. A higher score indicating better annotation performance. It is computed as follows:

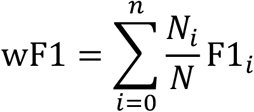

where *N*_*i*_ denotes the number of spots of the *i*-th spot-type, and *N* is the total number of spots.

## Data availability

The MISAR-seq MB dataset^15^ can be downloaded from the National Genomics Data Center with accession number OEP003285. The spatial ATAC-RNA-seq MB dataset^16^ is available at the Gene Expression Omnibus with accession code GSE205055. The spatial-ATAC-seq ME dataset^13^ is deposited in the Gene Expression Omnibus with accession code GSE171943. The spatial ATAC ME dataset^12^ can be obtained at Gene Expression Omnibus using accession code GSE214991.

## Code availability

INSTINCT and all the code for reproducing the analyses and benchmarking can be found at https://github.com/yyLIU12138/INSTINCT. Detailed tutorials can be found at https://instinct.readthedocs.io/en/latest/index.html.

## Acknowledgements

This work was supported by the National Key Research and Development Program of China (grant nos. 2023YFF1204802 and 2021YFF1200902), the National Natural Science Foundation of China (grant nos. 62273194), and Beijing Natural Science Foundation (L242026).

## Author contributions

R.J. conceived the study and supervised the project. Y.L., Z.L. designed and implemented and validated INSTINCT. X.Y.C., X.J.C. and Z.G. helped with analyzing the results. Y.L., Z.L. and R.J. wrote the manuscript, with input from all the authors.

