## Supplementary Materials for "INSTINCT: Multi-sample integration of spatial chromatin accessibility sequencing data via stochastic domain translation"

### Supplementary Tables

#### Supplementary Tab. 1 | Summary of the real data used in our work.

| Name | Index | n_spots | n_peaks_origin | n_peaks_merged |
|---|---|---|---|---|
| **MISAR-seq MB** | | | | |
| E11_0-S1_ATAC | 1 | 1258 | 242024 | 1+2+3+4: 99311<br>1+5: 128708<br>2+6: 164185<br>3+7: 182687<br>4+8: 176215 |
| E13_5-S1_ATAC | 2 | 1777 | 271126 | |
| E15_5-S1_ATAC | 3 | 1949 | 265014 | |
| E18_5-S1_ATAC | 4 | 2129 | 294734 | |
| E11_0-S2_ATAC | 5 | 1353 | 213562 | |
| E13_5-S2_ATAC | 6 | 2183 | 251555 | |
| E15_5-S2_ATAC | 7 | 1939 | 244394 | |
| E18_5-S2_ATAC | 8 | 2248 | 223333 | |
| E11_0-S1_RNA | 9 | 1263 | 32285 | |
| E13_5-S1_ RNA | 10 | 1777 | 32285 | |
| E15_5-S1_ RNA | 11 | 1949 | 32285 | |
| E18_5-S1_ RNA | 12 | 2129 | 32285 | |
| E11_0-S2_ RNA | 13 | 1353 | 32285 | |
| E13_5-S2_ RNA | 14 | 2183 | 32285 | |
| E15_5-S2_ RNA | 15 | 1939 | 32285 | |
| E18_5-S2_ RNA | 16 | 2248 | 32285 | |
| **spatial ATAC-RNA-seq MB** | | | | |
| P21_1_ATAC | 17 | 2500 | 233155 | 17+18+19: 178207 |
| P21_1_ATAC | 18 | 2500 | 290047 | |
| P22_ATAC | 19 | 9994 | 517411 | |
| P21_1_ RNA | 20 | 2373 | 19859 | |
| P21_1_ RNA | 21 | 2498 | 20046 | |
| P22_ RNA | 22 | 9215 | 22914 | |
| **spatial-ATAC-seq ME** | | | | |
| E11 | 23 | 2500 | 290412 | 23+24+25: 166114 |
| E13_1 | 24 | 2500 | 593976 | |
| E13_2 | 25 | 2500 | 289542 | |
| **spatial ATAC ME** | | | | |
| E12.5-S1 | 26 | 2234 | 248047 | Already merged in the original study |
| E12.5-S2 | 27 | 2246 | 248047 | |
| E13.5-S1 | 28 | 2970 | 248047 | |
| E13.5-S2 | 29 | 3160 | 248047 | |
| E15.5-S1 | 30 | 3903 | 248047 | |
| E15.5-S2 | 31 | 3274 | 248047 | |

#### Supplementary Tab. 2 | Time cost of INSTINCT and baseline methods on real datasets.

| Indices | INSTINCT | Scanorama | SCALEX | PeakVI | SEDR | STAligner | GraphST |
|---|---|---|---|---|---|---|---|
| **MISAR-seq MB** | | | | | | | |
| **1~4** | 1 min 9 sec | 0 min 38 sec | 17 min 18 sec | 5 min 37 sec | 0 min 31 sec | 0 min 36sec | 2 min 25 sec |
| **spatial ATAC-RNA-seq MB** | | | | | | | |
| **17~19** | 3 min 50 sec | 0 min 46 sec | 16 min 10 sec | 2 min 42 sec | 0 min 49 sec | 0 min 48 sec | 1 min 42 sec |
| **spatial-ATAC-seq ME** | | | | | | | |
| **23~25** | 1 min 9 sec | 0 min 27 sec | 14 min 27 sec | 2 min 12 sec | 0 min 23 sec | 0 min 28 sec | 0 min 57 sec |
| **spatial ATAC ME** | | | | | | | |
| **26~31** | 3 min 52 sec | 2 min 25 sec | 22 min 18 sec | 3 min 3 sec | 0 min 48 sec | 1 min 40 sec | 5 min 27 sec |
| **Scanorama (CPU): Intel(R) Xeon(R) Platinum 8352V Others (GPU): RTX 4090** | | | | | | | |

#### Supplementary Tab. 3 | Memory usage of INSTINCT and baseline methods on real datasets.

| Indices | INSTINCT | Scanorama | SCALEX | PeakVI | SEDR | STAligner | GraphST |
|---|---|---|---|---|---|---|---|
| **MISAR-seq MB** | | | | | | | |
| **1~4** | 5619 MiB | 4127 MiB | 2781 MiB | 1249 MiB | 1273 MiB | 1257 MiB | 6107 MiB |
| **spatial ATAC-RNA-seq MB** | | | | | | | |
| **17~19** | 21147 MiB | 7222 MiB | 2897 MiB | 739 MiB | 2037 MiB | 999 MiB | 14463 MiB |
| **spatial-ATAC-seq ME** | | | | | | | |
| **23~25** | 6809 MiB | 4127 MiB | 2359 MiB | 1635 MiB | 1237 MiB | 735 MiB | 5209 MiB |
| **spatial ATAC ME** | | | | | | | |
| **26~31** | 18611 MiB | 8253 MiB | 4195 MiB | 535 MiB | 2657 MiB | 949 MiB | 17147 MiB |
| **Scanorama (CPU): Intel(R) Xeon(R) Platinum 8352V Others (GPU): RTX 4090** | | | | | | | |

#### Supplementary Tab. 4 | Results of tests for peak merging.

| Indices | n_spots | n_peaks_total | n_peaks_merged | time |
|---|---|---|---|---|
| **1+2** | 3035 | 513150 | 131336 | 4 min 18 sec |
| **1+2+3** | 4984 | 778164 | 109654 | 6 min 3 sec |
| **1+2+3+4** | 7113 | 1072898 | 99311 | 7 min 51 sec |
| **1+2+3+4+17** | 9613 | 1306053 | 74213 | 8 min 56 sec |
| **1+2+3+4+17+18** | 12113 | 1596100 | 69926 | 10 min 53 sec |
| **1+2+3+4+17+18+19** | 22107 | 2113511 | 69437 | 16 min 9 sec |
| **1+2+3+4+17+18+19+23** | 24607 | 2403923 | 65103 | 18 min 48 sec |
| **1+2+3+4+17+18+19+23+24** | 27107 | 2997899 | 64632 | 25 min 37 sec |
| **1+2+3+4+17+18+19+23+24+25** | 29607 | 3287441 | 62456 | 26 min 59 sec |
| **CPU info: Intel(R) Xeon(R) Platinum 8352V** | | | | |

**Supplementary Tab. 5 | The value of *rad_coef* and the corresponding average amount of neighbors tested.**

| | | | | | | | | |
|---|---|---|---|---|---|---|---|---|
| **rad_coef** | 1.0 | 1.1 | 1.2 | 1.3 | 1.4 | 1.5 | 1.6 | 1.7 |
| **neighbors** | 1.00 | 3.83 | 3.83 | 3.83 | 3.83 | 7.58 | 7.58 | 7.58 |
| **rad_coef** | 1.8 | 1.9 | 2.0 | 2.1 | 2.2 | 2.3 | 2.4 | 2.5 |
| **neighbors** | 7.58 | 7.58 | 7.58 | 12.27 | 12.27 | 19.51 | 19.51 | 19.51 |

**Supplementary Tab. 6 | The value of *min_cells_rate* and the corresponding number of peaks left after filtering.**

| | | | | | | |
|---|---|---|---|---|---|---|
| **min_cells_rate** | 0.00 | 0.03 | 0.06 | 0.09 | 0.12 | 0.15 |
| **peaks count** | 99311 | 64537 | 30820 | 20222 | 15088 | 12122 |
| **min_cells_rate** | 0.18 | 0.21 | 0.24 | 0.27 | 0.30 | |
| **peaks count** | 9980 | 8377 | 7034 | 5920 | 4947 | |

**Supplementary Tab. 7 | The summary table of simulated scenario 1.** The table includes the value of $r$, the number of spots in different pre-defined domains, and the number of spots of different types across the three slices.

| **Scenario 1 ($r$=0.8)** | **Slice 0** | **Slice 1** | **Slice 2** |
|---|---|---|---|
| Number of spots in domain 0 | 300 | 300 | 300 |
| Number of Spots in domain 1 | 500 | 500 | 500 |
| Number of Spots in domain 2 | 350 | 350 | 350 |
| Number of Spots in domain 3 | 400 | 400 | 400 |
| Number of Spots in domain 4 | 450 | 450 | 450 |
| Number of spot-type 0 | 300 | 300 | 302 |
| Number of spot-type 1 | 501 | 502 | 500 |
| Number of spot-type 2 | 350 | 350 | 349 |
| Number of spot-type 3 | 400 | 397 | 399 |
| Number of spot-type 4 | 449 | 451 | 450 |

**Supplementary Tab. 8 | The summary table of simulated scenario 2.** The table includes the value of $r$, the number of spots in different pre-defined domains, and the number of spots of different types across the three slices.

| **Scenario 2 ($r$=0.7)** | **Slice 0** | **Slice 1** | **Slice 2** |
|---|---|---|---|
| Number of spots in domain 0 | 300 | 300 | 300 |
| Number of Spots in domain 1 | 500 | 500 | 500 |
| Number of Spots in domain 2 | 350 | 350 | 350 |
| Number of Spots in domain 3 | 400 | 400 | 400 |
| Number of Spots in domain 4 | 450 | 450 | 450 |
| Number of spot-type 0 | 310 | 307 | 310 |
| Number of spot-type 1 | 498 | 502 | 499 |
| Number of spot-type 2 | 354 | 357 | 352 |
| Number of spot-type 3 | 398 | 395 | 394 |
| Number of spot-type 4 | 440 | 439 | 445 |

**Supplementary Tab. 9 | The summary table of simulated scenario 3.** The table includes the value of $r$, the number of spots in different pre-defined domains, and the number of spots of different types across the three slices.

| **Scenario 3 ($r$=0.6)** | **Slice 0** | **Slice 1** | **Slice 2** |
|---|---|---|---|
| Number of spots in domain 0 | 300 | 300 | 300 |
| Number of Spots in domain 1 | 500 | 500 | 500 |
| Number of Spots in domain 2 | 350 | 350 | 350 |
| Number of Spots in domain 3 | 400 | 400 | 400 |
| Number of Spots in domain 4 | 450 | 450 | 450 |
| Number of spot-type 0 | 330 | 329 | 342 |
| Number of spot-type 1 | 513 | 507 | 501 |
| Number of spot-type 2 | 344 | 351 | 348 |
| Number of spot-type 3 | 389 | 387 | 378 |
| Number of spot-type 4 | 424 | 426 | 431 |

**Supplementary Tab. 10 | The summary table of simulated scenario 4.** The table includes the value of $r$, the number of spots in different pre-defined domains, and the number of spots of different types across the three slices.

| **Scenario 4 (*r*=0.8)** | **Slice 0** | **Slice 1** | **Slice 2** |
|---|---|---|---|
| Number of spots in domain 0 | 300 | 300 | 300 |
| Number of Spots in domain 1 | 100 | 300 | 500 |
| Number of Spots in domain 2 | 350 | 250 | 150 |
| Number of Spots in domain 3 | 400 | 300 | 200 |
| Number of Spots in domain 4 | 450 | 450 | 450 |
| Number of spot-type 0 | 300 | 300 | 302 |
| Number of spot-type 1 | 101 | 301 | 499 |
| Number of spot-type 2 | 350 | 250 | 150 |
| Number of spot-type 3 | 00 | 298 | 199 |
| Number of spot-type 4 | 449 | 451 | 450 |

**Supplementary Tab. 11 | The summary table of simulated scenario 5.** The table includes the value of $r$, the number of spots in different pre-defined domains, and the number of spots of different types across the three slices.

| **Scenario 5 (*r*=0.8)** | **Slice 0** | **Slice 1** | **Slice 2** |
|---|---|---|---|
| Number of spots in domain 0 | 300 | 300 | 300 |
| Number of Spots in domain 1 | 500 | 500 | 500 |
| Number of Spots in domain 2 | 350 | 350 | 350 |
| Number of Spots in domain 3 | 400 | 400 | 0 |
| Number of Spots in domain 4 | 450 | 450 | 450 |
| Number of spot-type 0 | 300 | 300 | 301 |
| Number of spot-type 1 | 501 | 502 | 500 |
| Number of spot-type 2 | 350 | 350 | 349 |
| Number of spot-type 3 | 400 | 397 | 0 |
| Number of spot-type 4 | 449 | 451 | 450 |

**Supplementary Tab. 12 | The summary table of simulated scenario 6.** The table includes the value of $r$, the number of spots in different pre-defined domains, and the number of spots of different types across the three slices.

| **Scenario 6 (*r*=0.8)** | **Slice 0** | **Slice 1** | **Slice 2** |
|---|---|---|---|
| Number of spots in domain 0 | 300 | 300 | 300 |
| Number of Spots in domain 1 | 25 | 0 | 0 |
| Number of Spots in domain 2 | 350 | 350 | 350 |
| Number of Spots in domain 3 | 400 | 400 | 400 |
| Number of Spots in domain 4 | 450 | 450 | 450 |
| Number of spot-type 0 | 300 | 300 | 301 |
| Number of spot-type 1 | 26 | 2 | 1 |
| Number of spot-type 2 | 350 | 350 | 349 |
| Number of spot-type 3 | 400 | 397 | 399 |
| Number of spot-type 4 | 449 | 451 | 450 |

**Supplementary Tab. 13 | The summary table of simulated scenario 7.** The table includes the value of $r$, the number of spots in different pre-defined domains, and the number of spots of different types across the three slices.

| **Scenario 7 (*r*=0.6)** | **Slice 0** | **Slice 1** | **Slice 2** |
|---|---|---|---|
| Number of spots in domain 0 | 325 | 325 | 325 |
| Number of Spots in domain 1 | 325 | 325 | 325 |
| Number of Spots in domain 2 | 400 | 400 | 400 |
| Number of Spots in domain 3 | 400 | 400 | 400 |
| Number of Spots in domain 4 | 400 | 400 | 400 |
| Number of Spots in domain 5 | 50 | 50 | 50 |
| Number of Spots in domain 6 | 50 | 50 | 50 |
| Number of Spots in domain 7 | 50 | 50 | 50 |
| Number of spot-type 0 | 333 | 331 | 334 |
| Number of spot-type 1 | 326 | 328 | 334 |
| Number of spot-type 2 | 399 | 399 | 397 |
| Number of spot-type 3 | 393 | 398 | 391 |
| Number of spot-type 4 | 391 | 385 | 384 |
| Number of spot-type 5 | 56 | 55 | 58 |
| Number of spot-type 6 | 51 | 53 | 52 |
| Number of spot-type 7 | 51 | 51 | 50 |

### Supplementary Figures

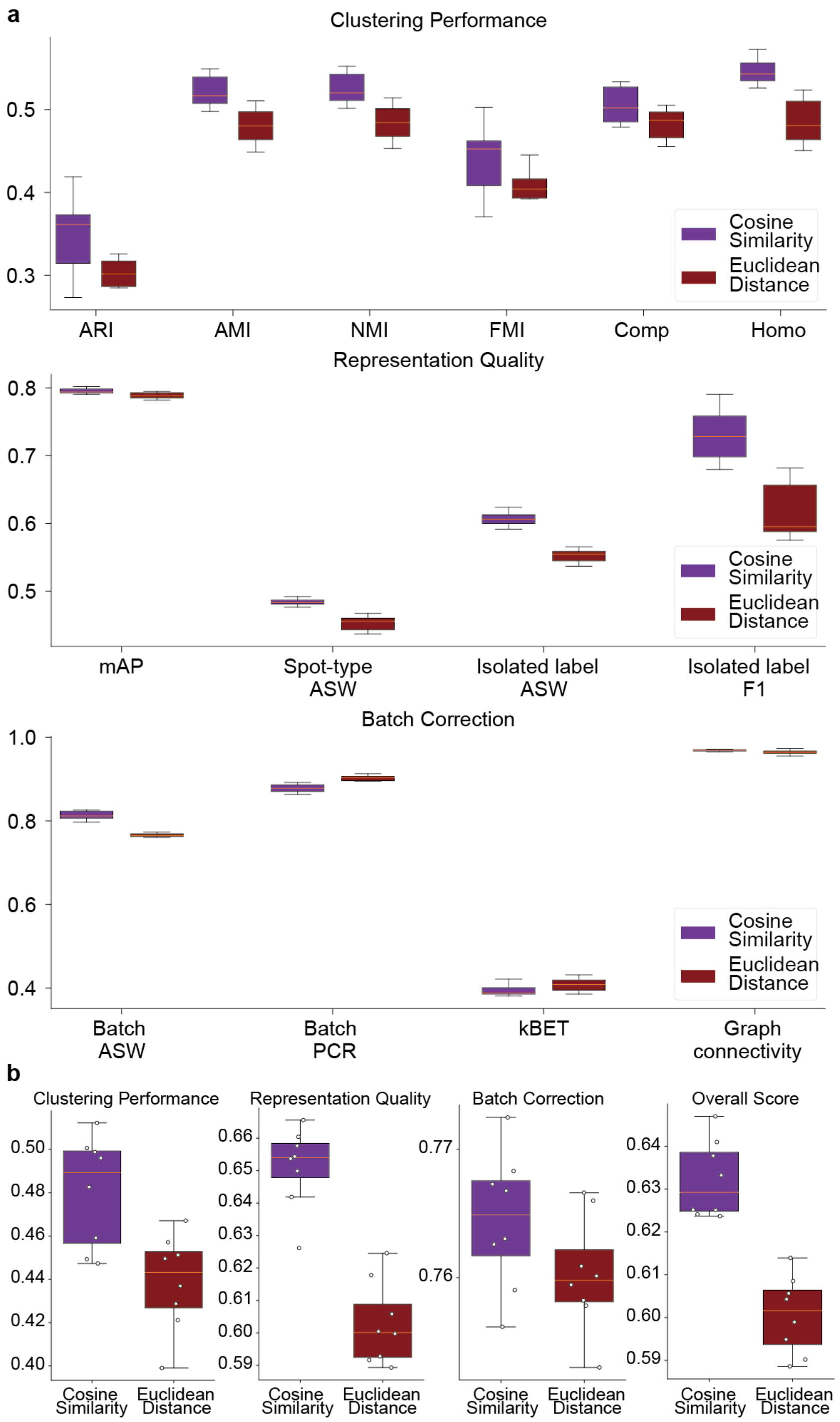

**Supplementary Fig. 1 | The comparison between using cosine similarity and Euclidean distance as the strategy in INSTINCT for selecting nearest neighbors for each spot. a,** integration of S1 slices from four developmental stages of the MISAR-seq MB dataset and comparison across 14 metrics. Using cosine similarity outperformed using Euclidean distance on 12 of the 14 metrics. **b,** the group-specific overall scores and the final overall score of different neighbor selection strategies. Employing cosine similarity consistently outperformed using Euclidean distance across all groups and in the final score.

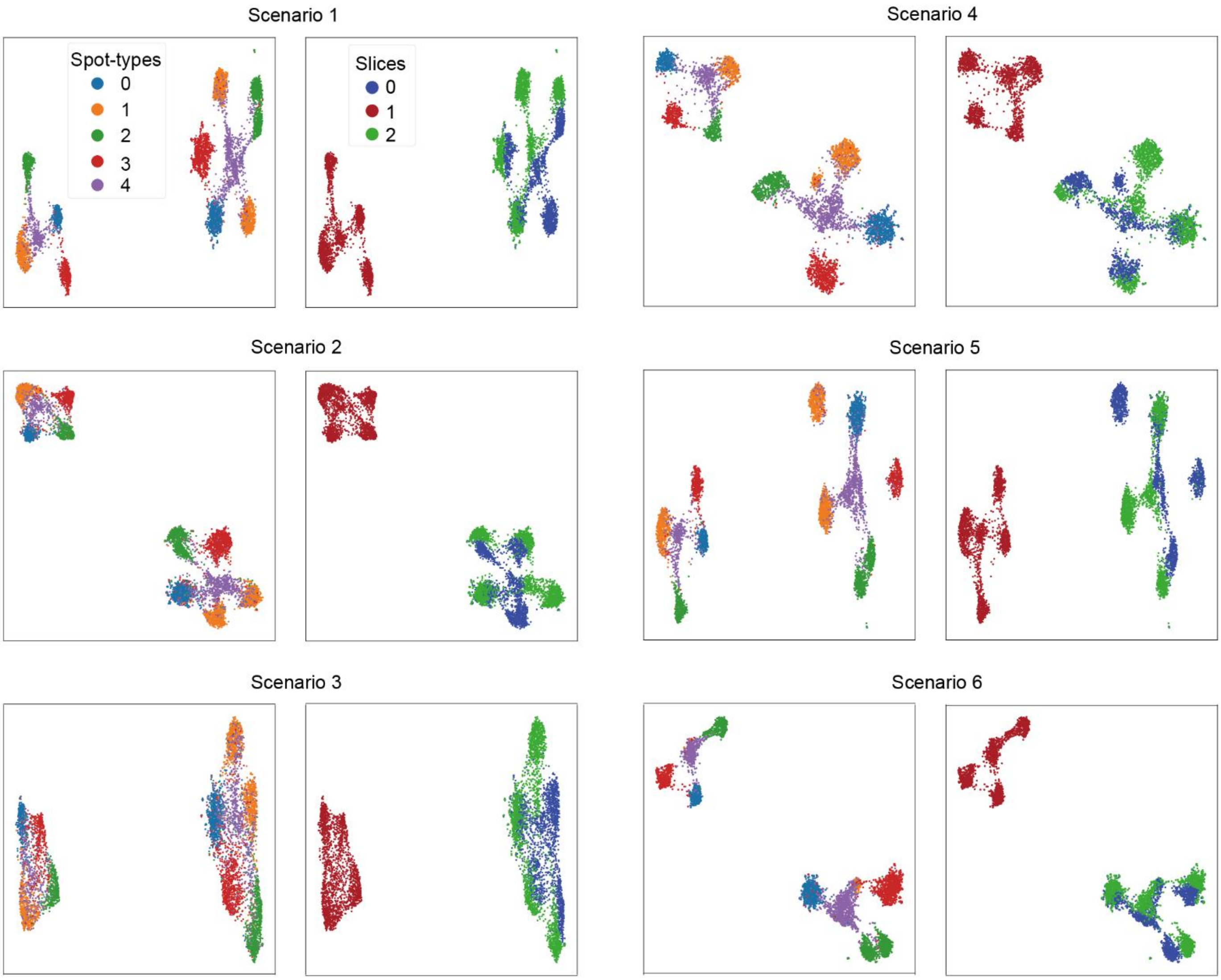

**Supplementary Fig. 2 | UMAP visualizations of simulated datasets under scenarios 1 to 6, with spots colored based on slice affiliation or spot-types.**

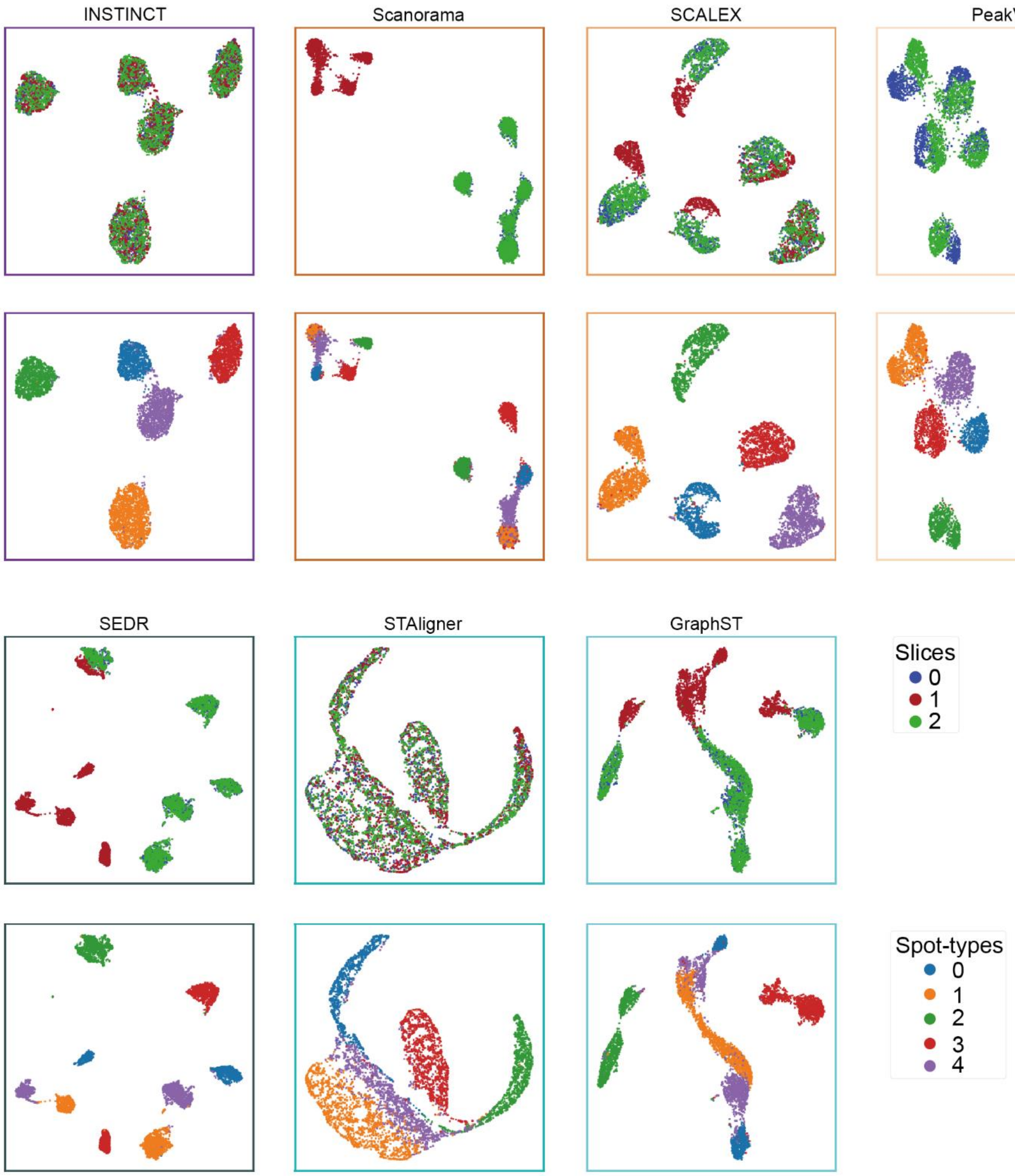

**Supplementary Fig. 3 | UMAP visualizations of the integration results of INSTINCT and six baseline methods in simulated scenario 1.** INSTINCT evenly mixed the three slices, effectively removing batch effects while preserving sufficient biological variations to distinguish between different spot-types. Most other methods only manage to mix slice 0 and 2, which were not significantly different from each other. STAligner uniformly mixed the three slices but struggles to effectively differentiate between different spot-types.

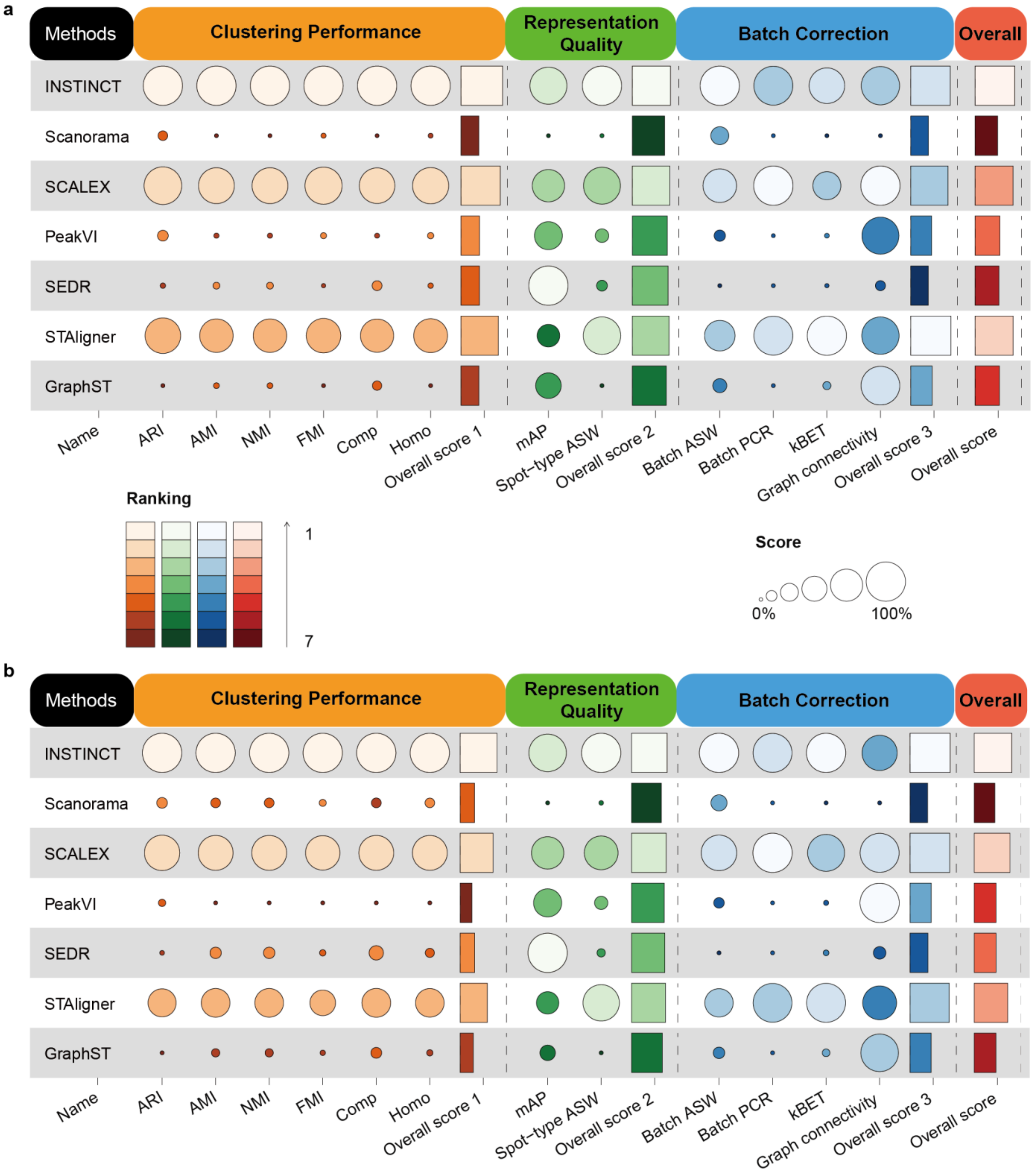

**Supplementary Fig. 4 | The overview table of metrics calculated in simulated scenario 2 and 3. a,** in scenario 2, INSTINCT ranked first in 8 of the 12 metrics and second in 2, It also achieved the best in clustering performance, representation quality, as well as the final overall score, and ranked second on batch correction. **b,** in scenario 3, INSTINCT ranked first in 9 of the 12 metrics and second in 2, It also achieved the best in clustering performance, representation quality, batch correction, as well as the final overall score.

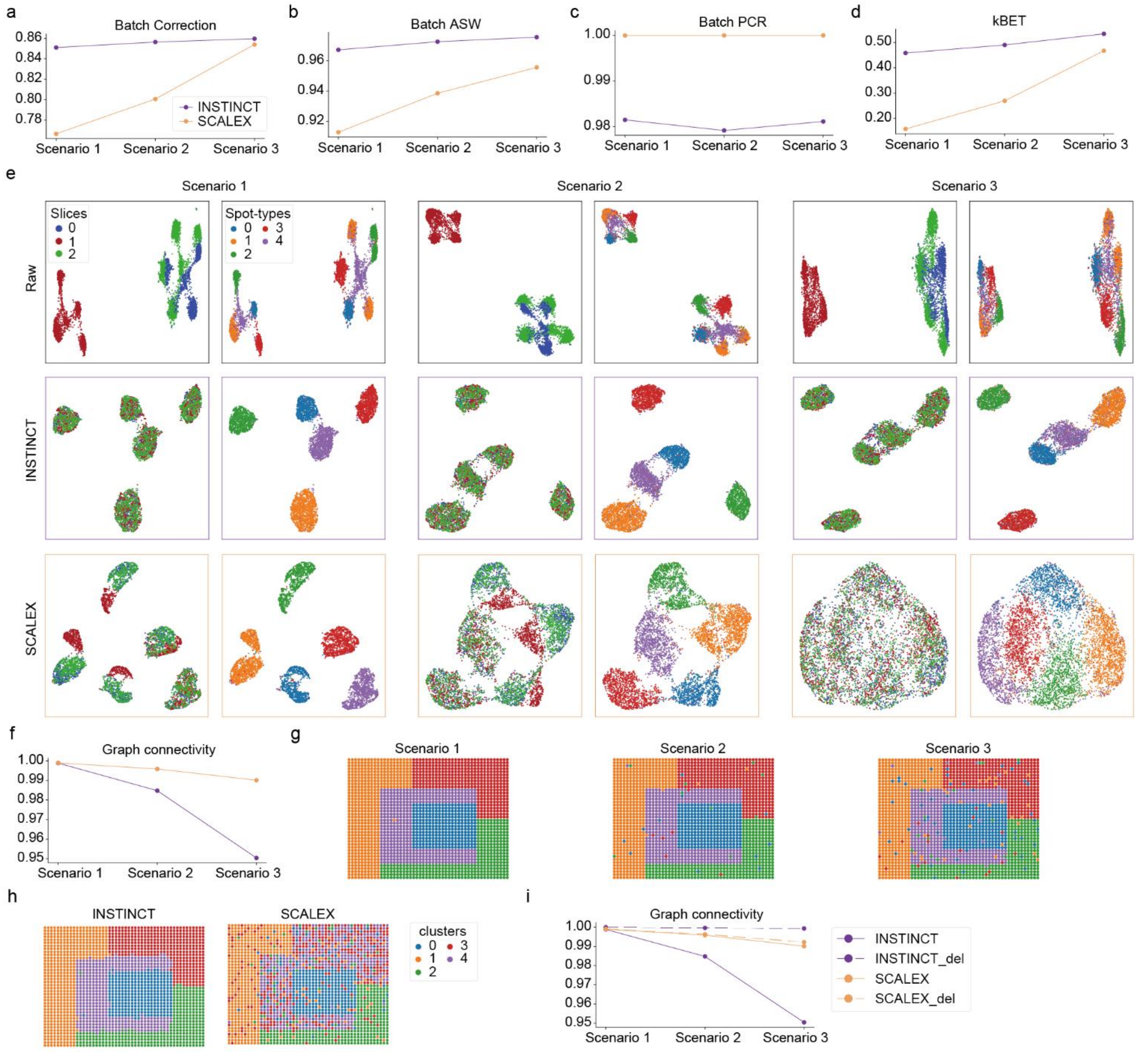

**Supplementary Fig. 5 | Comparison of batch correction performance for INSTINCT and SCALEX on simulated scenarios.** **a,** group-specific overall scores for INSTINCT and SCALEX across the three scenarios. **b,** batch ASW scores for INSTINCT and SCALEX across the three scenarios. **c,** batch PCR scores for INSTINCT and SCALEX across the three scenarios. **d,** kBET scores for INSTINCT and SCALEX across the three scenarios. **e,** UMAP visualizations of the raw data and integration results of INSTINCT and SCALEX across the three scenarios, with spots colored by slice affiliation and spot-type. **f,** graph connectivity scores for INSTINCT and SCALEX across the three scenarios. **g,** slice 0 of the three scenarios, with spots colored by spot-type. **h,** clustering results of INSTINCT and SCALEX of slice 0 in scenario 3, with spots colored by clusters. **i,** graph connectivity scores for INSTINCT and SCALEX across the three scenarios, with dashed lines representing the results after removing scattered spots.

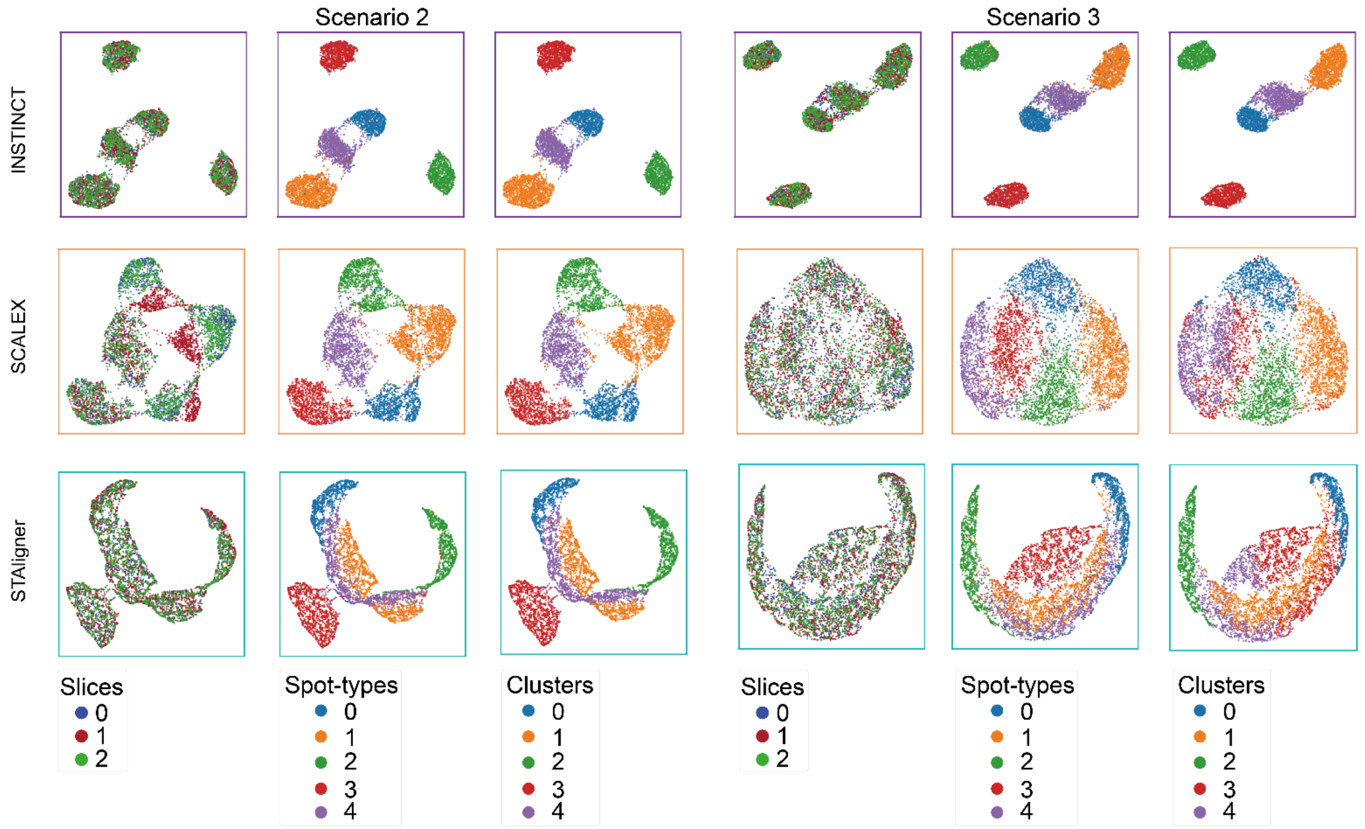

**Supplementary Fig. 6 | UMAP visualizations of the integration results of INSTINCT, SCALEX and STAligner in simulated scenario 2 and 3.** As scenarios became more complex, and different types of spots became increasingly mixed before integration, the confusion in the low-dimensional representations obtained by SCALEX for each spot also increased. Consequently, distinguishing between different spot types became more difficult. STAligner encountered the same issue across all three scenarios. In contrast, INSTINCT not only evenly mixed data from three slices but also accurately distinguished between different spot-types in the latent space across all scenarios.

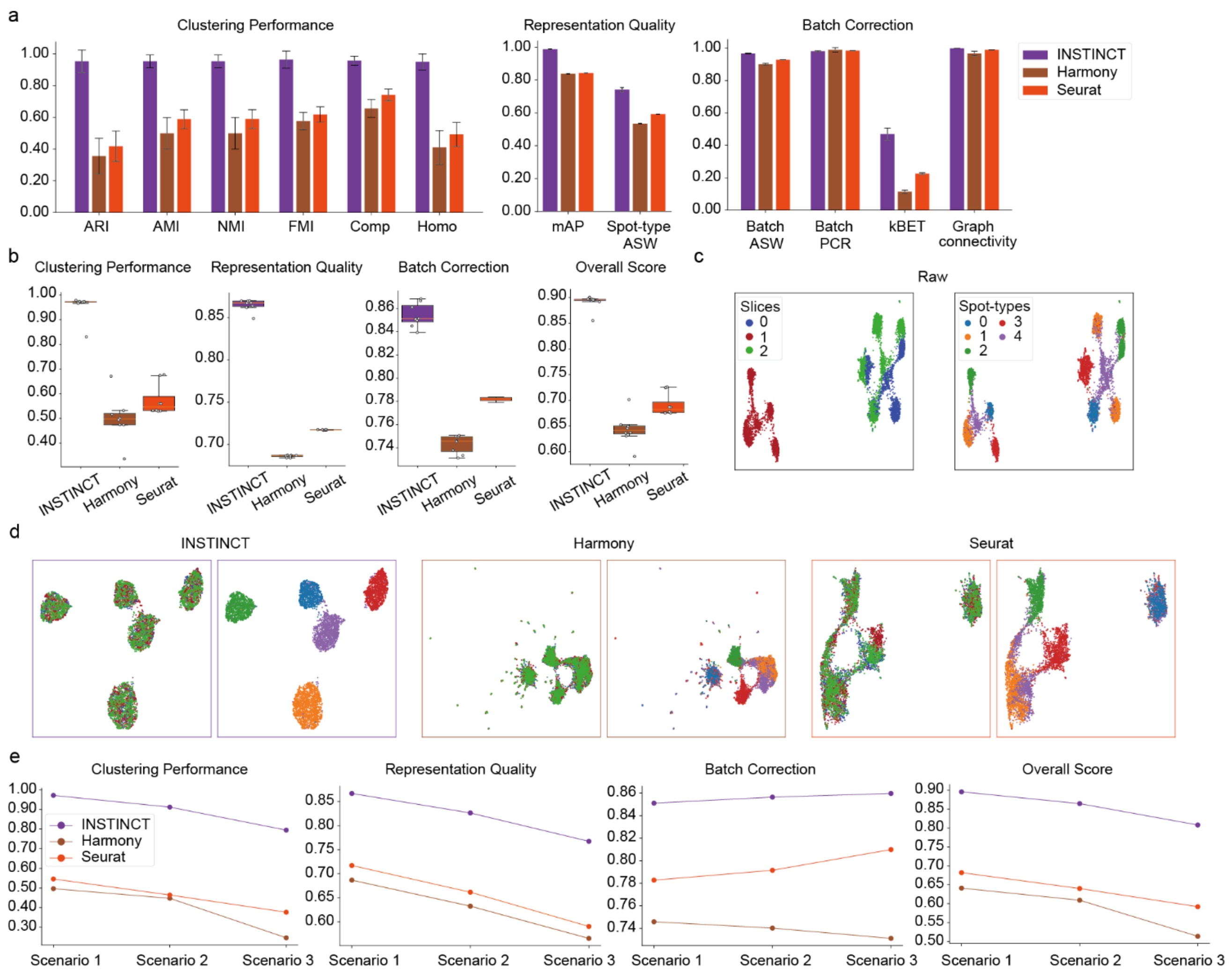

**Supplementary Fig. 7 | Comparison of INSTINCT with Harmony and Seurat on the simulated data. a,** individual scores of metrics for clustering performance, representation quality, and batch correction for the joint evaluation of all slices. **b,** group-specific overall scores and final overall scores for the joint evaluation of all slices. **c,** UMAP visualizations of the raw data from simulated scenario 1, with spots colored by slice affiliation and spot-type. **d,** UMAP visualizations for the integration results of INSTINCT, Harmony, and Seurat, with spots colored by slice affiliation and spot-type. **e,** comparison of median values the group-specific overall scores and final overall scores (median value) for INSTINCT, Harmony, and Seurat across simulated scenarios 1 to 3.

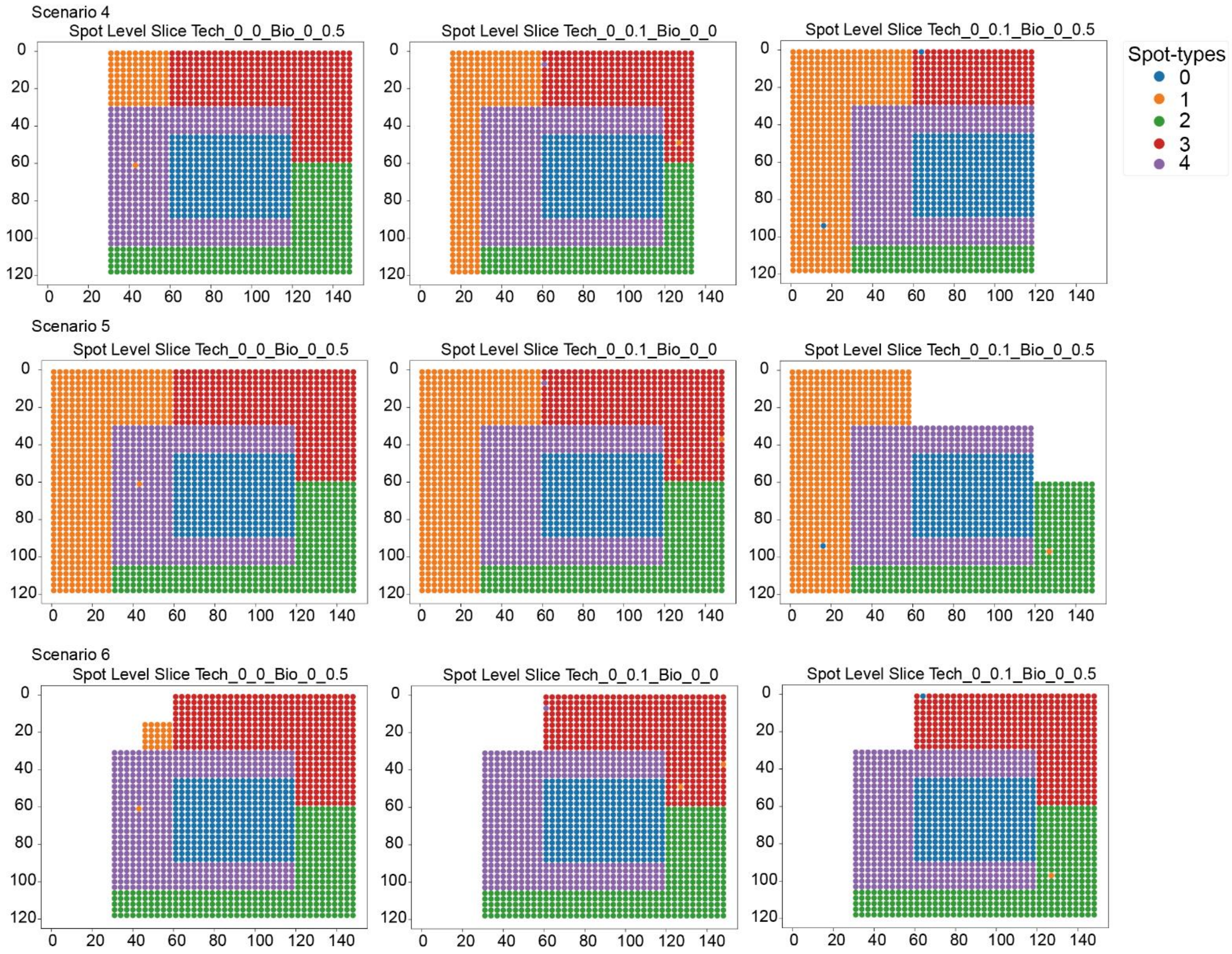

**Supplementary Fig. 8 | The three spatial resolution slices for each scenario, from scenarios 4 to 6.**

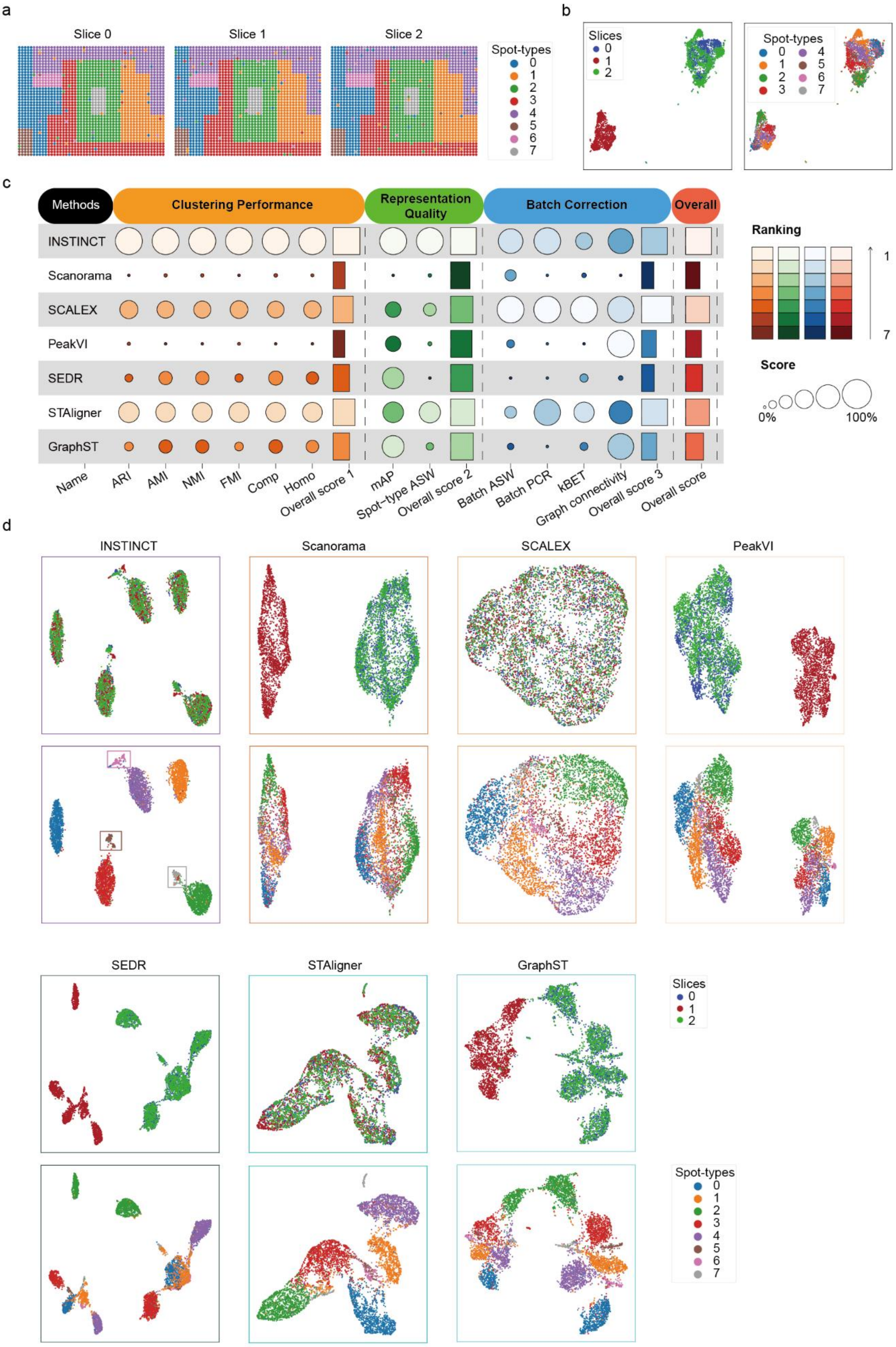

**Supplementary Fig. 9 | Comparison of INSTINCT with baseline methods on distinguishing rare spot-types. a,** the three spatial resolution slices from simulated scenario 7. **b,** UMAP visualizations of the raw data from simulated scenario 7, with spots colored based on slice affiliation and spot-type. **c,** summary table of integration performance for INSTINCT and baseline methods on simulated scenario 7, including scores of individual metrics for clustering performance, representation quality, and batch correction, as well as group-specific overall scores and final overall scores. **d,** UMAP visualizations of integration results for INSTINCT and baseline methods on simulated scenario 7, with spots colored based on slice affiliation and spot-type.

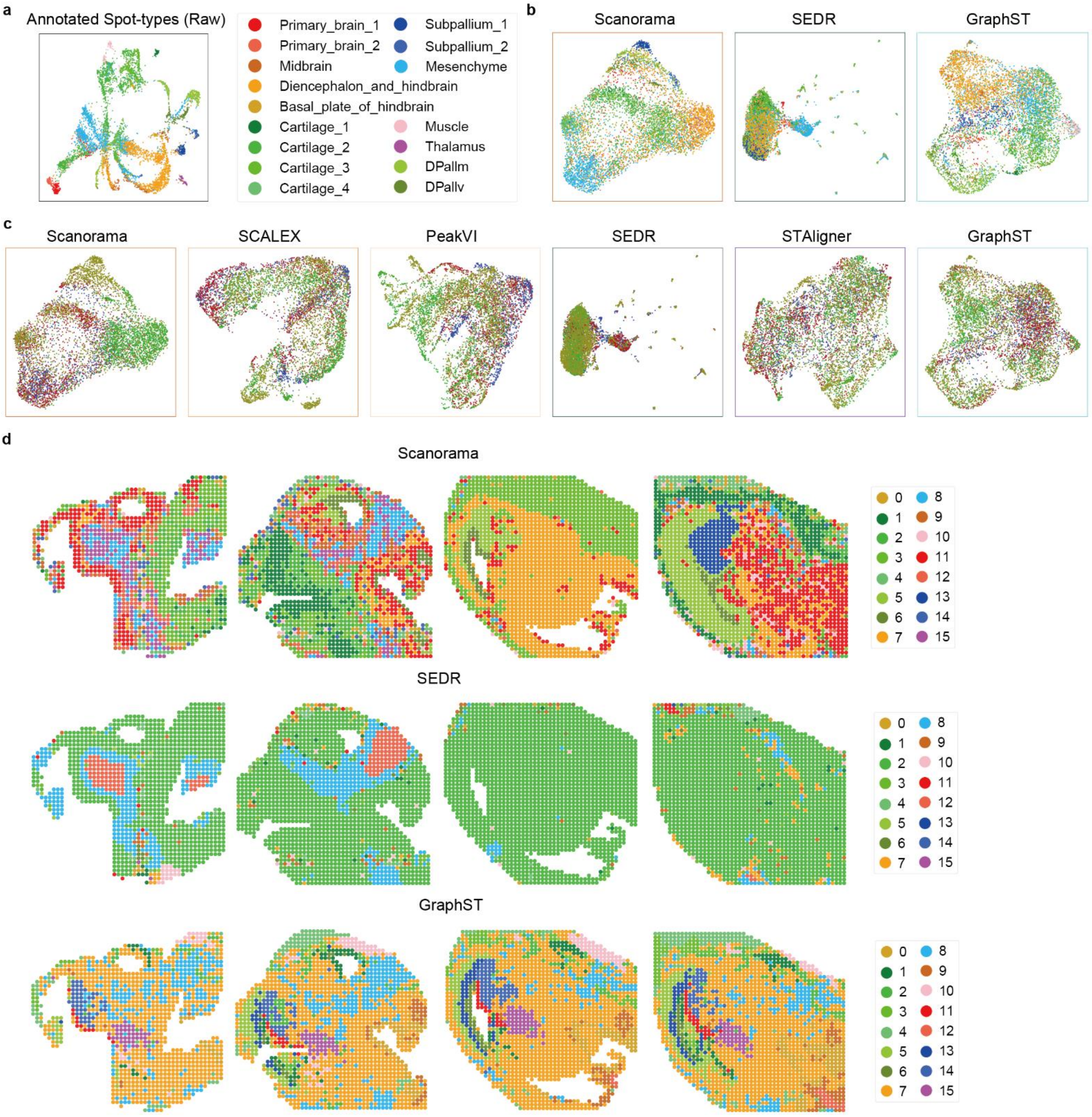

**Supplementary Fig. 10 | Comparison of INSTINCT with baseline methods on MISAR-seq MB dataset. a,** UMAP visualizations of integration results for the four methods that rank in the top for the final overall score, with spots colored based on spot-type. **b,** UMAP visualizations of integration results for the four methods that rank in the top for the final overall score, with spots colored based on slice affiliation. **c,** four S1 slices of the MISAR-seq MB dataset, with spots colored based on the provided annotations. **d,** clustering results of the four methods that rank in the top for the final overall score, with identified clusters first mapped to true labels and then colored accordingly.

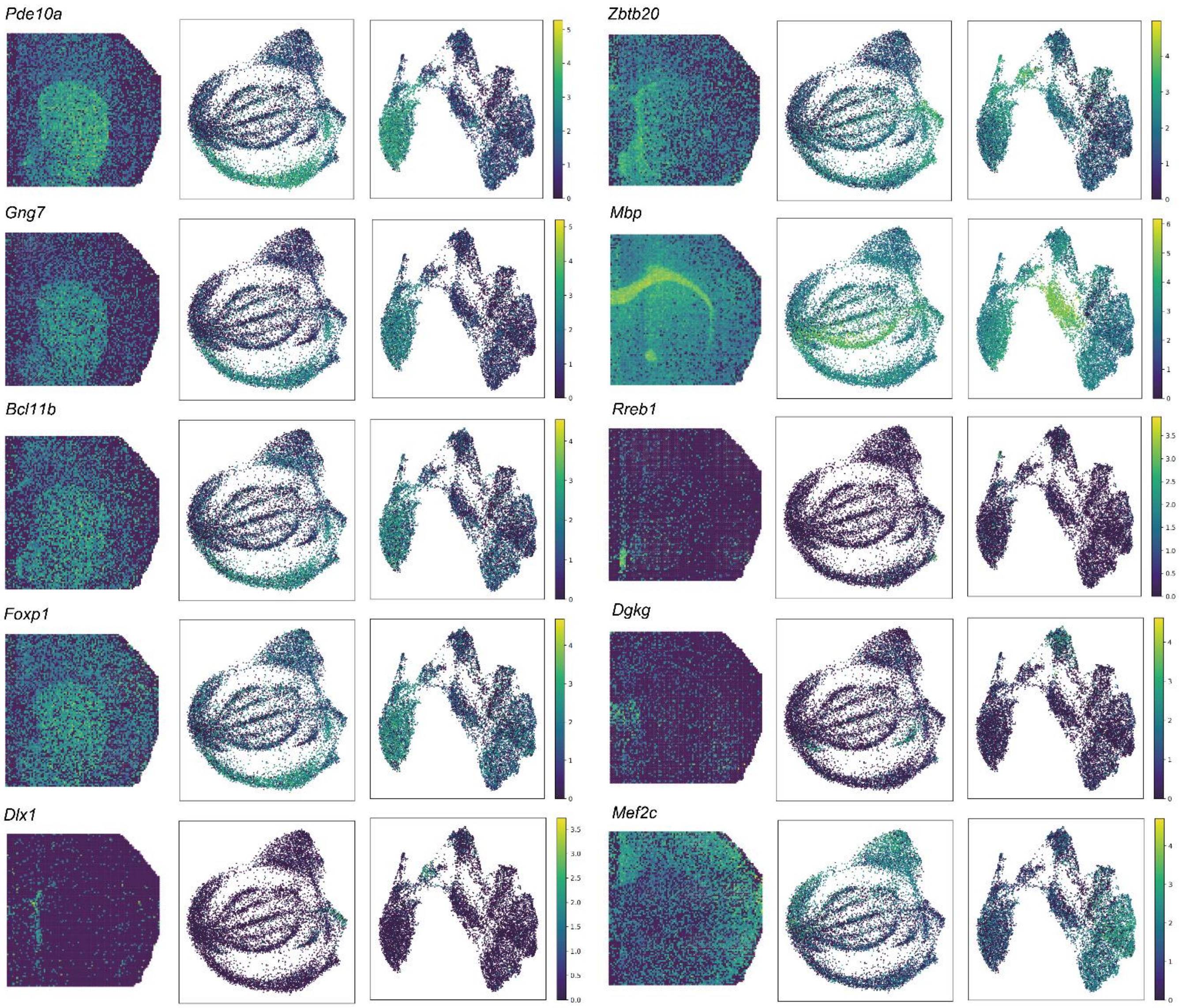

**Supplementary Fig. 11 | The marker genes of several spatial domains, with spots colored based on their expression levels.** The spatial organization of slice 2 ($100 \times 100$ barcodes, left), UMAP visualization of data before integration (middle), and UMAP visualization of the integration results (right) are shown.

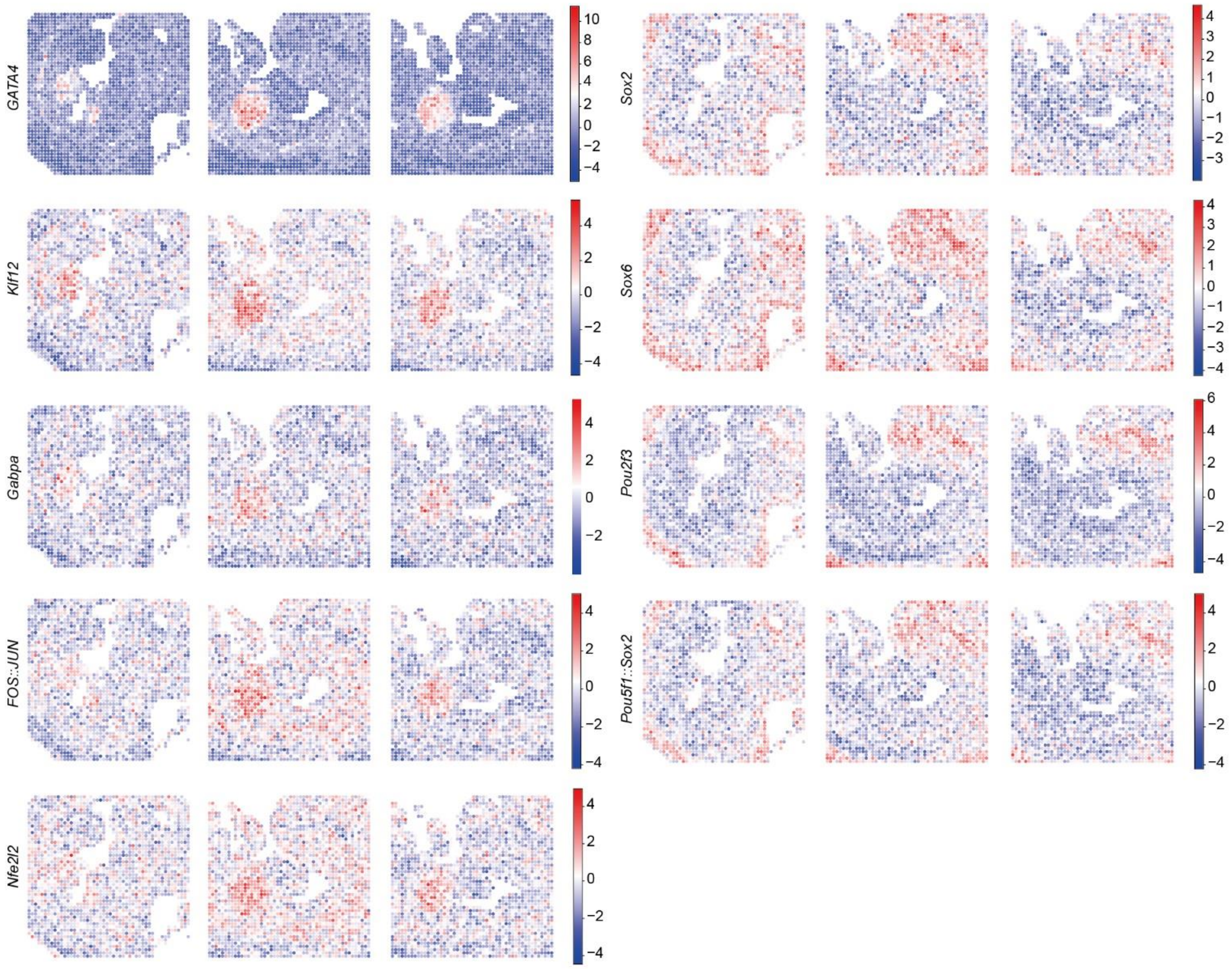

**Supplementary Fig. 12 | Motif enrichment analysis results for the three slices from the spatial-ATAC-seq ME dataset.** spots are colored based on the motif enrichment score with respect to specific motifs.

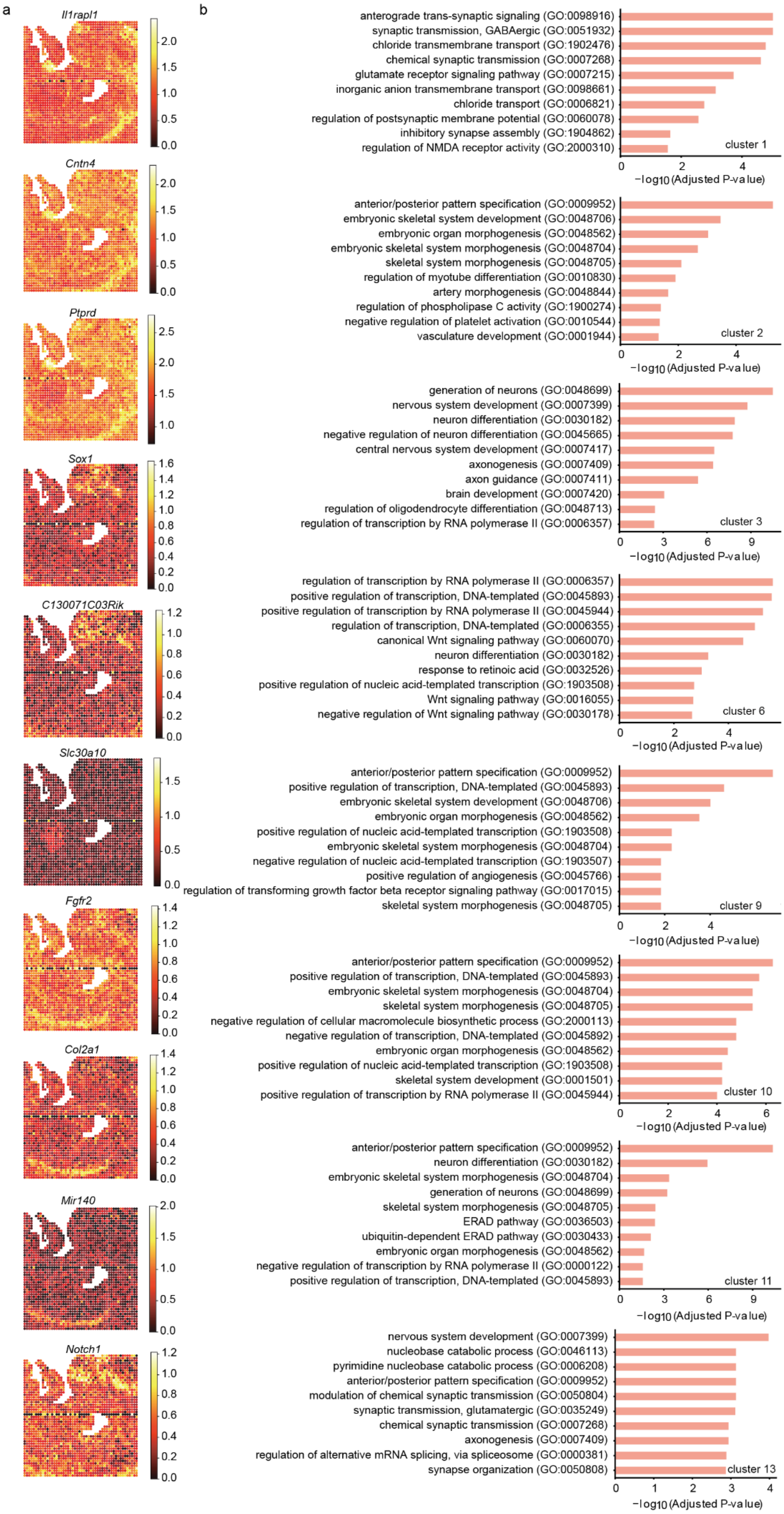

**Supplementary Fig. 13 | Analyses based on the gene score matrix of ME13_1 from the spatial-ATAC-seq ME dataset. a,** spatial organization of the DEGs. **b,** GO analysis performed based on the DEGs.

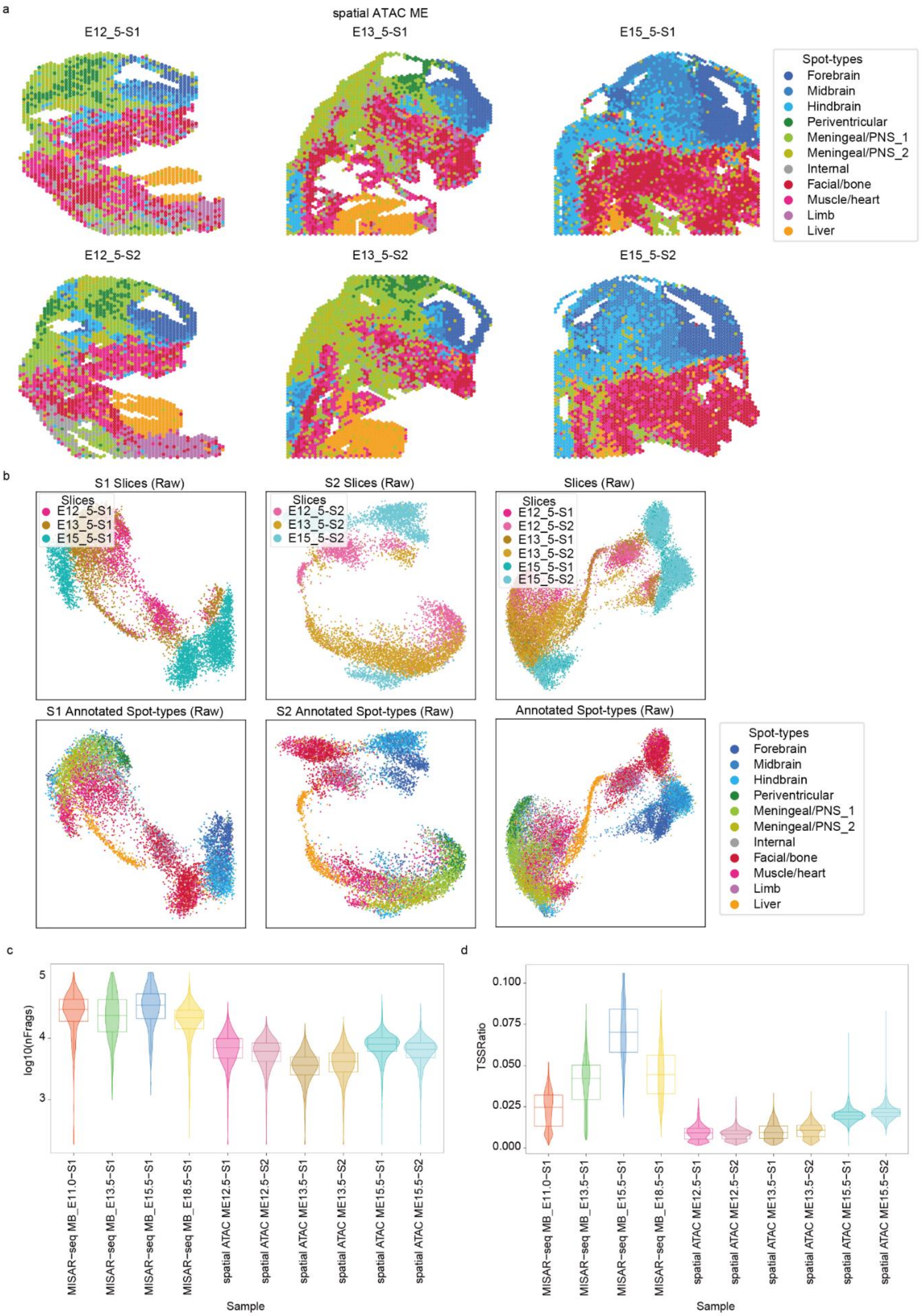

**Supplementary Fig. 14 | The spatial ATAC ME dataset. a,** six slices from spatial ATAC ME dataset with spots colored based on the provided annotations. **b,** the UMAP visualizations for three S1 slices (left), three S2 slices (middle), and all six slices (right) are shown. **c,** comparison of the log-normalized fragment counts across the four S1 samples from the MISAR-seq MB dataset and the six samples from the spatial ATAC ME dataset. **d,** comparison of the TSS ratios across the four S1 samples from the MISAR-seq MB dataset and the six samples from the spatial ATAC ME dataset.

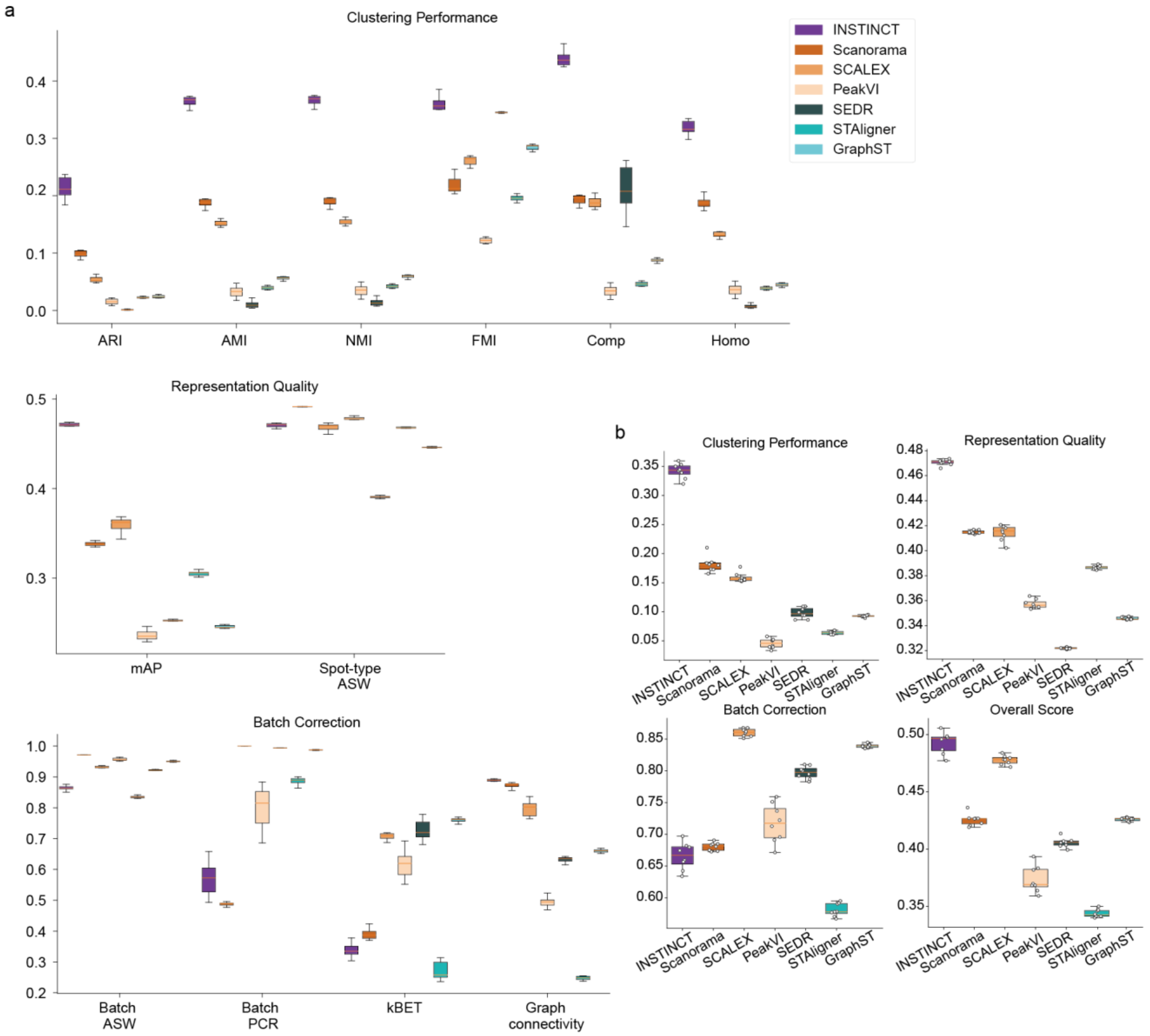

**Supplementary Fig. 15 | Comparison of INSTICNT with baseline methods in integrating three S1 slices from spatial ATAC ME dataset. a,** scores for integrating the three S1 slices of the spatial ATAC ME dataset using INSTINCT and baseline methods across 12 metrics. INSTINCT ranked first on 8 of the 12 metrics. **b,** the group-specific overall scores and the final overall score. INSTINCT ranked first in clustering performance, representation quality, as well as the final overall score.

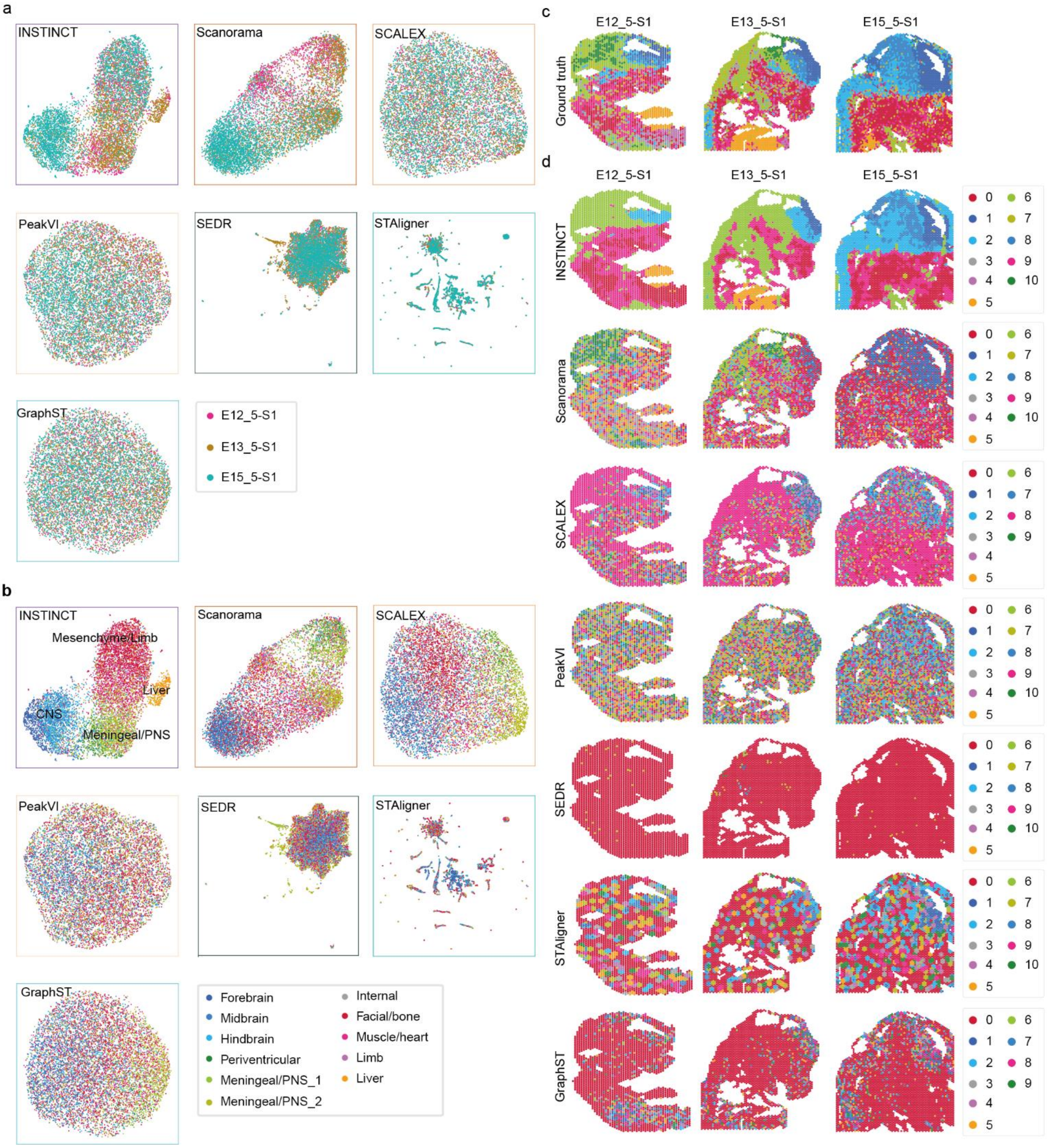

**Supplementary Fig. 16 | Integration results of INSTINCT and baseline methods in integrating three S1 slices from spatial ATAC ME dataset. a,** UMAP visualization of integration results, with spots colored according to slice affiliation. **b, U**MAP visualization of integration results, with spots colored according to the provided annotations. **c,** the provided annotations of different slices. **d,** clustering results with identified clusters first mapped to true labels and then colored accordingly.

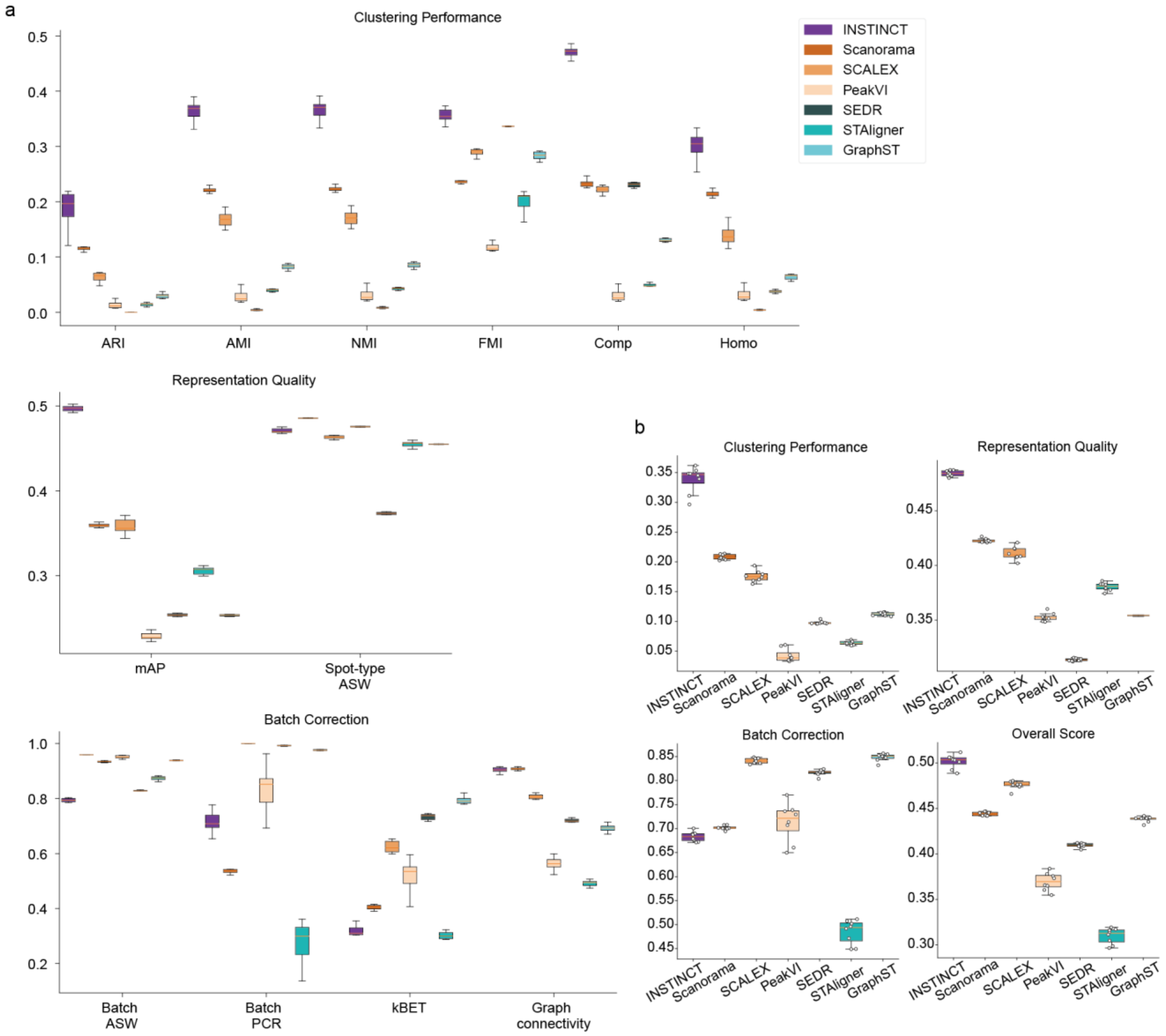

**Supplementary Fig. 17 | Comparison of INSTICNT with baseline methods in integrating three S2 slices from spatial ATAC ME dataset. a,** scores for integrating the three S2 slices of the spatial ATAC ME dataset using INSTINCT and baseline methods across 12 metrics. INSTINCT ranked first on 7 of the 12 metrics and second on 1. **b,** the group-specific overall scores and the final overall score. INSTINCT ranked first in clustering performance, representation quality, as well as the final overall score.

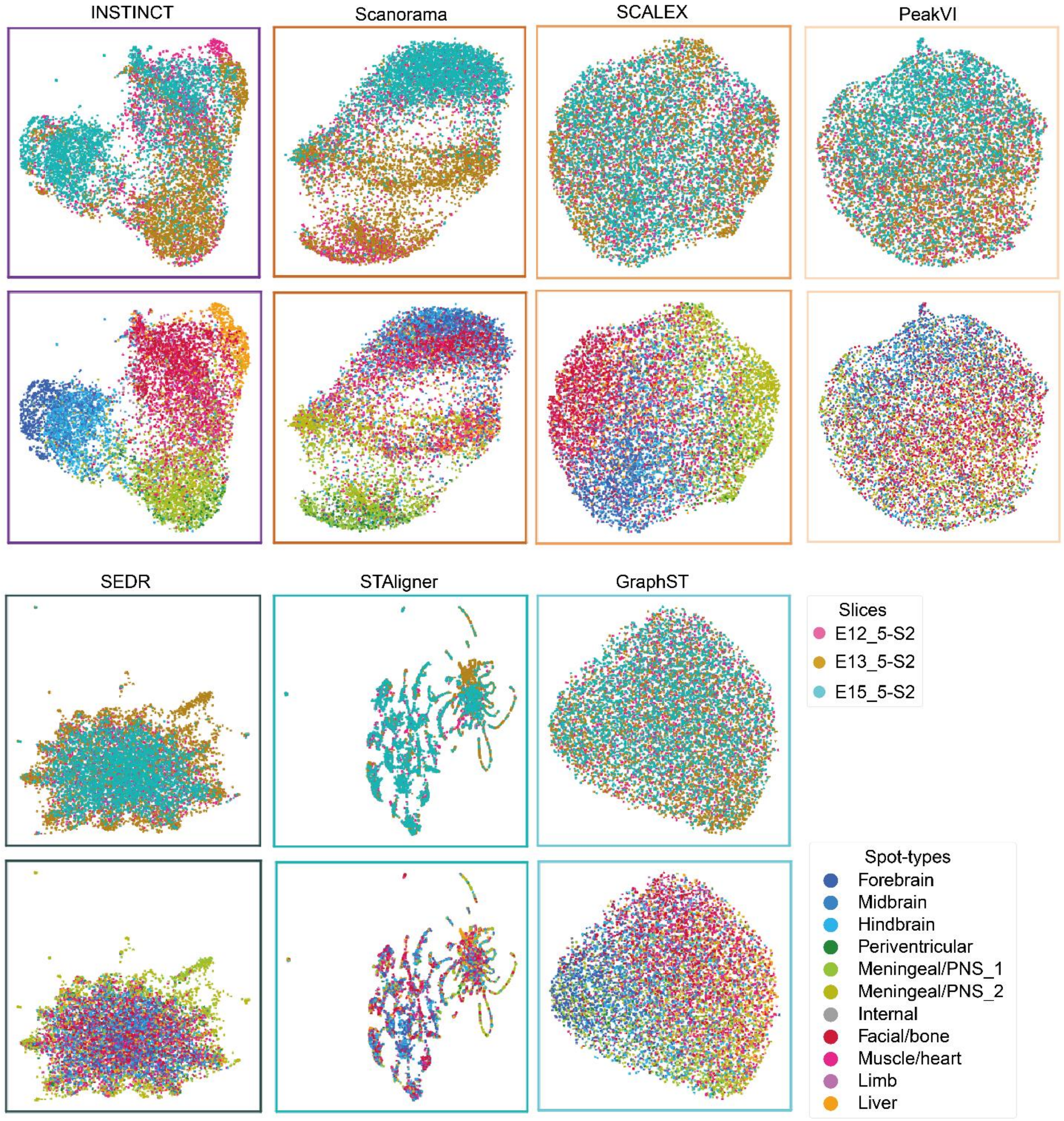

**Supplementary Fig. 18 | UMAP visualizations of the integration results of the three S2 slices from the spatial ATAC ME dataset.** Spots are colored based on slice affiliation or the provided annotations.

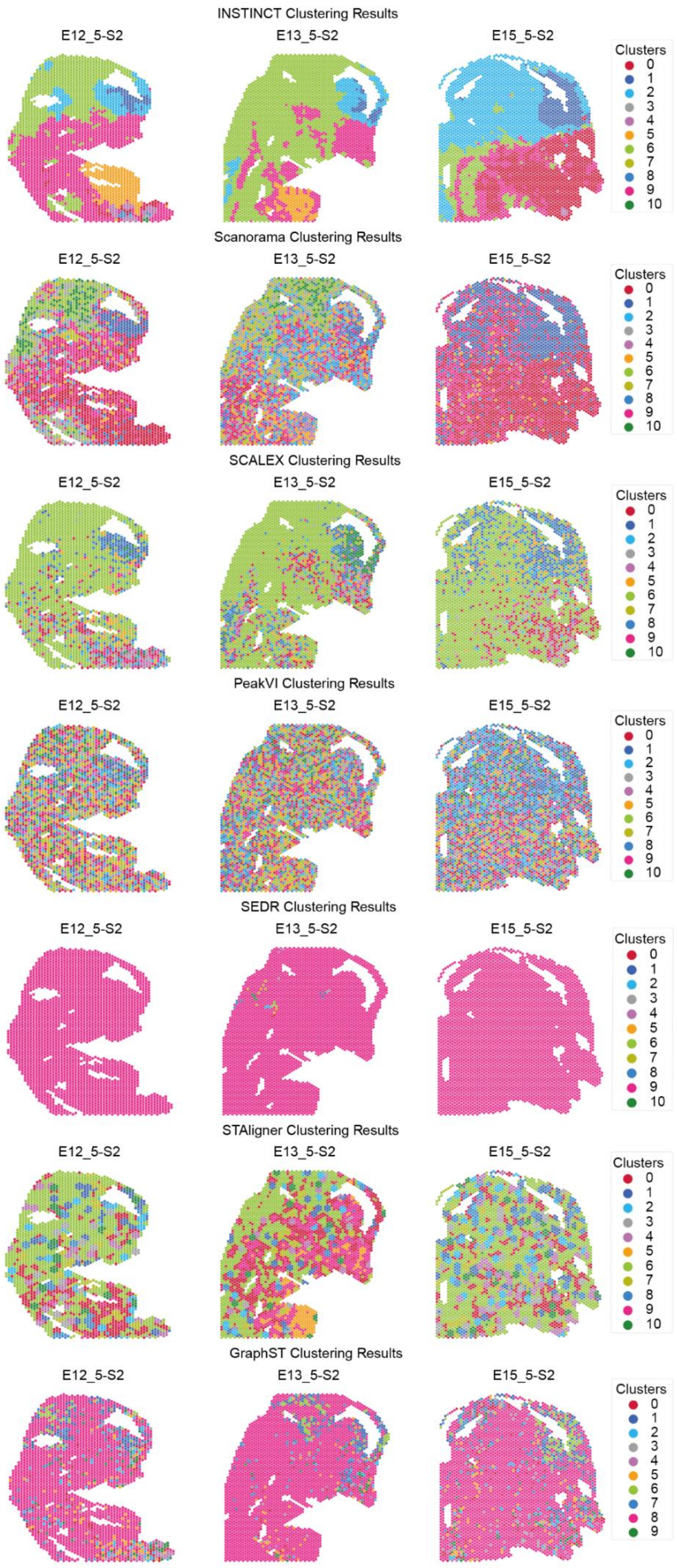

**Supplementary Fig. 19 | The clustering results of the three S2 slices from the spatial ATAC ME dataset.** The clusters were first mapped to the true spot-types and then colored accordingly.

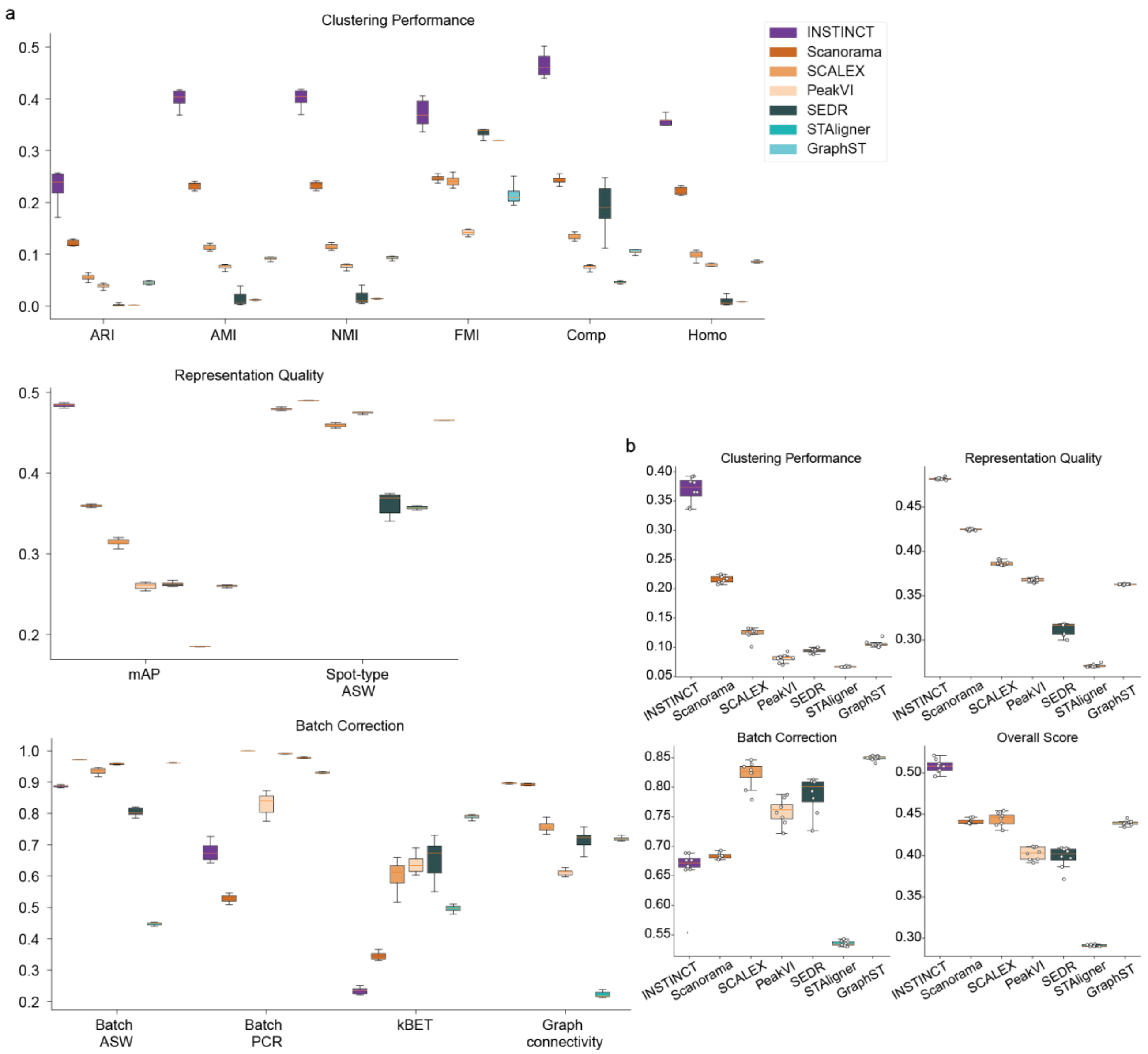

**Supplementary Fig. 20 | Comparison of INSTICNT with baseline methods in integrating all six slices from spatial ATAC ME dataset. a,** scores for integrating all six slices of the spatial ATAC ME dataset using INSTINCT and baseline methods across 12 metrics. INSTINCT ranked first on 8 of the 12 metrics and second on 1. **b,** the group-specific overall scores and the final overall score. INSTINCT ranked first in clustering performance, representation quality, as well as the final overall score.

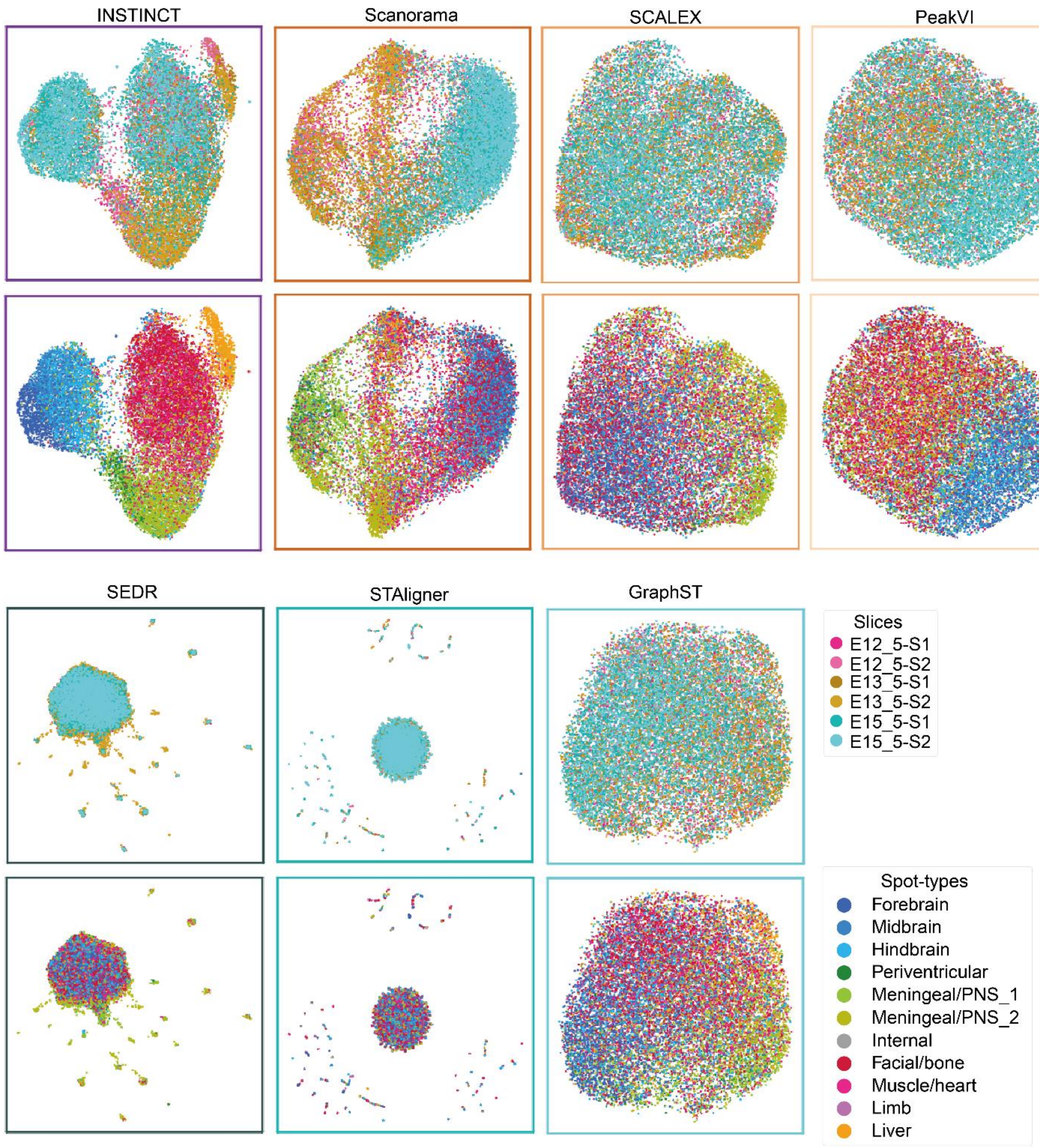

**Supplementary Fig. 21 | UMAP visualizations of the integration results of all six slices from the spatial ATAC ME dataset.** Spots are colored based on slice affiliation or the provided annotations.

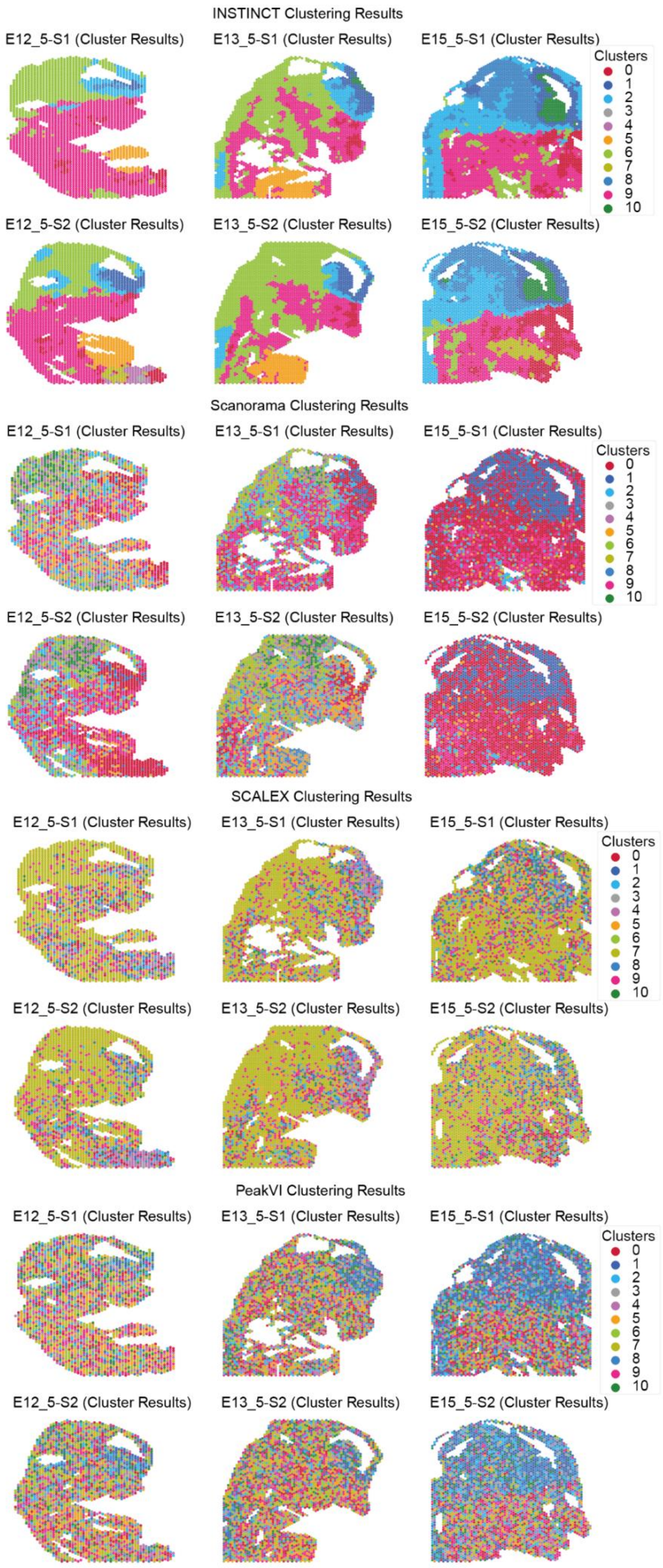

**Supplementary Fig. 22 | The clustering results of all six slices from the spatial ATAC ME dataset (INSTINCT, Scanorama, SCALEX, PeakVI).** The clusters were first mapped to the true spot-types and then colored accordingly.

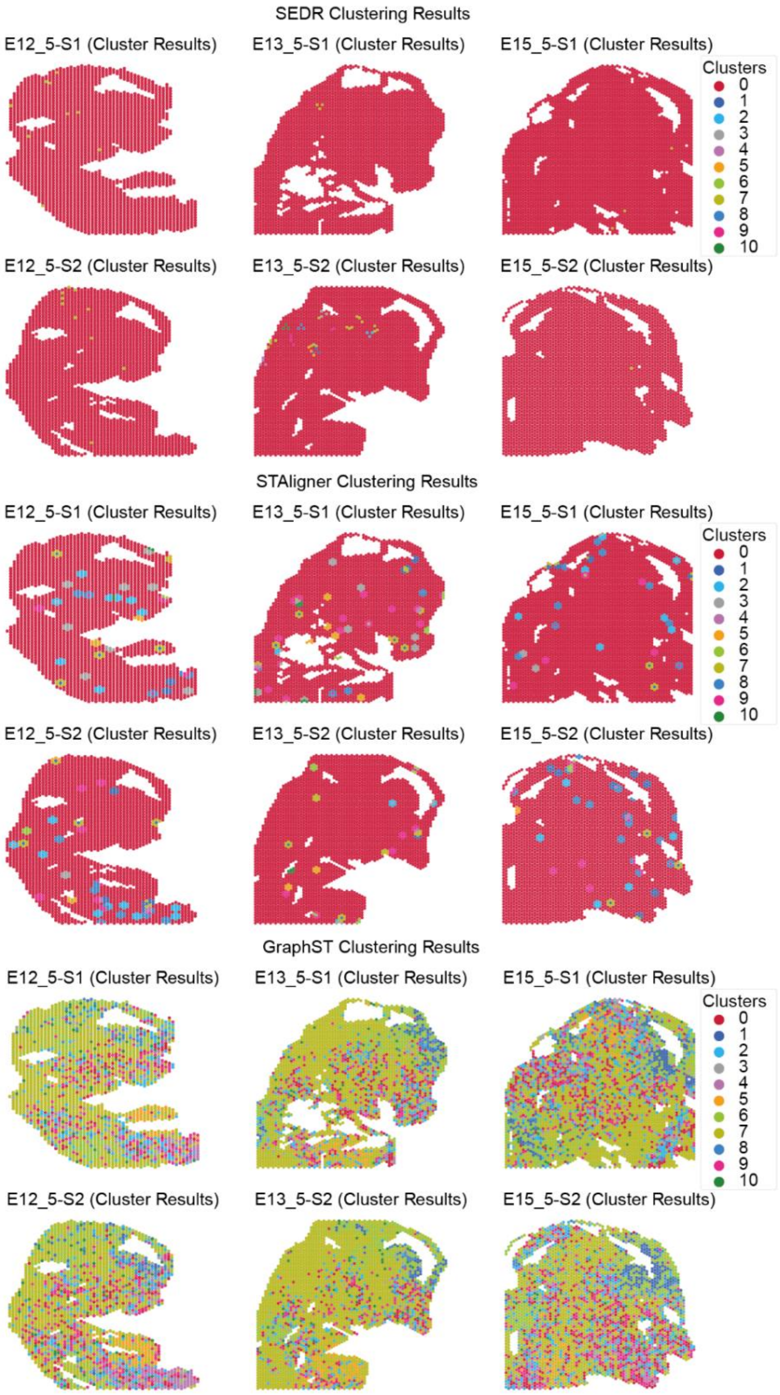

**Supplementary Fig. 23 | The clustering results of all six slices from the spatial ATAC ME dataset (SEDR, STAligner, GraphST).** The clusters were first mapped to the true spot-types and then colored accordingly.

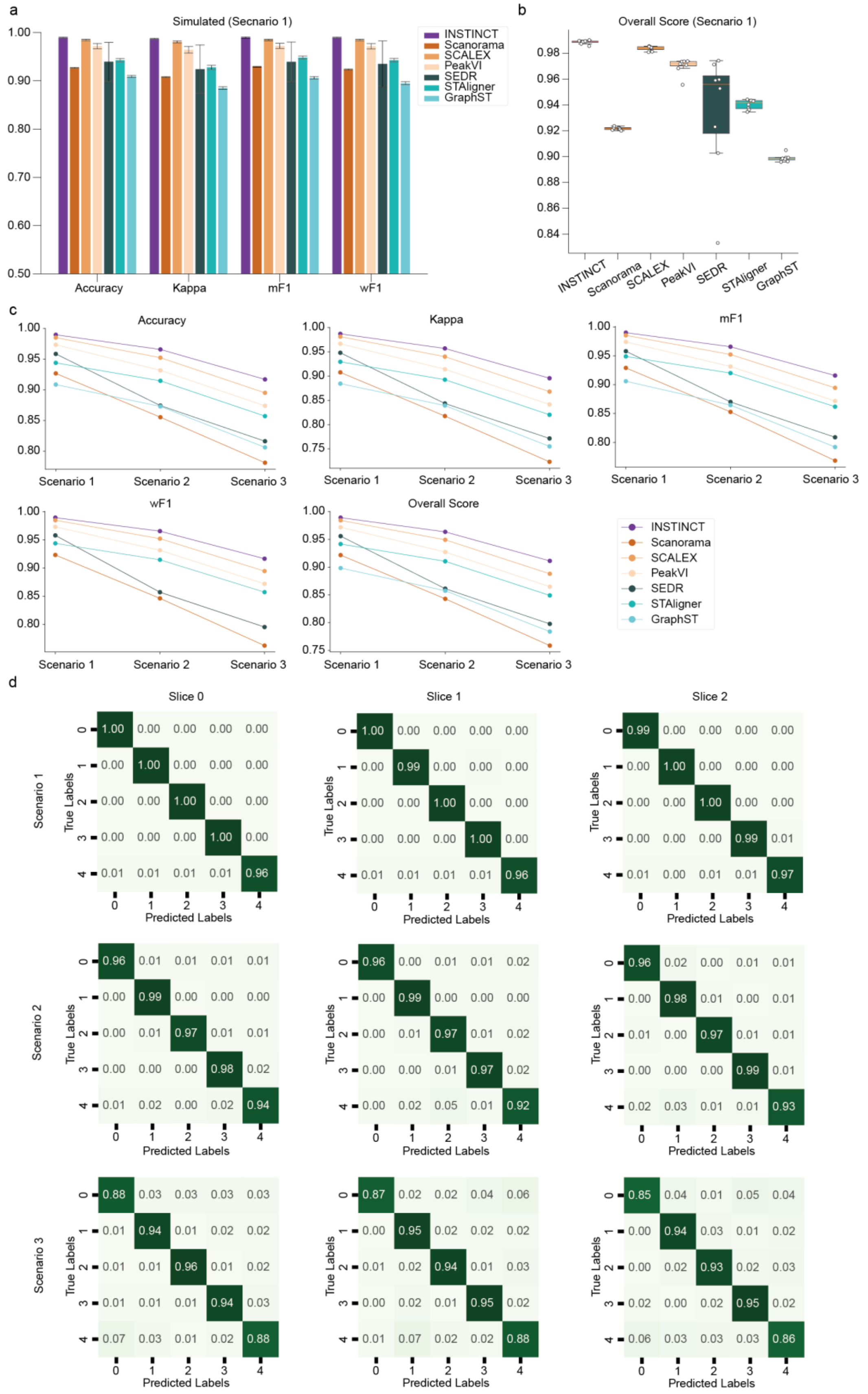

**Supplementary Fig. 24 | Comparison of annotation results of INSTINCT and baseline methods on the simulated data. a,** comparison on four metrics: accuracy, kappa, mF1, and wF1, for evaluating annotation results using cross-validation on simulated data from scenario 1. **b,** overall score of annotation results on simulated data from scenario 1. **c,** comparison of the individual scores and overall score across simulated scenarios 1 to 3. **d,** confusion matrices of the annotation results provided by INSTINCT for each slice across simulated scenarios 1 to 3.

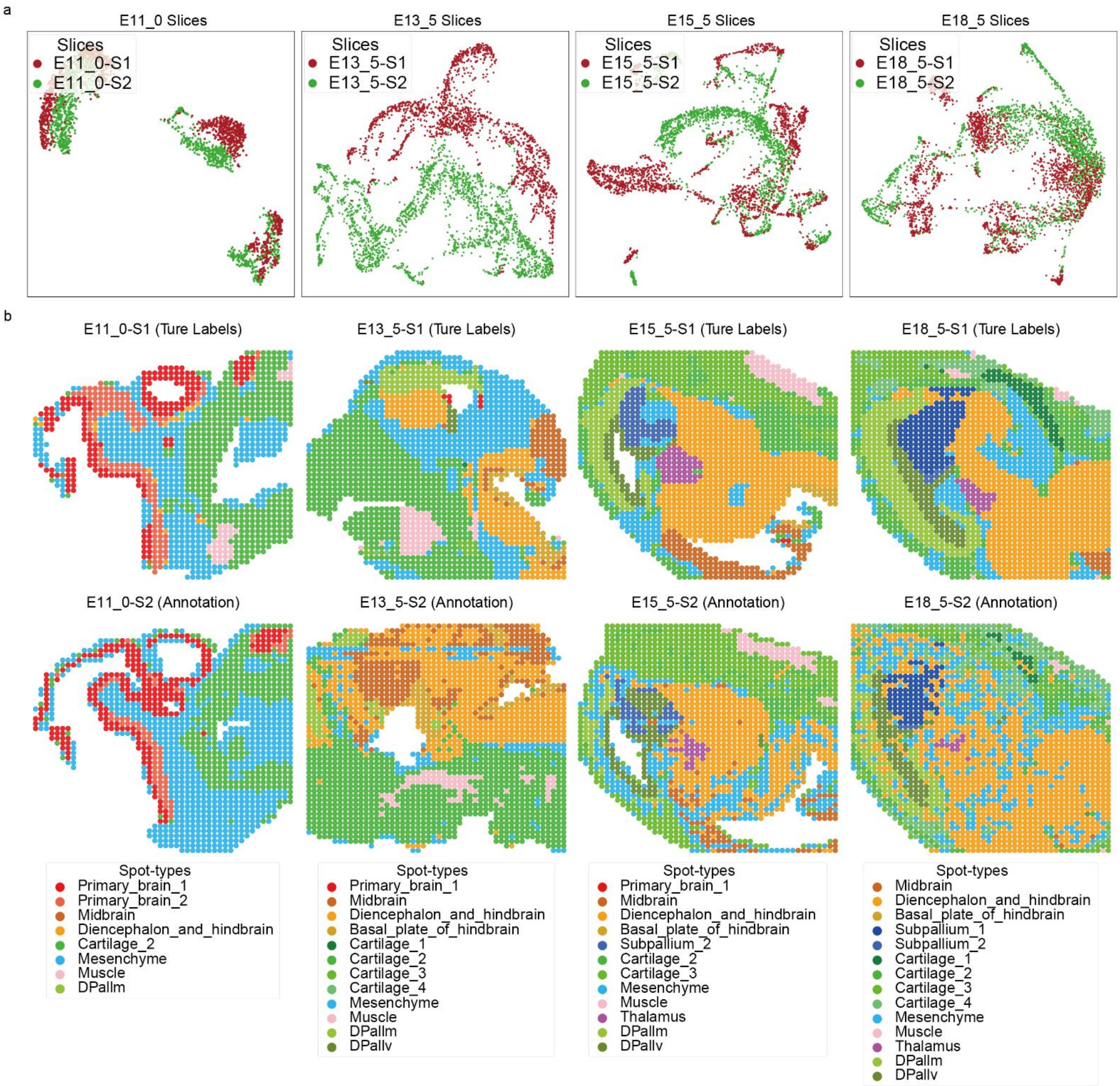

**Supplementary Fig. 25 | Annotation results on the MISAR-seq dataset based on PCA. a,** UMAP visualizations of data from each developmental stages before integration. Evident batch effects can be observed in each pair of slices. **b,** the annotation results of PCA. In slices from the more structurally complex tissues of E15.5 and E18.5, PCA annotations exhibit weak spatial continuity, making it challenging to identify clear spatial domains. Additionally, in the S2 slice of E13.5, the proportion of mesenchyme is significantly smaller compared to the corresponding S1 slice.

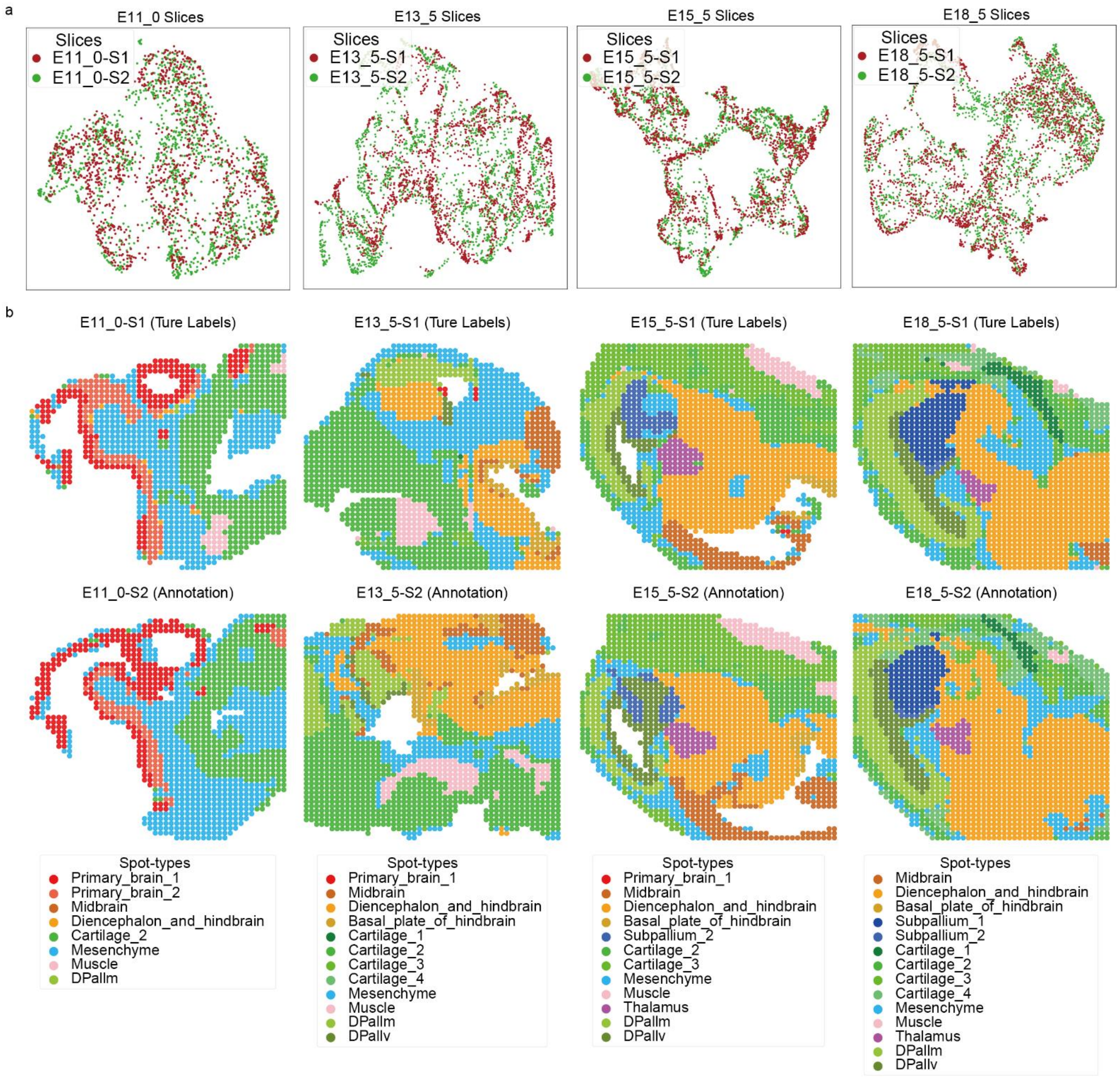

**Supplementary Fig. 26 | Annotation results on the MISAR-seq dataset based on INSTINCT. a,** UMAP visualizations of stage-specific integration results of INSTINCT. Spots from different slices were mixed and batch effects were eliminated. **b,** the annotation results of INSTINCT. Annotations of S2 slices shared similar structure with corresponding S1 slices.

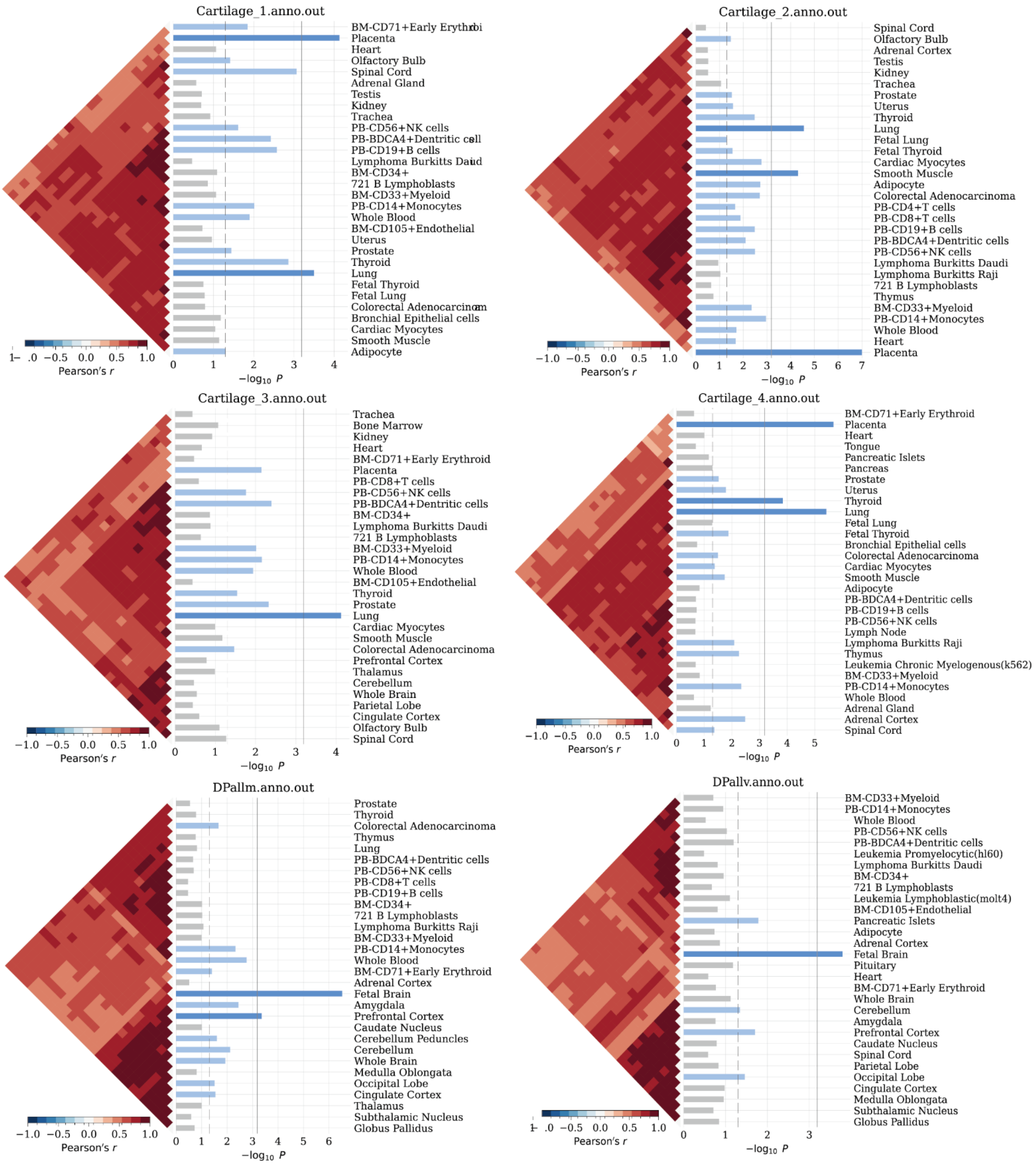

**Supplementary Fig. 27 | The results of expression enrichment analysis (cartilage_1, cartilage_2, cartilage_3, cartilage_4, DPallm, and DPallv).** The top 30 enriched tissues are displayed.

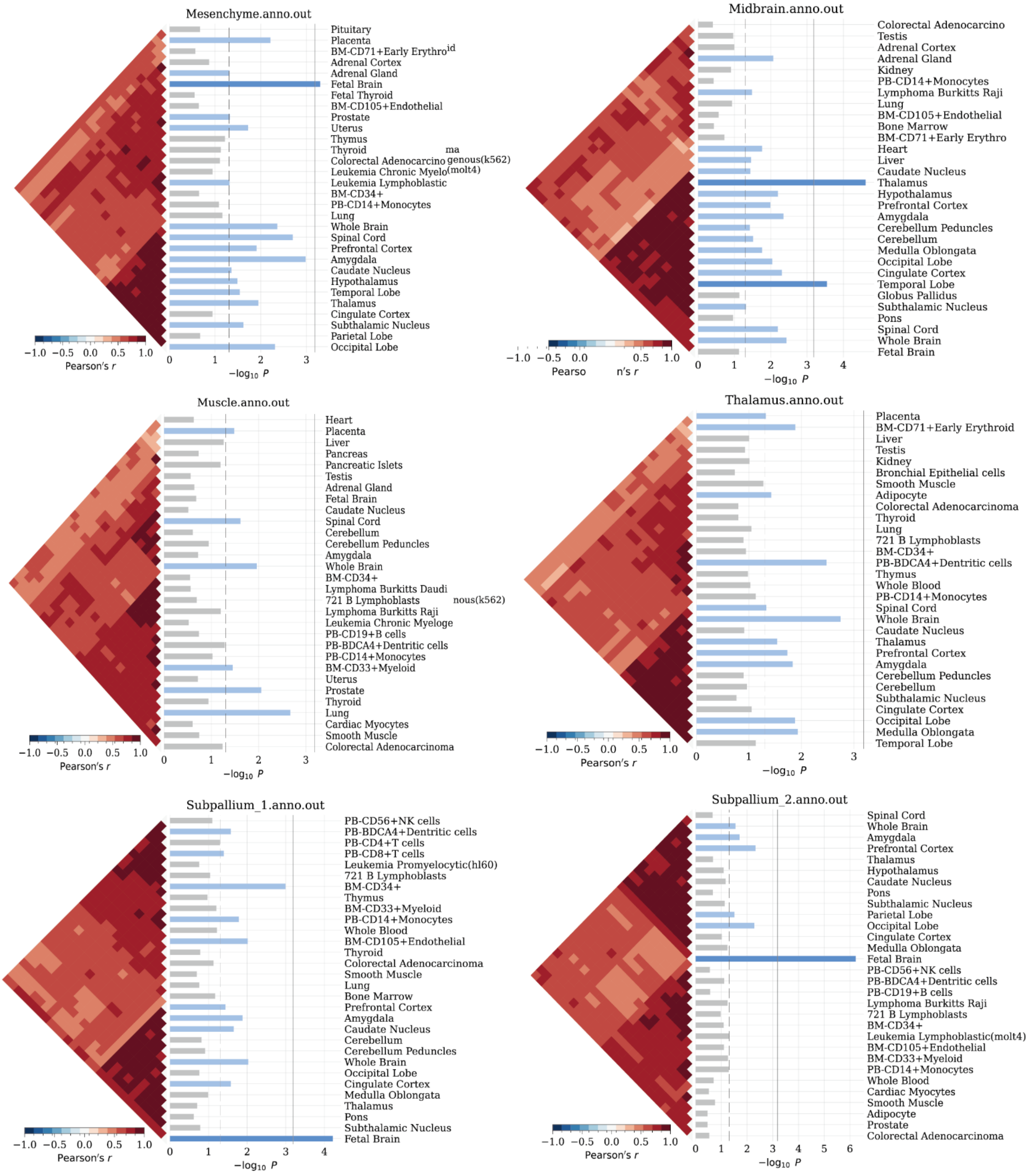

**Supplementary Fig. 28 | The results of expression enrichment analysis (mesenchyme, midbrain, muscle, thalamus, subpallium_1, and subpallium_2).** The top 30 enriched tissues are displayed.

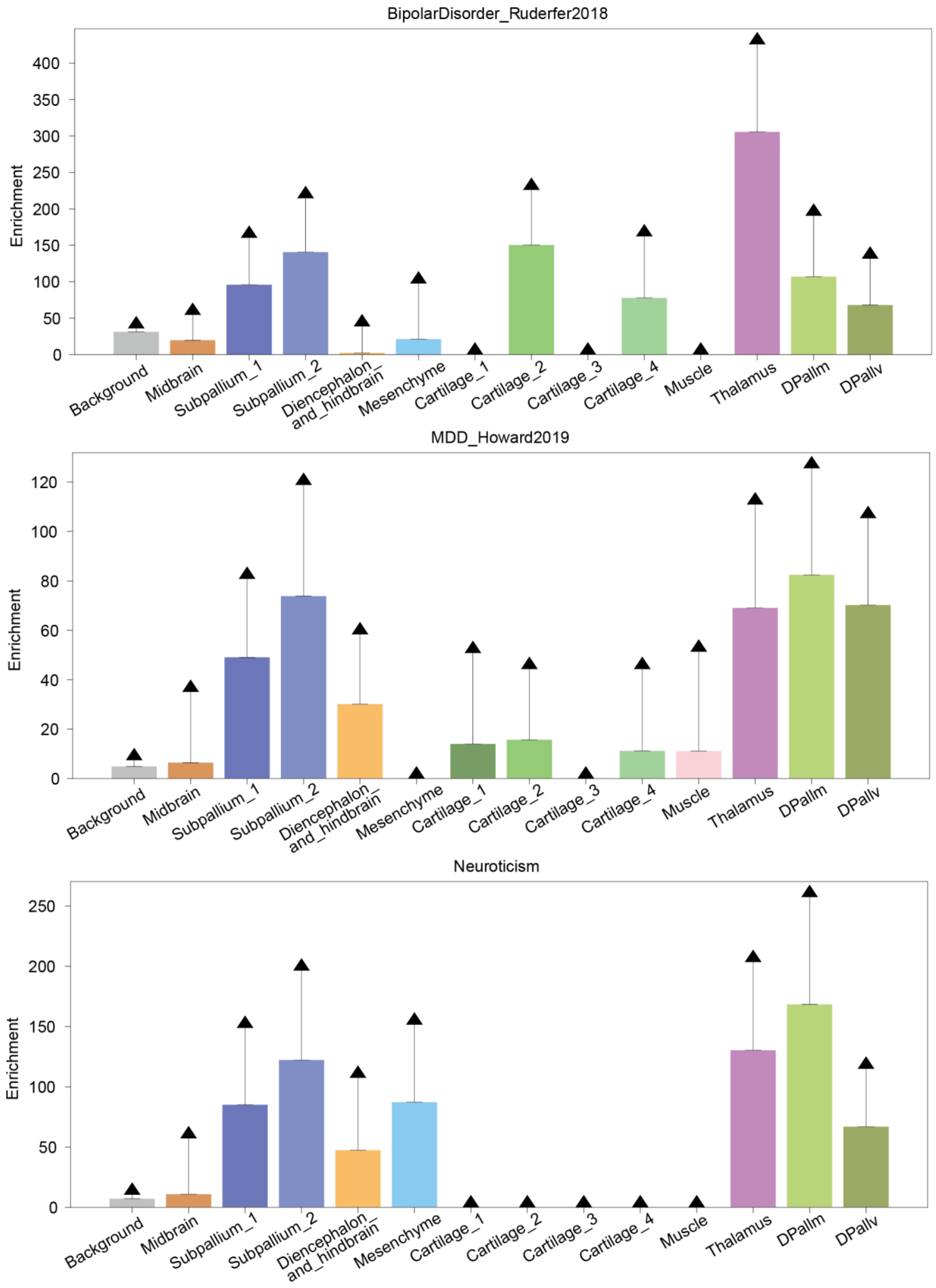

**Supplementary Fig. 29 | Partitioned heritability analysis of three mental diseases.** The enrichments of heritability in certain cerebral regions are significantly higher than that in mesenchyme, cartilage, and muscle.

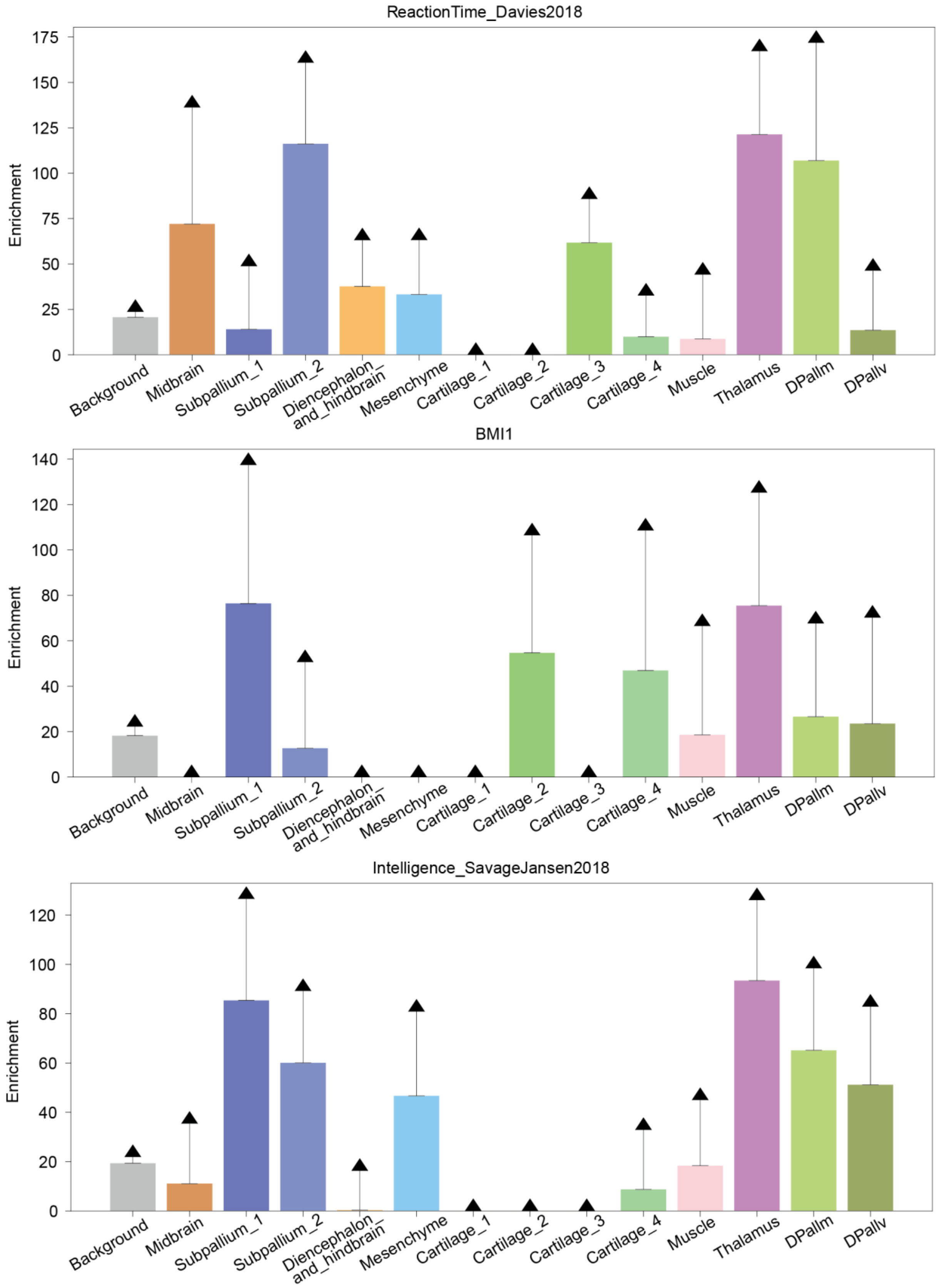

**Supplementary Fig. 30 | Partitioned heritability analysis of three mental-related phenotypes.** The enrichments of heritability in certain cerebral regions are significantly higher than that in mesenchyme, cartilage, and muscle.

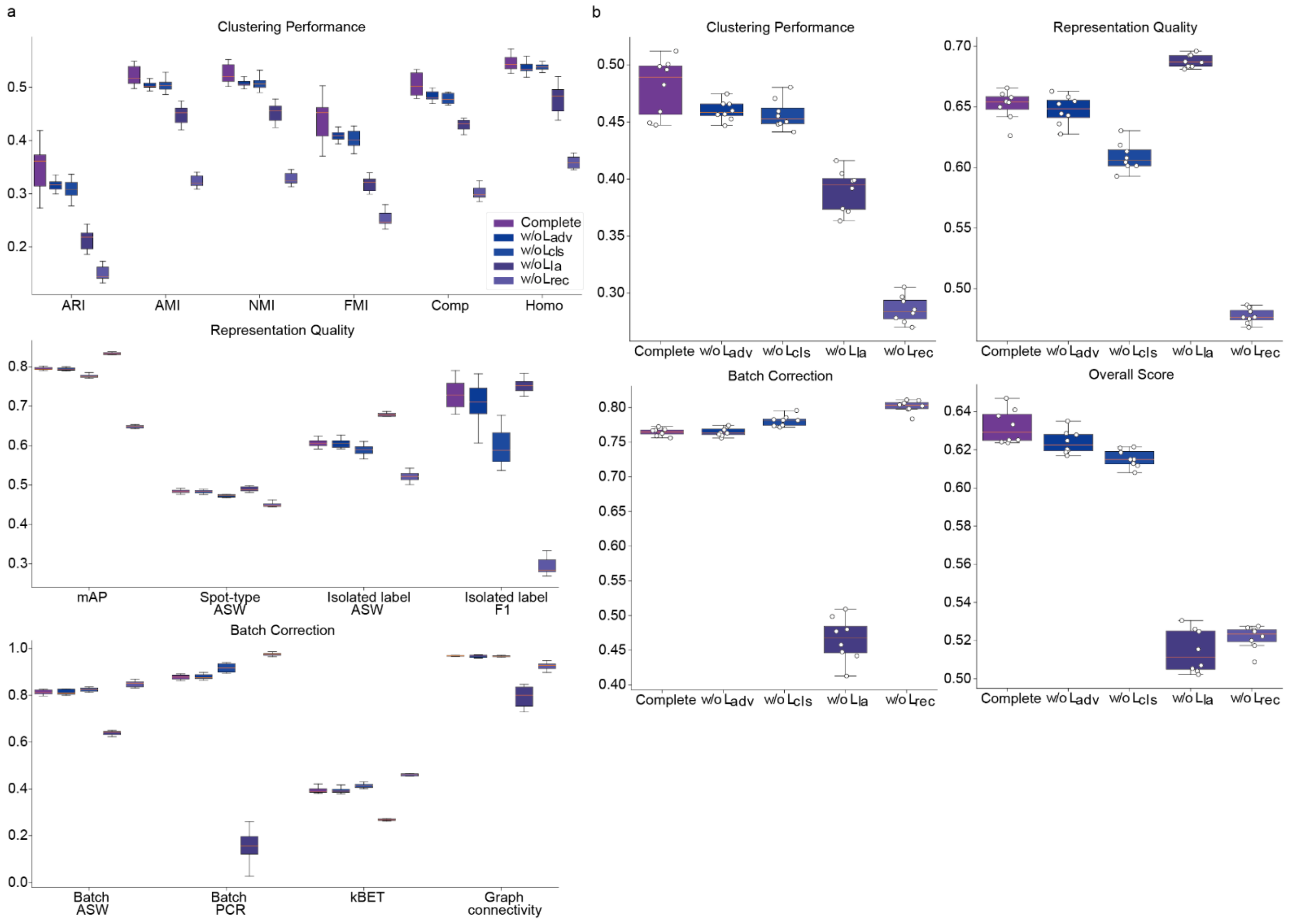

**Supplementary Fig. 31 | Ablation study on the loss functions of INSTINCT. a,** we compared the effects of removing each loss function from the model, respectively. Integration was performed on the MISAR-seq MB dataset, and 14 metrics were compared. **b,** the group-specific overall scores and the final overall score of different compositions of loss functions. The complete model of INSTINCT outperformed the model with any loss function removed in the final overall score.

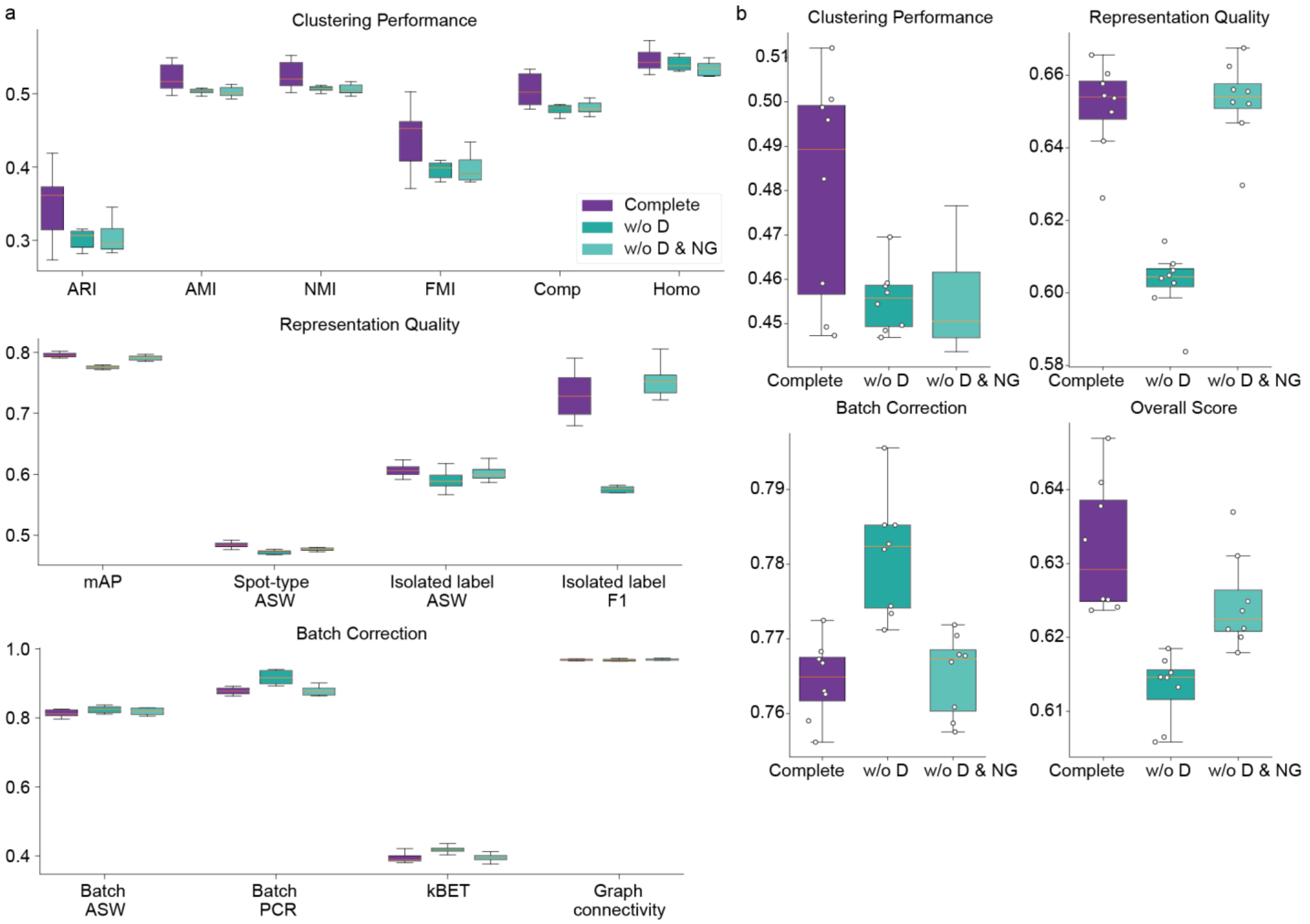

**Supplementary Fig. 32 | Ablation study on the model structures of INSTINCT. a,** we compared the effects of removing the discriminator (w/o D) and simultaneously removing both the discriminator and noise generator (w/o D & NG) from the model. Due to the structure of the INSTINCT model, it is not reasonable to solely remove the noise generator. Integration was performed on the MISAR-seq MB dataset, and 14 metrics were compared. **b,** the group-specific overall scores and the final overall score of different model structures. The complete model of INSTINCT outperformed the other two models in clustering performance, representation quality, as well as the final overall score.

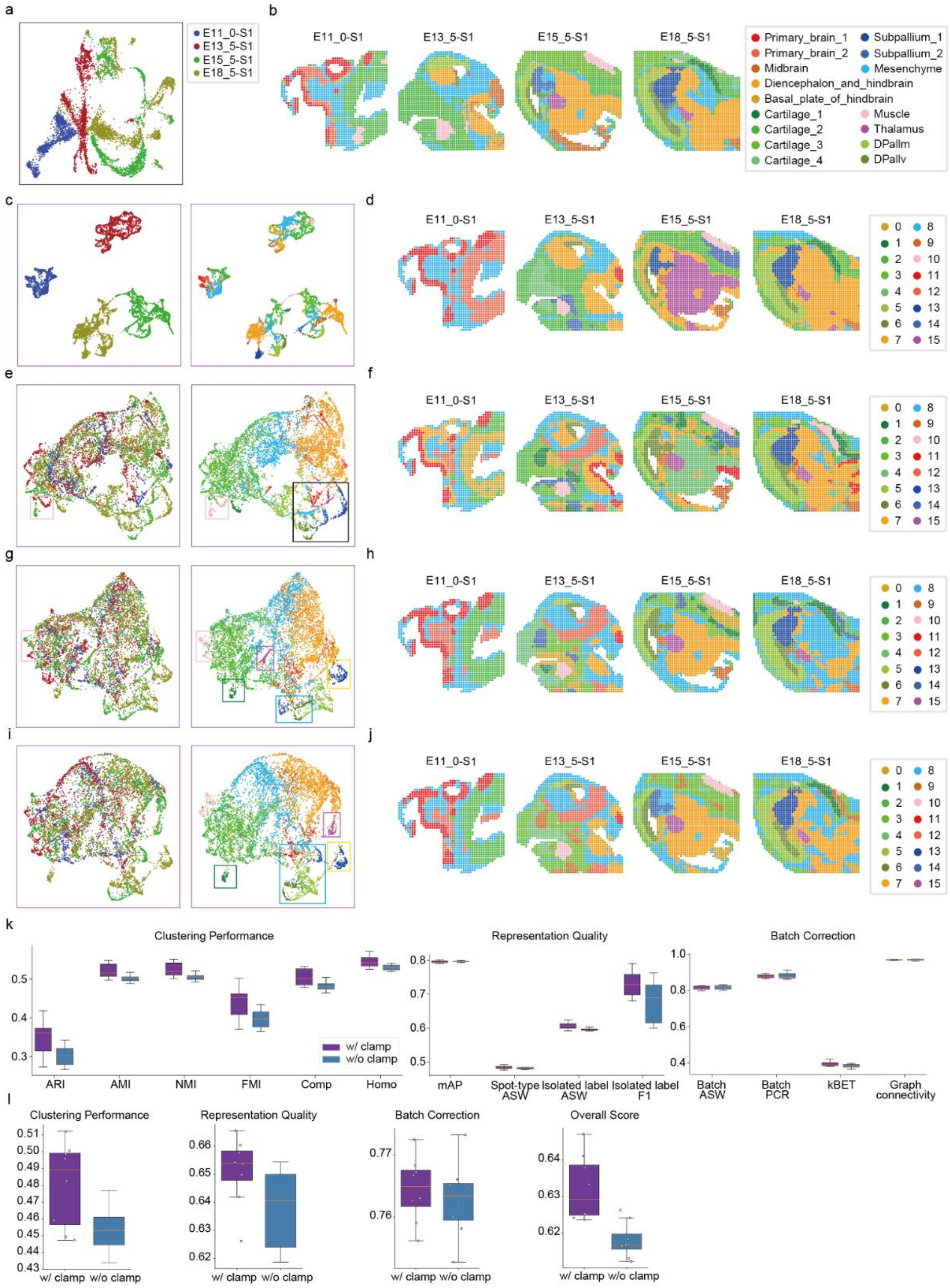

**Supplementary Fig. 33 | Step-by-step improvement of INSTINCT with gradual addition of mechanisms. a,** UMAP visualization of raw data from MISAR-seq MB dataset. **b,** four S1 slices of the MISAR-seq MB dataset. **c,** UMAP visualizations of the latent embeddings after training in stage 1. **d,** clustering results based on the latent embeddings after training in stage 1. **e,** UMAP visualizations of the latent embeddings obtained by adding the latent loss in stage 2. **f,** clustering results based on the latent embeddings obtained by adding the latent loss in stage 2. **g,** UMAP visualizations of the latent embeddings obtained by adding the latent loss and the noise generator in stage 2. **h,** clustering results based on the latent embeddings obtained by adding the latent loss and the noise generator in stage 2. **i,** UMAP visualizations of the latent embeddings obtained by adding the latent loss, the noise generator, and the stochastic domain translation procedure in stage 2. **j,** clustering results of the latent embeddings obtained by adding the latent loss, the noise generator, and the stochastic domain translation procedure in stage 2. **k,** comparison of the individual scores for the complete model and the model without the clamp strategy. **l,** comparison of the group-specific overall scores and the final overall score for the complete model and the model without the clamp strategy.

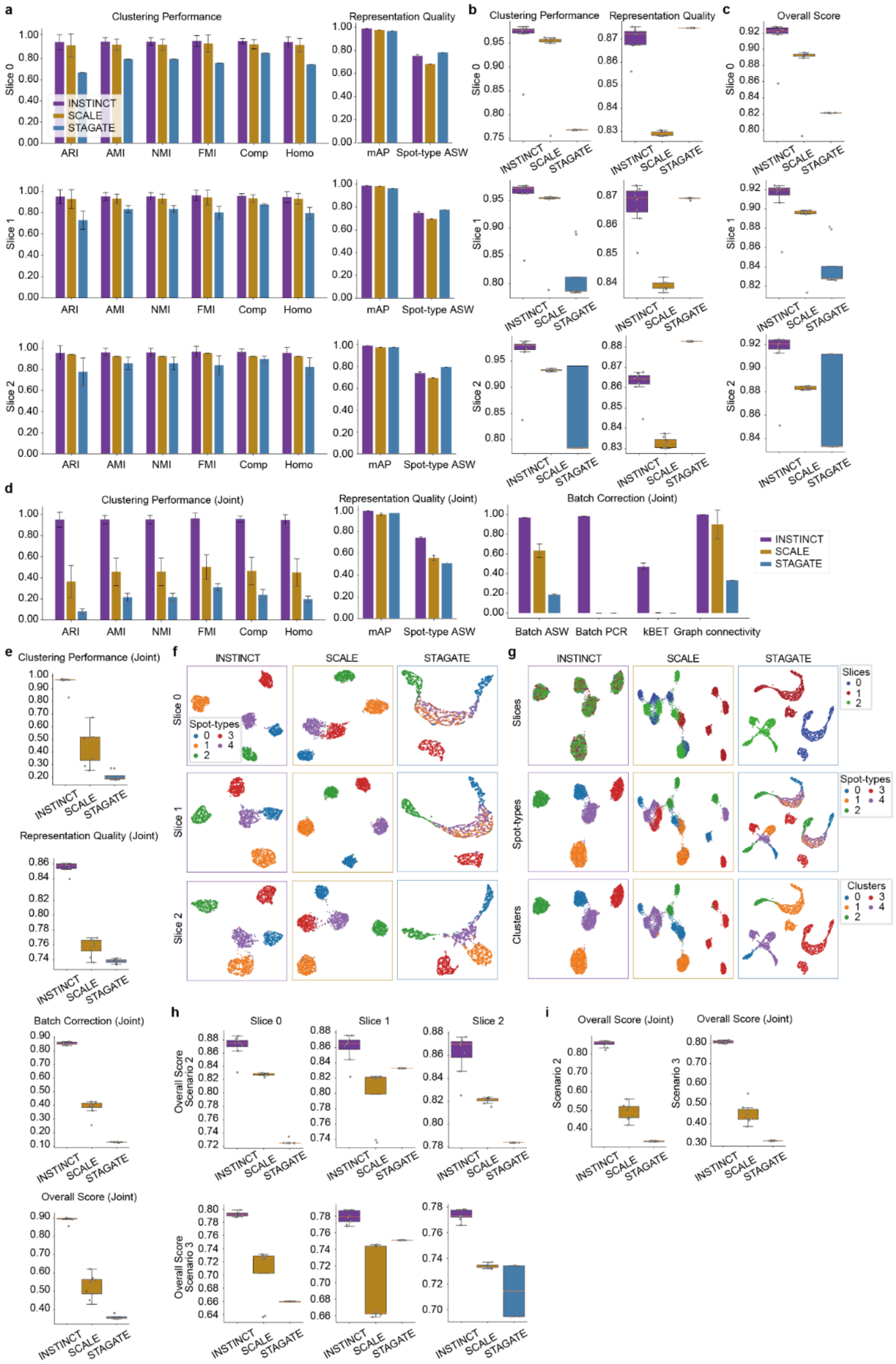

**Supplementary Fig. 34 | Comparison of INSTINCT with SCALE and STAGATE on simulated data. a,** individual scores of evaluation metrics for clustering performance and representation quality for each slice in simulated scenario 1. **b,** comparison of the group-specific overall scores for each slice. **c,** comparison of final overall scores for each slice. **d,** individual scores of metrics for clustering performance, representation quality, and batch correction for the joint evaluation of all slices. **e,** group-specific overall scores and final overall scores for the joint evaluation of all slices. **f,** UMAP visualizations of latent representations for each slice. **g,** UMAP visualizations of joint latent embeddings of the three slices, with spots colored based on slice affiliation (top), ground truth spot-type (middle), and mapped clustering result (bottom). **h,** comparison of final overall scores for each slice in simulated scenarios 2 and 3. **i,** final overall scores for the joint evaluation of all slices in simulated scenario 2 and 3, respectively.

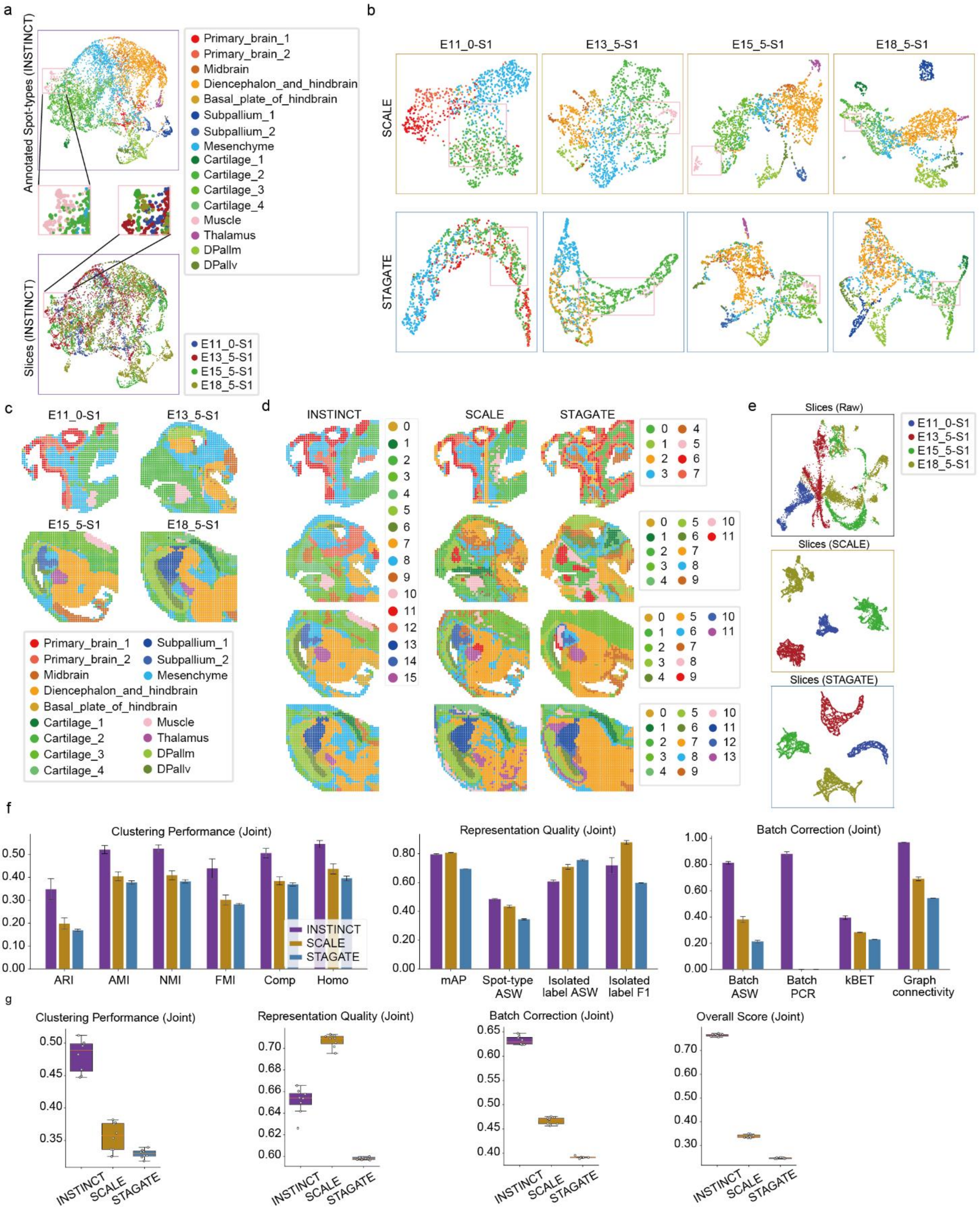

**Supplementary Fig. 35 | Comparison of INSTINCT with SCALE and STAGATE on the MISAR-seq MB dataset. a,** UMAP visualizations of the integrated results by INSTINCT, with spots colored based on slice affiliation and provided annotations. The muscle spot-type is highlighted. **b,** UMAP visualizations of latent embeddings for each slice obtained by SCALE and STAGATE, with muscle spot-type highlighted. **c,** four S1 slices of the MISAR-seq MB dataset, with spots colored based on provided annotations. **d,** clustering results from INSTINCT, SCALE, and STAGATE. Joint clustering of spots across all slices is performed for INSTINCT, while clustering was performed for each single slice based on the latent embeddings obtained by SCALE and STAGATE. **e,** UMAP visualizations of raw data and embeddings obtained by SCALE and STAGATE across all slices, with spots colored based on slice affiliation. **f,** individual scores of metrics for clustering performance, representation quality, and batch correction for the joint evaluation of all slices. **g,** group-specific overall scores and final overall scores for the joint evaluation of all slices.

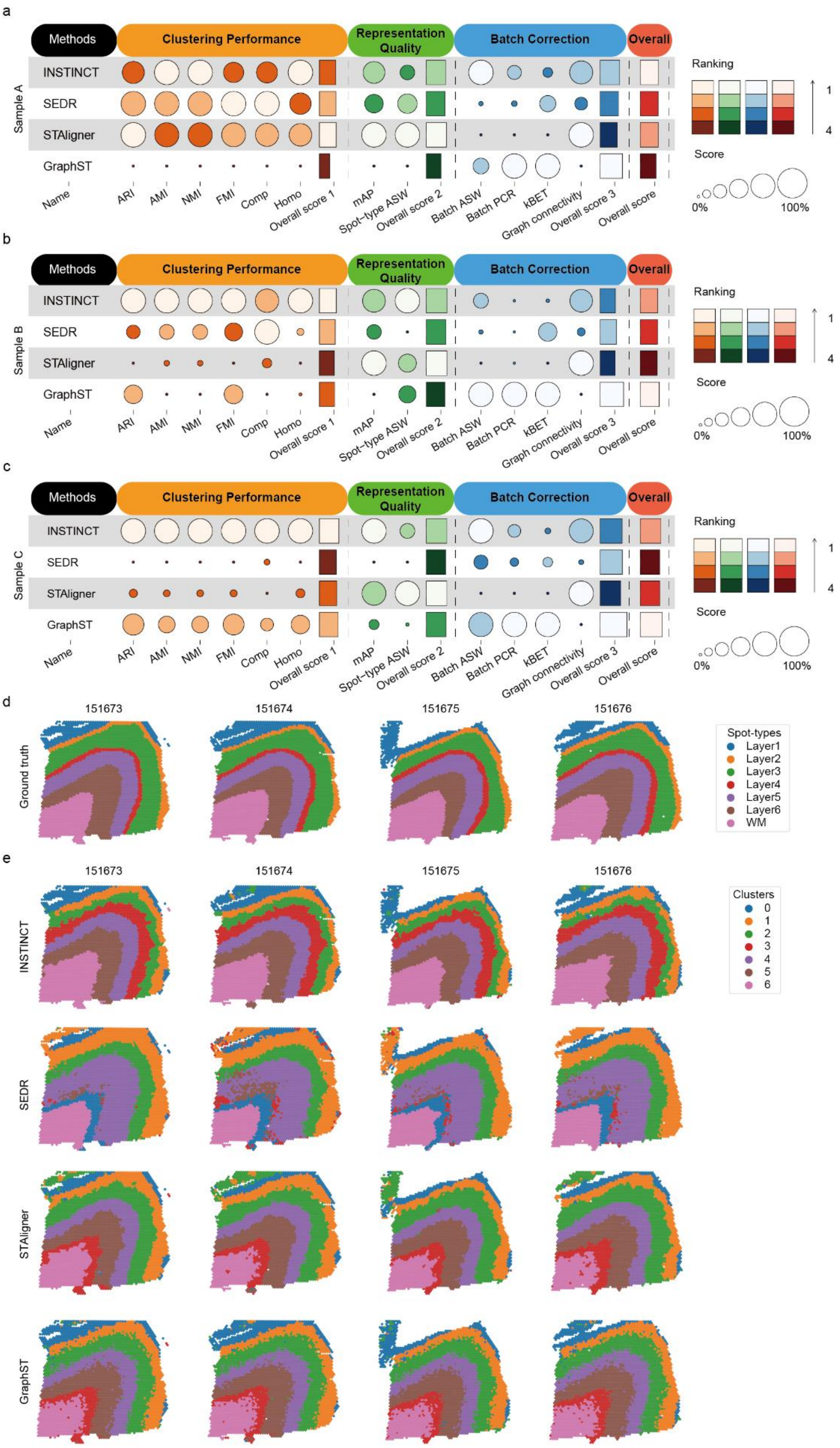

**Supplementary Fig. 36 | Integration results of INSTINCT and baseline methods on the three sample from the DLPFC dataset.** **a,** summary table of integration performance for INSTINCT and baseline methods on sample A, including scores of individual metrics for clustering performance, representation quality, and batch correction, as well as group-specific overall scores and final overall scores. **b,** summary table of integration performance for INSTINCT and baseline methods on sample B, **c,** summary table of integration performance for INSTINCT and baseline methods on sample C. **d,** provided annotations of different slices from sample C. **e,** clustering results of with identified clusters first mapped to true labels and then colored accordingly.

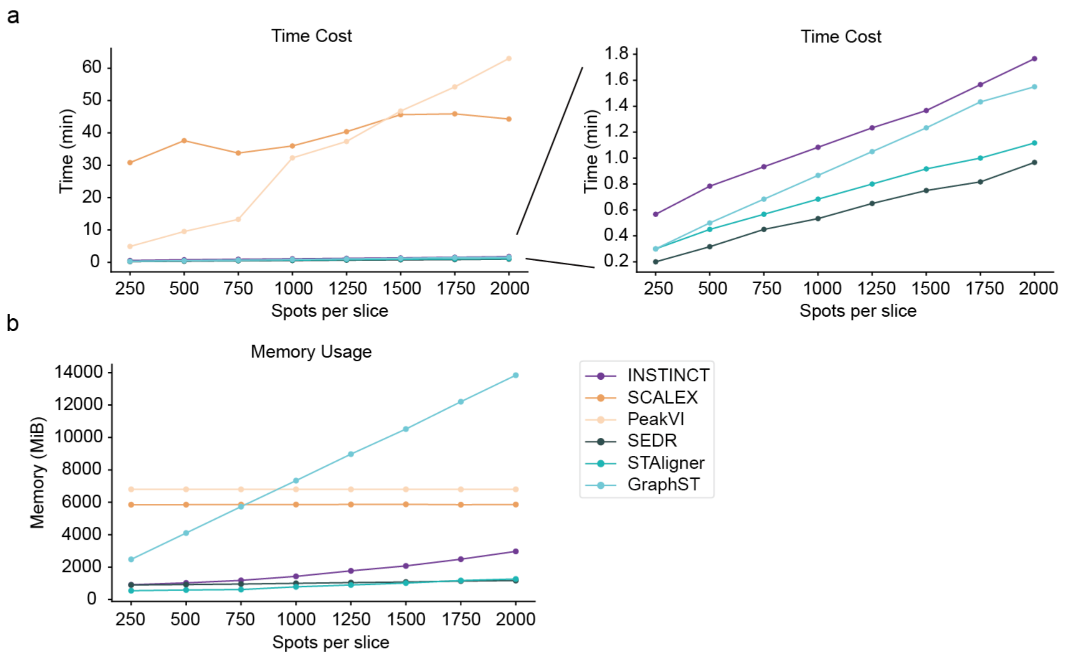

**Supplementary Fig. 37 | Comparison of time cost and memory usage between INSTINCT and baseline methods. a,** comparison of time cost between INSTINCT and baseline methods. **b,** comparison of memory usage between INSTINCT and baseline methods.

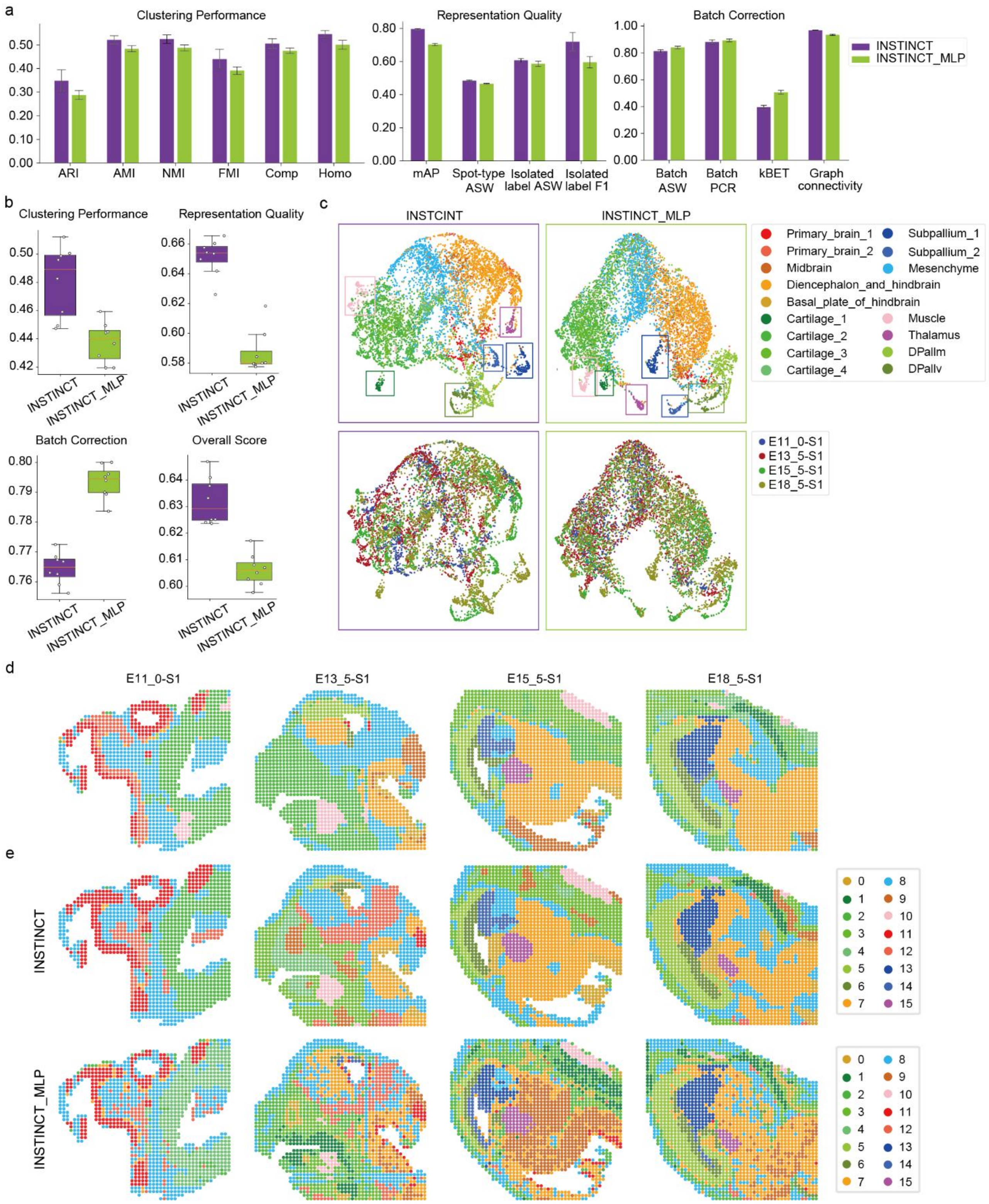

**Supplementary Fig. 38 | Comparison of the integration results between INSTINCT and INSTINCT_MLP. a,** individual scores of metrics for clustering performance, representation quality, and batch correction. **b,** group-specific overall scores and final overall scores. **c,** UMAP visualizations of integration results for INSTINCT and INSTINCT_MLP, with spots colored based on spot-type and slice affiliation. **d,** four S1 slices of the MISAR-seq MB dataset, with spots colored based on provided annotations. **e,** clustering results of INSTINCT and INSTINCT_MLP, with identified clusters first mapped to true labels and then colored accordingly.

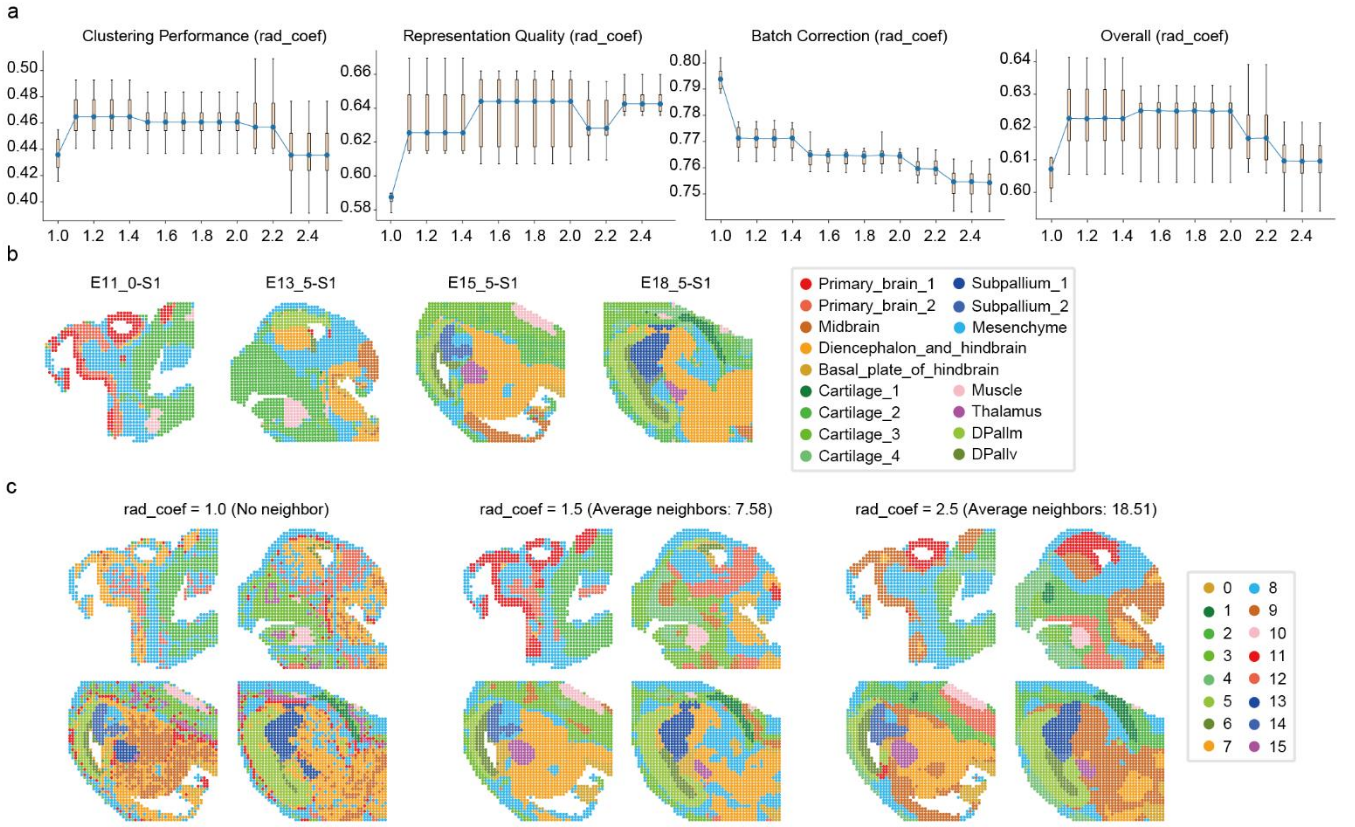

**Supplementary Fig. 39 | Sensitivity analysis for *rad_coef* (threshold for neighbor graph construction). a,** comparison of group-specific overall scores and final overall score for different values of *rad_coef*. **b,** four S1 slices of the MISAR-seq MB dataset, with spots colored based on provided annotations. **c,** clustering results when *rad_coef* is set to 1.0, 1.5, and 2.5 and the corresponding average amount of neighbors is 0, 7.58, and 18.51, respectively.

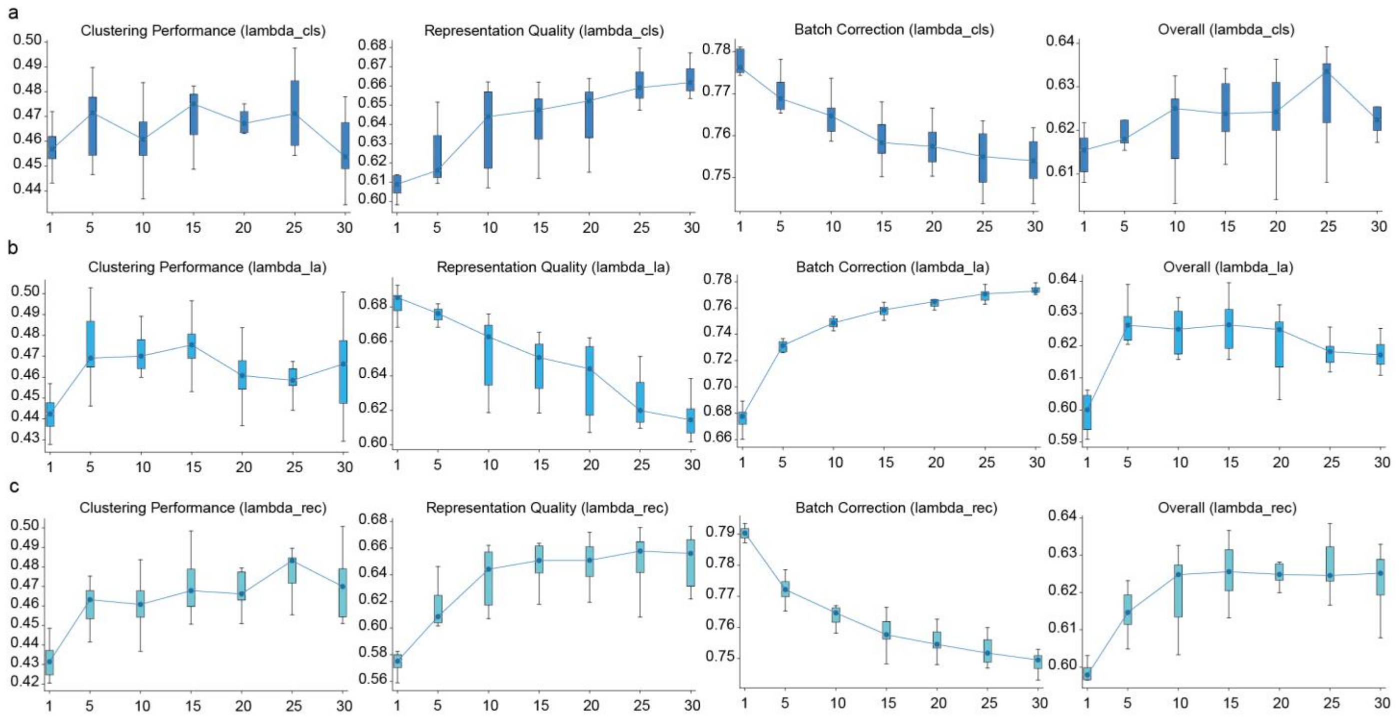

**Supplementary Fig. 40 | Sensitivity analysis for $\boldsymbol{\lambda_{cls}}$, $\boldsymbol{\lambda_{la}}$ and $\boldsymbol{\lambda_{rec}}$. a,** comparison of group-specific overall scores and final overall score for different values of $\boldsymbol{\lambda_{cls}}$. **b,** comparison of group-specific overall scores and final overall score for different values of $\lambda_{la}$. **c,** comparison of group-specific overall scores and final overall score for different values of $\boldsymbol{\lambda_{rec}}$.

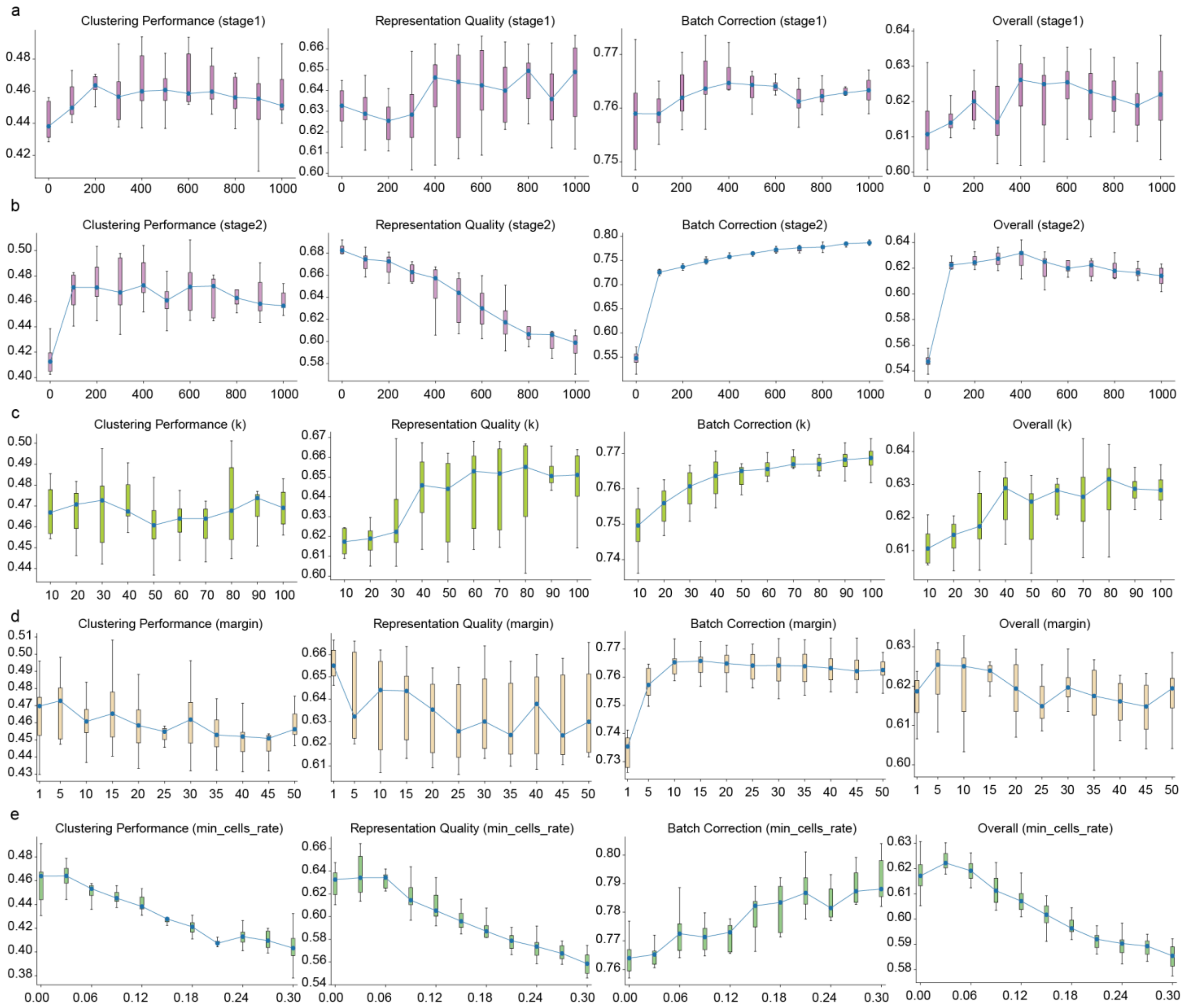

**Supplementary Fig. 41 | Sensitivity analysis for the number of training epochs, the selected neighbor count $k$, the margin for the clamp strategy, and the hyperparameter for peak filtering. a,** comparison of group-specific overall scores and final overall score for different values of the number of training epochs in stage 1. **b,** comparison of group-specific overall scores and final overall score for different values of the number of training epochs in stage 2. **c,** comparison of group-specific overall scores and final overall score for different values of the selected neighbor count $k$. **d,** comparison of group-specific overall scores and final overall score for different values of the margin for the clamp strategy. **e,** comparison of group-specific overall scores and final overall score for different values of the hyperparameter for peak filtering.

### Supplementary Texts

#### Supplementary Text 1: Comprehensive comparison on batch correction between INSTINCT and SCALEX

It is worth noting that in the three simulated scenarios (scenarios 1 to 3) used for quantitative comparison, as the complexity of the scenarios increases and the differentiation between different spot-types becomes more challenging, INSTINCT consistently achieves a higher group-specific overall score in batch correction than SCALEX (Supplementary Fig. 5a). Both INSTINCT and SCALEX show an increasing overall score in batch correction across the three scenarios. However, the increase in the batch correction score of SCALEX is more pronounced. Therefore, we will individually compare the four metrics used to evaluate batch correction through theoretical analysis and experimental evidence, and discuss the strengths and weaknesses of INSTINCT and SCALEX in handling batch effects.

Firstly, silhouette width is a measure of how similar a sample is to others of its own class compared to different classes, with ASW representing the average silhouette width for all samples. Batch ASW is the ASW score calculated with respect to slice labels, ranging from -1 to 1, where a score closer to 0 indicates that spots from different slices are well mixed. We computed it using the function implemented in the scib package, which scales the score to a value between 0 and 1, with a higher value indicating better mixing of data from different slices. In this metric, the score of INSTINCT gradually increases across the three scenarios, similar to SCALEX (Supplementary Fig. 5b). Although increase of SCALEX is slightly faster than that of INSTINCT, this is primarily because INSTINCT consistently mixes data from different slices very well in all three scenarios, achieving near-perfect scores in each. In the third scenario, the batch ASW score of INSTINCT is still significantly higher than that of

SCALEX. Therefore, we believe that INSTINCT outperforms SCALEX in this metric.

Principal component regression across batches (batch PCR) evaluates the effectiveness of batch correction by comparing the variance contribution of batch-specific information to the data matrix before and after integration. The concatenation of the raw data from multiple samples forms the data matrix before integration, and the concatenation of latent embeddings of spots represents the data matrix after integration. We used the function implemented in the scib package, which scales the batch PCR score to a value between 0 and 1, where a higher value indicates better elimination of batch-specific information. In this metric, SCALEX consistently achieves a perfect score of 1 across all three scenarios. While INSTINCT also achieves high score around 0.98 in each scenario, it does not match the performance of SCALEX (Supplementary Fig. 5c). We conducted a thorough analysis from the methodological perspective. SCALEX adopts a mini-batch strategy to iteratively optimize the model. In each mini-batch, it randomly samples data from all slices instead of from the same slice. SCALEX treats the input expression profile across all slices as a whole mixture distribution without distinguishing their sources. Specifically, after mapping the high-dimensional data to a lower-dimensional latent space using a batch-free encoder, it extracts the mean ($\mu$) and variance ($\sigma^2$) of the latent representations ($z$). A standard multivariate Gaussian prior is used for $z$, while the approximated distribution of $z$ is re-parameterized by $z = \mu + \sigma \times \varepsilon$, where $\varepsilon$ is sampled from $\mathbb{N}(0, I)$. One of the objectives of SCALEX is to minimize the Kullback-Leibler divergence between the posterior distribution $\mathbb{N}(\mu, \sigma^2)$ and the prior distribution $\mathbb{N}(0, I)$ of the latent representations. This mini-batch selection strategy, combined with a statistically-based modeling approach, effectively eliminates the impact of batch information on latent representations. In contrast, INSTINCT uses the stochastic domain translation process to remove batch effects, without enforcing a unified prior distribution of data from all slices. Therefore, while INSTINCT also effectively eliminates slice-specific information and achieves very high scores, it cannot guarantee a perfect score of 1. However, it has been demonstrated in the main manuscript that the mechanisms of INSTINCT help to retain more biological variations than SCALEX when applied to real datasets.

k-nearest-neighbor batch effect test (kBET) measures the removal of batch effects by using a Pearson's $\chi^2$-based test to determine whether the batch label distribution in the k nearest neighbors (kNN) of spots is similar to the global distribution of the batch labels. It is calculated within each spot type and returns the rejection rate, with a smaller value indicating better mixing of spots. We computed it using the function implemented in the scib package, which summarizes the kBET scores for all spot types and scales the overall score to a value between 0 and 1, where a higher value indicates better removal of batch effects. In this metric, INSTINCT consistently achieves significantly higher scores than SCALEX across the three scenarios (Supplementary Fig. 5d). Through visualization, we can observe that INSTINCT consistently mixes the data from the three slices evenly across all three scenarios and clearly distinguishes between the five different spot-types (Supplementary Fig. 5e). In scenario 1, while SCALEX can separate different spot-types, it does not mix the spots from slice 1 well with those from the other two slices, resulting in a noticeably low kBET score. As the scenarios become more complex and differentiation between different spot-types becomes more challenging, the mixing of data from the three slices improves for SCALEX, leading to an increase in kBET scores. However, in scenario 3, although SCALEX mixes the data from the three slices, it does not differentiate between the different spot-types as distinctly as INSTINCT does. This indicates that while SCALEX is effective at removing batch effects, it does not retain enough biological variations. In contrast, INSTINCT is able to better eliminate batch effects while preserving sufficient biological diversity.

Graph connectivity measures whether the kNN graph directly connects all spots of the same type. The computing method is detailed as follows:

$$\text{Graph connectivity} = \frac{1}{n}\sum_{i}\frac{\mathrm{LCC}(G_i)}{N_i}$$

where $n$ is the total number of spot-types, $N_i$ is the number of spots of the $i$-th spot-type, $G_i$ represents the subset kNN graph that contains only spots from the $i$-th spot-type, and $\mathrm{LCC}(\cdot)$ returns the number of nodes in the largest connected component of the graph. The value of graph connectivity score is higher than 0 and no larger than 1, with a higher value indicating that more spots with the same identity are connected in the integrated kNN graph. We computed it with the function implemented in the scib package. For this metric, INSTINCT and SCALEX perform similarly in scenario 1. However, as the scenarios become more complex, the score for INSTINCT decreases more significantly than for SCALEX (Supplementary Fig. 5f). This indicates that in this metric, INSTINCT does not perform as well as SCALEX. This difference is due to the method used to generate the simulated data. As detailed in the original manuscript, we first defined a single-cell resolution slice containing five spatial domains as a template. scATAC-seq data was then randomly assigned to each location based on a certain probability, and each $3 \times 3$ cell grid aggregated as a spot. The type of spot was determined by the majority cell-type within the grid. For each spatial domain, the cell-type corresponding to it was assigned with a probability of $r$, while the remaining cell-types were each assigned with a probability of $\frac{1-r}{4}$. Scenarios 1, 2, and 3 corresponded to $r$ values of 0.8, 0.7, and 0.6, respectively. This method causes some spots not belonging to the pre-defined domain to be randomly scattered within that domain. As the $r$ value decreases, the proportion of these scattered spots increases (Supplementary Fig. 5g). Since INSTINCT uses a GAT as the encoder, the low-dimensional representation of each spot incorporates information of its neighborhood, which has a smoothing effect and tends to identify more continuous spatial domains (Supplementary Fig. 5h). On the other hand, SCALEX, being a method developed for single-cell data, does not leverage spatial information and therefore has the ability to identify spots with different characteristics scattered within large domains. As a result, with an increasing number of scattered spots, the decline of graph connectivity score for INSTINCT is more pronounced compared to SCALEX.

To validate this conclusion, we removed the latent representations of spots that do not belong to the pre-defined spatial domains and recalculated the graph connectivity scores for INSTINCT and SCALEX, denoted as INSTINCT_del and SCALEX_del, respectively. We observed that INSTINCT achieved near-perfect scores in each scenario, while the scores for SCALEX remained largely unchanged compared to previous results, confirming our conclusion (Supplementary Fig. 5i). This indicates that SCALEX is better at identifying spots with different biological characteristics within large spatial domains, while INSTINCT excels at identifying more continuous spatial domains.

Based on the above analysis, it can be seen that INSTINCT and SCALEX each have their strengths and weaknesses in batch correction.

#### Supplementary Text 2: Comparison of INSTINCT with Harmony and Seurat

We compared INSTINCT with two classic integration methods, Harmony and Seurat. The comparison was first conducted on data from simulated scenario 1, which consists of three spatially resolved slices, each containing 2,000 spots across five spot types. For the PCA-based methods Harmony and Seurat, we set the input dimension the same as for INSTINCT, using the first 100 PCs of the data matrix for integration. We evaluated the latent embeddings of spots obtained by these three methods from three aspects: clustering performance, representation quality, and batch correction, using 12 metrics for quantitative comparison. INSTINCT ranked first in 11 individual metrics (Supplementary Fig. 7a) and scored significantly higher than Harmony and Seurat in the group-specific overall scores for all three aspects, as well as in the final overall score (Supplementary Fig. 7b).

Visualizations showed that, before integration, there were significant batch effects across the three slices in simulated scenario 1 (Supplementary Fig. 7c). After integration, INSTINCT was able to uniformly mix the data from the three slices while clearly distinguishing the five different spot-types (Supplementary Fig. 7d). Although Harmony also mixed the data from the three slices, many spots were scattered outside the main clusters, demonstrating low robustness. Seurat, on the other hand, failed to evenly mix spot-type 3 from slice 1 with that of the other two slices. Both Harmony and Seurat failed to distinguish between spot-type 1 and spot-type 4. Furthermore, for Harmony and Seurat, the cluster for spot-type 0 contained many misclassified spots from other types. In contrast, the cluster for spot-type 0 identified by INSTINCT contained nearly no other spot-types. This is because INSTINCT takes spatial location information into account, with each spot borrowing information from its spatial neighbors. These results indicate that Harmony and Seurat cannot preserve enough biological variations while removing batch effects to accurately distinguish different spot types, whereas

INSTINCT can.

We conducted similar experiments on simulated scenarios 2 and 3, where batch effects were progressively more complex and spot-types increasingly difficult to distinguish. In all three scenarios, INSTINCT significantly outperformed Harmony and Seurat in group-specific overall scores for all three aspects and in the final overall score (Supplementary Fig. 7e). These results demonstrate that traditional PCA-based integration methods like Harmony and Seurat perform poorly on simulated data and are clearly inferior to INSTINCT.

#### Supplementary Text 3: Comparison of INSTINCT with baseline methods on simulated scenario 7

To further explore the ability of INSTINCT to distinguish rare spot-types, we designed simulated scenario 7, which comprises more spot-types compared to the previous six scenarios. Specifically, we employed the discrete mode of simCAS, with the *tree_text* parameter set to "((((0:0.2,1:0.2):0.2,2:0.4):0.5,(3:0.5,4:0.5):0.4):0.1, ((5:0.1,6:0.4):0.3, 7:0.2):0.2);", producing three scATAC-seq datasets with varying batch effects. Each dataset comprises 8,000 cells distributed among eight cell-types, with 1,000 cells per type labeled as 0 to 7. Then, following the approach used for generating the spATAC-seq data in the previous six scenarios, we employed another pre-defined single-cell resolution slice as the template and generated three spatial resolution slices, each comprising eight spot-types (Supplementary Fig. 9a). Each slice contains a total of 2,000 spots, with spot-types 5, 6, and 7 being rare, each comprises approximately 50 spots per slice. The slices exhibit significant batch effects (Supplementary Fig. 9b).

We then applied INSTINCT and baseline methods to integrate these three slices and quantitatively compared the integration results from three aspects: clustering performance, representation quality, and batch correction, using a total of 12 metrics. INSTINCT achieved the highest score in eight metrics, and second place in two. It also ranked first in group-specific overall scores of clustering performance and representation quality, as well as the final overall score (Supplementary Fig. 9c).

Through visualization, we found that INSTINCT not only effectively eliminated batch effects across the three slices but also perfectly distinguished the three rare spot-types (Supplementary Fig. 9d). GraphST managed to distinguish the three rare spot-types in slices 0 and 2 with less precision but failed to eliminate batch effects between slice 1 and the other two slices. STAligner mixed spots from all three slices but could only distinguish one rare spot-type (spot-type 7). PeakVI could identify spot-types 6 and 7 in slice 1 but did not correct batch effects between the three slices. Other baseline methods failed to identify any rare spot-types.

The quantitative analysis and visualization of simulated scenario 7 further demonstrate the superior ability of INSTINCT to simultaneously remove batch effects and distinguish rare spot-types.

#### Supplementary Text 4: Comparison of INSTINCT with baseline methods on the low-quality spatial ATAC ME dataset

We applied INSTINCT to a mouse embryo dataset generated by spatial ATAC (spatial ATAC ME), which consists of six annotated slices from developmental stages E12.5, E13.5, and E15.5, with two vertically adjacent slices from each stage (Supplementary Fig. 14a, b). However, we found that, compared to MISAR-seq MB dataset, which is also annotated, each slice in the spatial ATAC ME dataset has significantly lower fragment counts captured within spots (Supplementary Fig. 14c), and the ratio of reads in transcription start sites (TTS) was also notably lower (Supplementary Fig. 14d), indicating extremely low sequencing depth and data quality for the spatial ATAC ME dataset. Despite this, our method still outperformed other baseline methods on this low-quality dataset.

Initially, we utilized INSTINCT alongside the six baseline methods to integrate the S1 slices of the three stages and calculated evaluation metrics based on their provided annotations. INSTINCT ranked first on 8 of the 12 metrics (Supplementary Fig. 15a). It also secured the first place in the group-specific scores for clustering performance and representation quality, as well as the final overall score (Supplementary Fig. 15b).

However, its performance in batch correction was not particularly high. We observed that most baseline methods failed to accurately capture biological variations. Visualization results showed that these methods improperly mixed data from the three slices, leading to a blend of various spot-types that made it challenging to discern any specific spot-type accurately (Supplementary Fig. 16a, b). Consequently, these methods had low representation quality and clustering scores but relatively higher batch correction performance. In contrast, INSTINCT successfully distinguished the liver from other spot-types and avoided excessive mixing of all slices (Supplementary Fig. 16a, b). It accurately categorized these spots into four major groups: central nervous system (CNS), meningeal and peripheral nervous system (PNS), mesenchyme and limb, and liver, consistent with the results of the original study.

Through clustering, while other methods struggled to identify any spatial domain accurately, INSTINCT performed significantly better (Supplementary Fig. 16c, d). For instance, it accurately identified the liver (cluster 5) in E12.5 and E13.5 slices. In the E15.5 slice, it delineated spatial relationships within brain tissue, distinguishing between the forebrain (cluster 1), midbrain (cluster 8), and hindbrain (cluster 2). Although Scanorama was able to identify the periventricular category that INSTINCT did not, it failed to accurately identify any other domain across the three slices.

Additionally, we utilized INSTINCT to integrate S2 slices from the three developmental stages, as well as collectively integrating all six slices, and compared with other methods, obtaining results consistent with those of integrating S1 slices (Supplementary Fig. 17 to Supplementary **Fig. 23**).

These results showcase that INSTINCT is relatively more accurate in capturing the underlying biological variations compared to other methods when integrating samples of low quality.

#### Supplementary Text 5: Analysis of the mechanisms implemented in INSTINCT

We validated the effectiveness of the mechanisms implemented in INSTINCT using the MISAR-seq MB dataset (Supplementary Fig. 33a, b). Initially, we trained the model only in stage 1. After training, the batch effects between different slices remained very prominent (Supplementary Fig. 33c), which indicates that stage 1 merely serves as a pretraining step and does not have the capability of batch correction. This made it challenging to accurately identify associations between different slices during clustering, such as mistakenly classifying mesenchyme from E15.5 and E18.5 as diencephalon and hindbrain, or misclassifying diencephalon and hindbrain from the E15.5 slice as thalamus (Supplementary Fig. 33d), hindering the integrated analysis across slices.

Subsequently, we enabled the stage 2 for training. Compared to the stage 1, we initially added only the latent loss to train the model. As observed, since we directly sought neighbors in the low-dimensional representation matrix $Z$ without the noise generator to separate batch effects from the low-dimensional representations of spots, although latent loss caused the four slices to mix, the identification of positive samples was not precise enough. This led to many spot-types clustering together, making distinctions less clear, as highlighted by the black box in Supplementary Fig. 33e. Consequently, during spatial domain identification, some types were misclassified, such as incorrectly identifying cartilage_2 in the E11.0 slice as basal plate of hindbrain (Supplementary Fig. 33f). This indicates that using only the latent loss for batch correction does not guarantee the effective preservation of biological variations.

We then further incorporated the noise generator into the stage 2 model to capture patterns of noise and batch effects. After integration, compared to the model with only the latent loss,

INSTINCT distinguished previously clustered rare spot-types more clearly (Supplementary Fig. 33g). For instance, in the model with only latent loss, the muscle category included spots from only the last three slices, whereas after adding the noise generator, the muscle category included spots from all four slices, as highlighted by the pink boxes in Supplementary Fig. 33e and Supplementary Fig. 33g. Additionally, compared to the model with only the latent loss, the model with the noise generator accurately distinguished the cartilage_2 in E11.0 and the diencephalon and hindbrain of E15.5 (Supplementary Fig. 33h), which the previous model failed to do. This indicates that the noise generator allows the encoder to better retain biological information.

Finally, we applied the complete model with the stochastic domain translation process to the data. Compared to the previous version of the model without this process, the differences between spot-types were more pronounced. For example, the complete model distinguished DPallv from subpallium_2, cartilage_1 from other cartilage subtypes, and subpallium_1 from diencephalon and hindbrain more clearly than the previous version (Supplementary Fig. 33i). Additionally, it was able to differentiate thalamus, which the previous model could not, as highlighted in Supplementary Fig. 33g and Supplementary Fig. 33i. Consequently, the complete model achieved better spatial domain identification results, such as correctly identifying cartilage_1, which is present only in E18.5 (Supplementary Fig. 33j), while the previous model mistakenly classified some cartilage subtypes from E15.5 as cartilage_1. This indicates that stochastic domain translation allows the noise generator to more purely simulate noise and batch effects, further ensuring that the encoder extracts sufficient biological variations crucial for distinguishing different spot-types.

Moreover, we conducted an ablation study on the clamp strategy. Among all 14 metrics, the complete model outperformed the model without the clamp strategy in 11 metrics

(Supplementary Fig. 33k). We also found that adopting this strategy led to improvements in all three aspects (Supplementary Fig. 33l). Notably, employing this strategy enhanced the performance of INSTINCT on isolated label ASW and isolated label F1-score, demonstrating its ability to preserve sample-specific spot-types.

#### Supplementary Text 6: Comparison of INSTINCT with single-slice analysis methods

To clarify the significance of integrating multiple samples, we compared INSTINCT with two other methods: SCALE and STAGATE. SCALE is designed for single-cell assay for scATAC-seq data, while STAGATE is used for SRT data. Both methods are developed for single-sample analysis and do not include modules for batch correction.

The comparison was first conducted on data from simulated scenario 1. Initially, we integrated the three slices using INSTINCT, obtaining shared latent embeddings for all spots across the slices. For the comparison of each specific slice, we extracted the latent embeddings of the spots from the corresponding slice and compared them with the representation results from SCALE and STAGATE, which were applied individually to the data from that slice. Since the analysis was conducted on individual slices, we evaluated clustering performance and representation quality using a total of eight evaluation metrics for quantitative comparison. Across all three slices, INSTINCT achieved the highest score in seven metrics and the second-highest in one metric (Supplementary Fig. 34a). Besides, for slice 1, INSTINCT obtained the highest group-specific overall scores in both clustering performance and representation quality (Supplementary Fig. 34b). Although for slices 0 and 2, the overall scores of INSTICNT in representation quality were slightly lower than that of STAGATE, it significantly outperformed STAGATE in the overall scores for clustering performance. This led to INSTINCT achieving the best final overall score for each slice (Supplementary Fig. 34c).

Furthermore, we jointly evaluated the latent embeddings of spots across all three slices in scenario 1 from three aspects: clustering performance, representation quality, and batch correction, using 12 metrics for quantitative comparison. INSTINCT ranked first in all 12 individual metrics, in the group-specific overall scores for all three aspects, and in the final overall score (Supplementary Fig. 34d, e). For SCALE and STAGATE, significant drops were observed in the metrics for clustering performance and representation quality, with the exception of mean average precision (mAP).

Next, we visualized the latent representations of spots for each slice (Supplementary Fig. 34f). INSTINCT and SCALE perfectly separated the five unique spot-types, demonstrating their ability to capture biological variations within a single slice. STAGATE managed to separate the five spot-types in slice 2 but erroneously mixing spot-type 1 and 4 in slice 0 and 1. We then visualized the latent embeddings for spots across all three slices (Supplementary Fig. 34g). INSTINCT not only evenly mixed the spots from the three slices but also still ideally separated all five spot-types. The joint clustering based on the latent embeddings obtained by INSTINCT led to satisfied results consistent with the ground truth. However, noticeable batch effects still remain in the latent embeddings of the three slices obtained by SCALE and STAGATE, with no clear evidence that the same spot-types could be identified across different slices. This led to different spot-types being incorrectly grouped together during joint clustering, proving that the spot representations obtained from single-slice analyses are not comparable and cannot be analyzed jointly. In contrast, INSTINCT not only provides the most effective representation but also enables the simultaneous analysis of different slices, expanding the potential scenarios for analysis. For simulated scenarios 2 and 3, where batch effects are more pronounced and distinguishing between different spot-types is more challenging, INSTINCT continued to achieve the best performance both in individual slice analyses and in joint analysis (Supplementary Fig. 34h, i). This further validates the advantage of the integrative analysis of INSTINCT over single-slice analysis.

Subsequently, we compared the integration results of INSTINCT on the four S1 slices of the MISAR-seq MB dataset, which has provided annotations, with the results obtained by analyzing each slice separately using SCALE and STAGATE. We found that the integration of multiple samples by INSTINCT allowed for the differentiation of certain spot-types that were difficult to distinguish in single-slice analyses. As highlighted in Supplementary Fig. 35a, the UMAP visualizations of the integration results from INSTINCT show that the muscle spot-type from all four slices is clustered together and separated from other types of spots. However, when analyzing each slice individually, SCALE could only partially distinguish the muscle category from other categories in the E13.5 and E15.5 slices, but failed to do so in the E11.0 and E18.5 slices. STAGATE, on the other hand, was unable to distinguish the muscle category from other categories in any of the four slices (Supplementary Fig. 35b).

We performed joint clustering on the latent embeddings obtained from the integration of all four slices using INSTINCT. For SCALE and STAGATE, we clustered the embeddings of each slice separately, with the number of clusters set to match the number of annotated spot-types for each slice. We found that the clustering results from integrative analysis of INSTINCT led to better outcomes for spatial domain identification (Supplementary Fig. 35c, d). For example, INSTINCT accurately identified the muscle regions (cluster 10) in the E13.5, E15.5, and E18.5 slices, whereas SCALE only identified the muscle category in the E13.5 slice, and STAGATE failed to accurately identify the muscle category in any of the four slices. Additionally, INSTINCT correctly identified the thalamus (cluster 15), which is present only in the E15.5 and E18.5 stages. In the E18.5 slice, SCALE incorrectly identified part of the basal plate of the hindbrain as the thalamus, and STAGATE completely misidentified this region. Furthermore, the Subpallium_2 subclass is mainly present in the E15.5 stage and appears as a few spots below the Subpallium_1 subclass in the E18.5 stage. SCALE failed to identify this region in the E18.5 slice, and STAGATE overestimated its size, while INSTINCT accurately identified this region in the E18.5 slice (cluster 14).

Visualizations showed that the batch effects between the four slices were not eliminated in the representations obtained by SCALE and STAGATE (Supplementary Fig. 35e), further proving that the representations of individual slices cannot be used for joint analysis, thereby hindering comparisons across multiple slices (Supplementary Fig. 35f, g).

#### Supplementary Text 7: Comparison of INSTINCT with SRT data integration methods on human DLPFC dataset

We collected the human dorsolateral prefrontal cortex (DLPFC) dataset to validate the performance of INSTINCT on SRT data. This dataset is a classic dataset in the SRT field and is widely used in related researches. It comprises three samples (sample A, B, and C), each containing four vertically adjacent slices (slice A, B, C, and D).

For the SRT data integration task, we applied a different preprocessing procedure compared to that used for spATAC-seq data. We first concatenated the slices to be integrated, then identified 5,000 highly variable genes from the intersect gene set. We scaled the counts to 10,000 for each spot in the original data, followed by log-normalization. We then extracted the data corresponding to these 5,000 genes, and reduced the dimensionality of the data matrix to 100 using PCA as input for INSTINCT. We respectively integrated the three samples using INSTINCT and compared the results with methods developed specifically for SRT data, including SEDR, STAligner, and GraphST, which were used as baseline methods in the main manuscript.

INSTINCT achieved comparable metrics scores to these mature SRT methods, obtaining the highest final overall score for sample A and second place for samples B and C, slightly below GraphST (Supplementary Fig. 36a, b, c). Notably, for sample C, INSTINCT accurately identified the spatial arrangement of white matter (WM) and different layer structures (Supplementary Fig. 36d, e). Specifically, INSTINCT accurately identified the structure of WM, whereas other methods incorrectly divided WM into two layers. Additionally, SEDR and STAligner misidentified part of the outermost layer 0 as layer 2 and layer 3, respectively, while INSTINCT and GraphST accurately recognized layer 0. However, data integration and batch correction performed by GraphST rely on a pre-alignment process for the slices, and the effectiveness of its integration depends on the accuracy of this pre-alignment. In this experiment, we followed its tutorial and used PASTE as the pre-alignment algorithm. In contrast, INSTINCT does not require any pre-alignment procedure.

This case demonstrates that, despite being developed for spATAC-seq data integration, INSTINCT can achieve performance comparable to methods designed for SRT data integration, and even surpasses certain methods in some cases.

#### Supplementary Text 8: Comparison of time cost and memory usage for INSTINCT and baseline methods

We first test the time cost and memory usage of different methods on simulated scenario 1, which contains three slices. We randomly retained a certain number of spots from each slice, varying the number from 250 to 2000, generating a total of eight datasets. Next, we integrated these eight datasets using INSTINCT and baseline methods, recording the time cost and GPU memory usage. Scanorama was excluded because its GitHub version does not support GPU computation. We found that the time cost of INSTINCT was minimal, typically completing the integration task within several minutes, significantly lower than the single-cell data integration methods SCALEX and PeakVI, and comparable to integration methods developed for SRT data. Moreover, as the number of spots increased, the time cost of INSTINCT showed a linear growth (Supplementary Fig. 37a). However, in terms of memory usage, although INSTINCT was lower than GraphST (using PASTE as the prealign method), SCALEX, and PeakVI, it exhibited a polynomial growth pattern (Supplementary Fig. 37b).

In Supplementary Tab. 2 and Supplementary Tab. 3, we recorded the time cost and memory usage of each method on four real datasets (detailed information of each sample can be found in Supplementary Tab. 1). Although INSTINCT had relatively low time costs, its memory usage was relatively high. Overall, we suggest that the current memory usage and time cost are acceptable, allowing INSTINCT to be deployed and used on personal computers in most cases. However, we will work to reduce the memory usage of INSTINCT in future online version of codes.

#### Supplementary Text 9: Analysis of the use of the GAT encoder module

We investigated the differences in integration performance between using GAT and using MLP as encoders, and to demonstrate the importance of GAT in our model.

Firstly, we replaced the encoder in INSTINCT with an MLP of a similar structure while keeping all other components unchanged, resulting in a new model named INSTINCT_MLP, which does not include the GAT mechanism. This means that when mapping spots to the shared latent space, the information of their surroundings are not considered. We compared the integration results of INSTINCT and INSTINCT_MLP using a total of 14 metrics, focusing on clustering performance, representation quality, and batch correction. We found that INSTINCT outperformed INSTINCT_MLP in 11 of these metrics (Supplementary Fig. 38a). It also showed better performance in group-specific overall scores for both clustering performance and representation quality, as well as in the final overall score (Supplementary Fig. 38b).

Through visualizations, we observed that both INSTINCT and INSTINCT_MLP were able to identify six rare spot-types without a significant difference in performance (Supplementary Fig. 38c). This suggests that the stochastic domain translation process, a core component of our method, is crucial for achieving satisfying integration results.

However, INSTINCT demonstrated a clear advantage over INSTINCT_MLP in clustering results. The use of GAT allowed INSTINCT to identify more accurate and continuous spatial domains. For instance, the identification of muscle in E13.5 by INSTINCT was closer to the true structure compared to INSTINCT_MLP (Supplementary Fig. 38d, e). Additionally, INSTINCT accurately identified the cartilage_1 domain, which was present only in E18.5, whereas INSTINCT_MLP not only inaccurately identified this domain in E18.5 but also incorrectly recognized some other subtypes of cartilage in E13.5 and E15.5 as cartilage_1. This proves that the use of GAT enhances the integration performance of INSTINCT, particularly in the spatial domain identification task.

#### Supplementary Text 10: Sensitivity analysis for hyperparameters

For the sensitivity analyses, we first set a range of test values for each hyperparameter, including those for neighbor graph construction, loss function weights, training epochs, the number of selected nearest neighbors during stage 2, the margin for the clamp strategy, and peak filtering. Then, we conducted eight experiments on the MISAR-seq MB dataset with different random seeds for each value. The experiments were compared from three aspects, including clustering performance, representation quality, and batch correction.

When constructing the neighbor graph used as input, it is necessary to define whether two spots are considered neighbors based on their distance. To avoid the need for manually setting different rationale for different datasets, as required by STAligner, we introduced a hyperparameter *rad_coef*. This hyperparameter is multiplied by the minimum distance between spots within a single slice to determine the threshold distance for the corresponding slice. If the distance between two spots is less than this threshold, they are considered as neighbors. We varied the hyperparameter value from 1.0 to 2.5 across 16 experiments (Supplementary Tab. 5). We found that as the average number of neighbors for spots increased, the clustering performance initially improved and then declined. Representation quality was optimal in the mid-range (average neighbors: 7.58), while batch correction consistently decreased. The final overall score also first increased and then decreased, indicating that both too few and too many average neighbors are detrimental to integration (Supplementary Fig. 39a). According to clustering results when *rad_coef* was set to 1.0, 1.5, and 2.5, with corresponding average numbers of neighbors being 0, 7.58, and 18.51 respectively, we observed that with no neighbors, the GAT does not function, leading to discontinuous clustering results and failure to identify some spatial domains like the thalamus (Supplementary Fig. 39b, c). When the number of neighbors was 7.58 or 18.51, clustering results were generally continuous, but with 18.51 neighbors, the volumes of many spatial domains were overestimated, such as the subpallium_1 and cartilage_1 in E18.5 slice, and subpallium_2 in E15.5 slice. STAligner suggests keeping the average neighbor count between 4 and 15. We set this hyperparameter to 1.5, which typically keeps the average neighbor count within this range.

Then, we conducted a sensitivity analysis for the hyperparameters $\lambda_{cls}$, $\lambda_{la}$ and $\lambda_{rec}$, which control the contribution of the classification loss, latent loss, and reconstruction loss to the total objectives, respectively.

We found that as $\lambda_{cls}$ increased, clustering performance remained relatively stable, while representation quality and batch correction scores showed opposite trends: the former gradually increased, while the latter gradually decreased (Supplementary Fig. 40a). Based on the final overall score, we recommend setting the value of $\lambda_{cls}$ between 10 and 25. In all the experiments in this manuscript, this hyperparameter was fixed at 10.

For $\lambda_{la}$, we observed that as its value increased, changing in scores for representation quality and batch correction also exhibited opposite trends: representation quality decreased while batch correction improved (Supplementary Fig. 40b). Since the final overall score initially increased and then decreased, and considering that latent loss directly controls the degree of mixing between different slices, we recommend setting $\lambda_{la}$ to a moderate value, between 10 and 20, to avoid overfitting or underfitting. In all the experiments in this manuscript, this hyperparameter was fixed at 20.

Regarding $\lambda_{rec}$, as its value increased, clustering performance and representation quality first improved and then remained stable, while batch correction consistently decreased. The final overall score also initially increased and then stabilized (Supplementary Fig. 40c). We regarded the reconstruction loss as a crucial component for ensuring the validity of our autoencoder-based methods, and therefore, we recommend setting $\lambda_{rec}$ to no less than 10. In all the experiments in this manuscript, this hyperparameter was fixed at 10.

Subsequently, we conducted sensitivity analyses on other designed hyperparameters. To start with, we tested the impact of the number of training epochs for stage 1 and stage 2, increasing them from 0 to 1000 in 11 sets of experiments. For stage 1, which serves as the pretraining stage, we observed that as the number of training epochs increased, clustering performance initially improved and then stabilized, while representation quality and batch correction scores did not exhibit significant trends. The final overall score increased and then decreased, with the best performance observed between 400 and 600 epochs (Supplementary Fig. 41a). In all the experiments in this manuscript, the number of training epochs for stage 1 was fixed at 500. We believe that modifying this value is usually unnecessary, and it was set corresponding to the number of training epochs set in STAligner for the pretraining stage (stage 1).

For stage 2, which is essential for integration, we found that with an increasing number of training epochs, clustering performance improved and then stabilized, representation quality scores gradually decreased, and batch correction scores steadily increased. The final overall score first increased and then gradually declined (Supplementary Fig. 41b). We recommend setting this hyperparameter within the range of 300 to 600 epochs to avoid overfitting or underfitting. In all the experiments in this manuscript, this hyperparameter was fixed at 500.

During training, the pseudo-noise generation and calculation of latent loss involve selecting a certain number ($k$) of nearest neighbors from the target slice for each spot to achieve stochastic domain translation. We conducted 10 experiments varying $k$ from 10 to

100. We found that increasing $k$ had little effect on clustering performance, while representation quality and batch correction scores both increased initially and then stabilized, leading to a similar trend in the final overall score (Supplementary Fig. 41c). Therefore, we recommend setting this hyperparameter to no less than 40. However, if the goal is to identify spot-types unique to a particular slice, this hyperparameter should not be set too high. In all the experiments in this manuscript, this hyperparameter was fixed at 50.

Since the latent loss is in the form of triplet loss51, we implemented a clamp strategy with a margin hyperparameter ($m$). Triplets where the difference between the distance of a spot to its positive sample and the distance of that spot to its negative sample exceeds the $m$ are excluded from the backpropagation process. This strategy helps to remove positive samples that are too distant from the spot (potentially incorrect positives) or negative samples that are too close to the spot (potentially incorrect negatives). As we gradually increased $m$ (where a larger margin reduces the impact of this strategy), we observed a slight decrease in clustering performance, a general decline in representation quality, and an increase followed by stabilization in batch correction scores (Supplementary Fig. 41d). Based on the final overall scores, we suggest setting the margin within the range of 5 to 15. In all the experiments in this manuscript, this hyperparameter was fixed at 10.

Lastly, our method includes a hyperparameter, *min_cells_rate*, which is set according to the dataset. When filtering peaks, we only retain peaks that are accessible in at least a proportion of spots equal to *min_cells_rate*. We gradually varied this hyperparameter, resulting in different numbers of retained peaks (Supplementary Tab. 6), and conducted experiments. We found that as this hyperparameter increased, clustering performance and representation quality scores initially remained unchanged and then decreased, while batch correction scores increased. The final overall score first increased and then decreased (Supplementary Fig. 41e).

To ensure sufficient biological variation is retained and to avoid overfitting, we recommend keeping at least 30,000 peaks for data integration.
